# Endophyte genomes support greater metabolic gene cluster diversity compared with non-endophytes in *Trichoderma*

**DOI:** 10.1101/2023.03.14.532605

**Authors:** Kelsey Scott, Zachary Konkel, Emile Gluck-Thaler, Guillermo E. Valero David, Coralie Farinas Simmt, Django Grootmyers, Priscila Chaverri, Jason Slot

## Abstract

*Trichoderma* is a cosmopolitan genus with diverse lifestyles and nutritional modes, including mycotrophy, saprophytism, and endophytism. Previous research has reported greater metabolic gene repertoires in endophytic fungal species compared to closely-related non-endophytes. However, the extent of this ecological trend and its underlying mechanisms are unclear. Some endophytic fungi may also be mycotrophs and have one or more mycoparasitism mechanisms. Mycotrophic endophytes are prominent in certain genera like *Trichoderma*, therefore, the mechanisms that enable these fungi to colonize both living plants and fungi may be the result of expanded metabolic gene repertoires. Our objective was to determine what, if any, genomic features are overrepresented in endophytic fungi genomes in order to undercover the genomic underpinning of the fungal endophytic lifestyle. Here we compared metabolic gene cluster and mycoparasitism gene diversity across a dataset of thirty-eight *Trichoderma* genomes representing the full breadth of environmental *Trichoderma*’s diverse lifestyles and nutritional modes. We generated four new *Trichoderma endophyticum* genomes to improve the sampling of endophytic isolates from this genus. As predicted, endophytic *Trichoderma* genomes contained, on average, more total biosynthetic and degradative gene clusters than non-endophytic isolates, suggesting that the ability to create/modify a diversity of metabolites potential is beneficial or necessary to the endophytic fungi. Still, once the phylogenetic signal was taken in consideration, no particular class of metabolic gene cluster was independently associated with the *Trichoderma* endophytic lifestyle. Several mycoparasitism genes, but no chitinase genes, were associated with endophytic *Trichoderma* genomes. Most genomic differences between *Trichoderma* lifestyles and nutritional modes are difficult to disentangle from phylogenetic divergences among species, suggesting that Trichoderma genomes maybe particularly well-equipped for lifestyle plasticity. We also consider the role of endophytism in diversifying secondary metabolism after identifying the horizontal transfer of the ergot alkaloid gene cluster to *Trichoderma*.

## INTRODUCTION

*Trichoderma* (Hypocreales, Sordariomycetes, Ascomycota) is a cosmopolitan genus of fungi found growing on many different substrates which define their “lifestyle.” The majority of *Trichoderma* species are soil inhabitants. However, many *Trichoderma* species live as endophytes, i.e., growing asymptomatically beneath the epidermis of a living plant host [1–3], and some prefer decaying plant matter or other living fungi as a source of nutrition [3, 4]. A subset of *Trichoderma* species appears to be limited to a single lifestyle, while other species have been recorded as having multiple lifestyle strategies [3–6]. Within these lifestyles, *Trichoderma* species usually have either a mycotrophic or saprophytic nutritional mode [4].

Multiple *Trichoderma* species, including endophytic species, are known for their ability to produce a variety of secondary metabolites and/or exhibit one or more mycoparasitic mechanisms [1,7–15]. Some *Trichoderma* secondary metabolites mimic phytohormones, which impact host plant vigor and trigger plant-wide defenses [16–21]. *Trichoderma* species also produce secondary metabolites that can antagonize invading microorganisms to the benefit of themselves and their host plants [1,22,23]. Furthermore, endophytism is expected to select for fungal metabolic pathways that provide resistance to defensive metabolites, such as flavonoids, terpenes, and phenolics, which plants produce constitutively or in response to damage or invasion [24–26]. Specifically, the ability to degrade or modify diverse phenolics might be favored in endophytic *Trichoderma* species [27–31]. The mechanisms that enable these fungi to colonize both living plants and fungi may be the result of expanded metabolic gene repertoires. Therefore, we expect that endophytic *Trichoderma* species would have an expanded metabolic repertoire compared to non-endophytic species.

Fungal metabolic pathways are often encoded in metabolic gene clusters, loci with closely-linked genes for enzymes, transporters, and regulation of a common pathway [32–36]. Metabolic gene clusters can encode the production of secondary metabolites (biosynthetic gene cluster, BGC) or their degradation (degradative gene clusters, DGC), among other functions. The clustering of metabolic gene clusters is evidence of selection on the metabolic function they perform. Therefore, it can suggest more precise hypotheses about the genetic basis of species ecology than genes in isolation [37, 38]. Fungal BGCs can encode the production of ecologically significant metabolites, such as the antibiotics trichodermin and gliotoxin [39, 40], and anti-herbivory ergot alkaloids produced by Clavicipitaceae species [27]. By contrast, DGCs may provide fitness benefits to plant-associated fungi by degrading exogenous toxins, such as the fungicide cyanate [41] or plant defense compounds, such as benzoxazolinones and stilbenes [42, 43].

Mycoparasitic fungi produce chitinases, serine proteases, glucanases, other cell-wall degrading enzymes, and antibiotics/antifungals, all of which have been associated with mycoparasitism [44–46]. Chitinase genes are necessary not only for fungal growth and development but also for the parasitism of competing organisms, such as insects, nematodes, and other fungi [47, 48]. Similar to the ecological patterns seen in certain gene clusters, chitinase genes have been identified as being expanded in mycoparasitic fungal lineages and reduced in non-mycoparasitic lineages, likely due to adaptive natural selection [49, 50]. In *Trichoderma*, chitinase genes in the BI/BII subgroup were reported to be expanded in mycoparasitic species, exhibiting positive selection [51]. Similar to BGCs and DGCs, an expanded repertoire of chitinases and other mycoparasitism-related genes may provide fitness benefits to plant-related fungi by facilitating competition with organisms competing with the fungal endophyte or the endophyte’s host plant.

The diversity of genes and gene clusters is reportedly associated with particular fungal lifestyles [26, 50]. For example, endophytic Xylariales species contain a greater diversity of metabolic gene clusters when compared to closely-related non-endophytic species [26]. Similarly, chitinase genes in *Trichoderma* are more diverse in mycoparasites than in saprophytes [50]. Many fungal species are not constrained to a single lifestyle and instead have the capability to interact with a variety of other organisms and substrates, a trait which is potentially explained by their expanded chitinolytic and metabolic diversity [52–54]. Specific fungal gene clusters are reported to be enriched in particular ecological niches [38] possibly due to horizontal gene transfer (HGT) among species with overlapping lifestyles and thus shared selective pressures [30,35,55,56]. Differential metabolic gene cluster diversification [57–59] and loss among fungi in different ecological niches may also contribute to uneven cluster distributions and diversity [32,35,52].

In this study, we asked whether endophytic *Trichoderma* genomes have a greater diversity of BGCs and DGCs than non-endophytic *Trichoderma* genomes. We also sought to determine whether the genomes of endophytic *Trichoderma* have more diverse repertoires of chitinase and other mycoparasitism-related genes compared to non-endophytic *Trichoderma* genomes. To address these questions, we sequenced the genomes of four Peruvian isolates of a recently described endophytic species *T. endophyticum*, which is placed in the *T. harzianum* species complex and found as both an endophyte of several species of Neotropical trees [3] as well as living freely in soil [6]. These new genomes were included in a robust comparative database with 34 reannotated publicly available *Trichoderma* genomes that exhibited various lifestyles and nutritional modes to identify ecological patterns. We specifically investigated trends in the composition of BGCs, DGCs, chitinase genes, and other mycoparasitism-associated genes across the 38 genomes.

We found that endophytic *Trichoderma* species have a significantly greater overall diversity of both DGC and BGC than non-endophytic *Trichoderma* species. Still, we found limited evidence that any particular gene cluster families or classes contributed to this overall trend. We did not find a significant association between chitinase diversity and the endophytic lifestyle, though we identified several other mycoparasitism genes that are expanded in endophytic *Trichoderma* genomes. Despite identifying several overarching trends in the genomic features present in endophytic *Trichoderma* species, the interrelatedness between *Trichoderma* lifestyle and phylogeny makes it difficult to infer specific genomic features that could underpin the endophytic lifestyle. Additionally, we identified the horizontal transfer of the ergot alkaloid gene cluster to *Trichoderma*.

## RESULTS

### Four new *T. endophyticum* genomes supplement the *Trichoderma* dataset with high-quality endophytic genomes

Four endophytic isolates of *T. endophyticum* were previously isolated from rubber trees (*Hevea* spp.) in Peru [3]. Isolates PP24 and PP89 were obtained from the trunk of wild *Hevea guianensis* and isolates LA10 and LA29 were obtained from the trunk of wild *Hevea brasiliensis*. All four isolates were sequenced with both Illumina and Nanopore sequencing technology and *de novo* assembled for this study. Twenty-nine additional *Trichoderma* assemblies were downloaded directly from NCBI, and six genomes were assembled using publicly available SRA data (Table S1). The four *T. endophyticum* genomes each have a genome completeness of ≥99.0%, referencing the Sordariomycetes BUSCO dataset (sordariomyceta_odb9) (Table 1). The remaining *Trichoderma* genomes exhibit completeness ranging from 61.2-99.0% (Table S2). Due to a severely fragmented and incomplete genome (single and complete BUSCOs = 61.2%), *T.* cf*. atroviride* isolate LU140 was included in the phylogenomic tree (Fig. S1) but excluded from all subsequent analyses, resulting in a total of 38 high quality genomes in the final dataset. The four *T. endophyticum* genomes ranged in length between 38.9-39.2Mb (average 39.0 Mb), and the overall full *Trichoderma* genomes in the dataset ranged from 31.7-45.8 Mb (average 37.1 Mb). The *T. endophyticum* genomes contained between 12,606 and 12,635 genes, and the remaining genomes in the dataset ranged from 9,242 and 14,297 genes (Table 1 and Table S2). The *Trichoderma* clade with the lowest average number of genes per genome was Longibrachiatum (average of 9776 genes per genome), and the clade with the highest was Virens (average of 13,231 genes per genome). The *T. endophyticum* assemblies are available from NCBI under the following BioProject Accession: PRJNA899549.

**Table 1.**
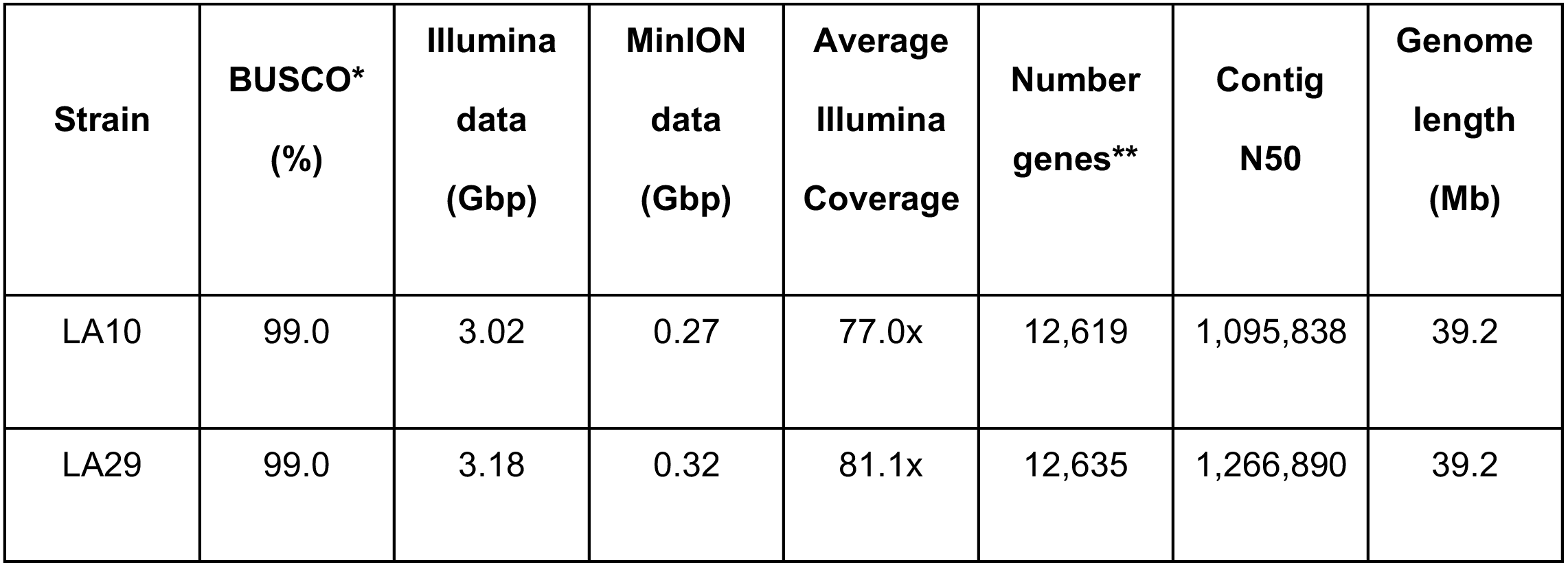
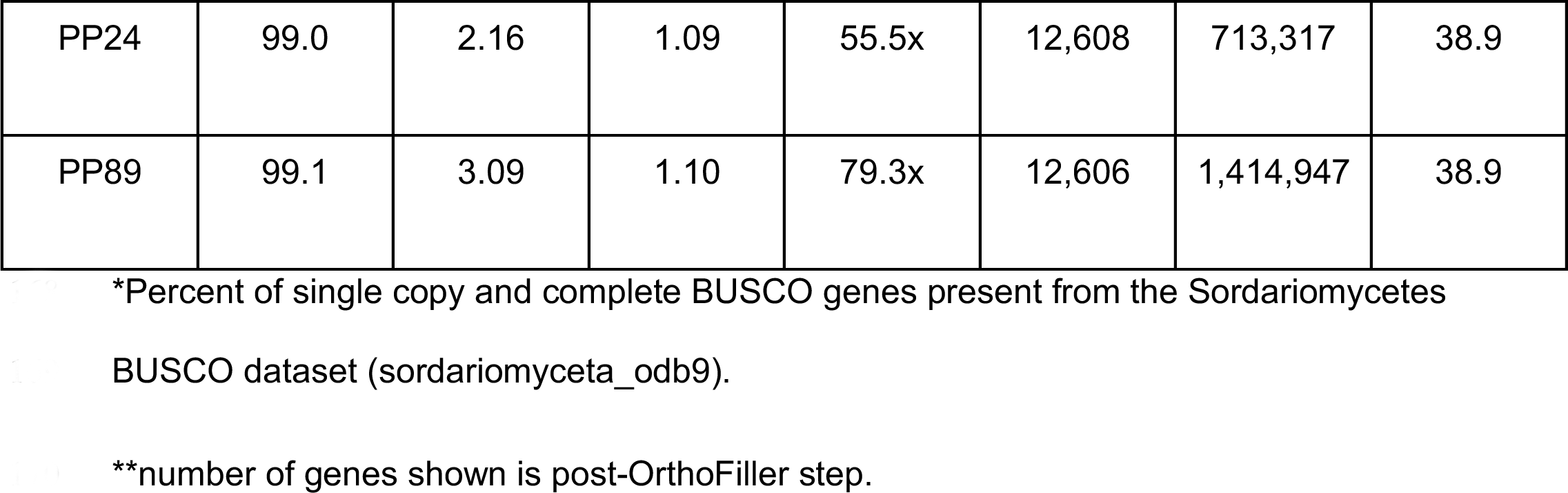
Genome statistics of each of the hybrid-assembly T. endophyticum genomes.

### Phylogenomic analysis of thirty-eight *Trichoderma* isolates from diverse lifestyles and nutritional modes yields a tree concordant with previous multi-gene phylogenies

We reconstructed a phylogenomic tree of 154 well-supported single-copy orthologs (SCO) (average ≥75% bootstrap percent support [BP] at each node) (Fig. S1). This phylogeny resolves into five distinct clades (100% BP), which are in agreement with previously defined groups in *Trichoderma* [3,4,60,61]. The resolved clades are identified as Harzianum, Virens, Longibrachiatum, section *Trichoderma*, and Brevicompactum. Harzianum contains *T. endophyticum* (4 isolates), *T. harzianum* (2), *T. simmonsii* (1), *T. guizhouense* (1), an unspecified *Trichoderma* species (OTPB3; 1 isolate), *T. afroharzianum* (2), and *T. plueroticola* (1). Virens contains three *T. virens* isolates. Longibrachiatum consists of *T. reesei* (5), *T. parareesei* (1), *T. longibrachiatum* (2), *T. bissettii* (1), and *T. citrinoviride* (1). Section *Trichoderma* contains *T. atroviride* (4), *T.* cf*. atroviride* (2), *T. gamsii* (2), *T. koningiopsis* (1), *T. asperellum* (2), and *T. hamatum* (1). Brevicompactum contains *T. brevicompactum* (1) and *T. arundinaceum* (1).

Harzianum contains the majority of the endophytic isolates in this dataset and is composed entirely of isolates with a mycotrophic nutritional mode. All four *T. endophyticum* genomes are located in a single subclade that is well-supported (100% BP). Furthermore, these genomes resolved into separate groups within the *T. endophyticum* species subclade based on their geographical origin/host tree species (PP vs. LA). The *Trichoderma* isolate with an undescribed species (OPTB3), previously considered to be *T. harzianum*, is most closely related to *T. guizhouense* (NJAU4742). The clade Virens is most closely related to Harzianum, and these clades are well-defined (100% BP).

Virens contains only the three *T. virens* isolates, which all have a mycotrophic nutritional mode and have been reported to live on other fungi, in the soil, and in wood and/or leaf litter.

Longibrachiatum is an entirely saprotrophic clade that lacks any fungi-dwelling or endophytic isolates and is the only clade that does not contain species with more than one reported lifestyle. This clade is well-supported (100% BP) and mostly composed of the five monophyletic (100% BP) *T. reesei* isolates. *Trichoderma bissettii* (JCM 1883) and the two *T. longibrachiatum* genomes (ATC 18648 and SMF2) make up the sister clade to the *T. reesei* genomes (100% BP), and *T. citrinoviride* (TUCIM 6016) diverged earliest in this Longibrachiatum clade.

Section *Trichoderma* is the only clade that contains both saprotrophic and mycotrophic species. All species in this clade can be found living freely in the soil, although most species are also reported to live as either endophytes or in wood/leaf litter. Most of this clade is composed of a subclade (100% BP) containing the four *T. atroviride* and two *T.* cf*. atroviride* isolates, all of which are saprotrophic and can live in both the soil and in wood/leaf litter. The two *T.* cf*. atroviride* isolates (LU140 and LU132) are clustered within this subclade with high support values (100% BP). *Trichoderma koningiopsis* (POS7) is located adjacent to the two *T. gamsii* genomes (A5MH and T6085). Another subclade that is marginally supported in section *Trichoderma* (81% BP) contains the saprotrophic *T. hamatum* (GD12) and two *T. asperellum* genomes (CBS 433.97 and B05). Most endophytic species in this dataset are reported to have a mycoparasitic lifestyle, while *T. hamatum* is one of the two endophytic *Trichoderma* species with a saprotrophic lifestyle.

Brevicompactum is a well-supported clade (100% BP) adjacent to section *Trichoderma* and contains only saprotrophic species. Both species that compose this clade are capable of living freely in the soil, although only *T. brevicompactum* is also found as an endophyte. Additionally, *T. brevicompactum* is the second of the two species in our dataset that is both saprotrophic and endophytic.

### Endophytic isolates have a greater number of DGCs and BGCs compared with non-endophytic isolates

*Trichoderma* genomes contain multiple DGCs, which we have grouped into evolutionarily-related cluster families and gene cluster classes defined by which common “core” processing functions are present in each cluster. We identified DGCs containing salicylate hydroxylase (SAH) (104 total cluster families identified in the dataset), quinate 5-dehydrogenase (QDH) (188 total), aromatic ring-opening dioxygenase (ARD) (53 total), naringenin 3-dioxygenase (NAD) (64 total), ferulic acid decarboxylase (FAD) (25 total), phenol 2-monooxygenase (PMO) (45 total), catechol dioxygenase (CCH) (71 total), benzoate 4-monooxygenase (BPH) (five total), pterocarpan hydroxylase (PAH) (two total). Additionally, we found multiple “hybrid” clusters that contain several different core genes (e.g., hybrid CCH-PMO cluster). On average, a *Trichoderma* genome contains 35.4 DGCs (range 24-56 clusters). One or more gene clusters containing SAH, CCH, QDH, PMO, or CCH-SAH are present in most *Trichoderma* genomes (Fig. 1). PAH clusters are only found in each of the two *T. asperellum* genomes, with a single DGC per genome. BPH clusters are found only in *T. harzianum* (T6776) and *T. gamsii* (T6085) genomes. SAH-SDO clusters are present only in *T. arundinaceum* (IBT-40837) and *T. brevicompactum* (IBT-40841). Despite varying assembly quality throughout the dataset, there is no correlation between the number of identified DGCs and assembly N50 (Fig. S2A). The number of identified BGCs per genome positively correlates with the number of identified DGCs per genome (Fig. S3A). The proportion of BGC to the total number of genes within a genome also positively correlates with the proportion of DGC to the total number of genes (Fig. S3B). Larger *Trichoderma* genomes have a greater diversity of DGCs (Fig. S4A) as well as higher proportions of DGCs to total number of genes (Fig.S5A). *Trichoderma* genomes that contain greater numbers of genes also contain greater numbers of DGCs (Fig. S6).

**Fig 1.**
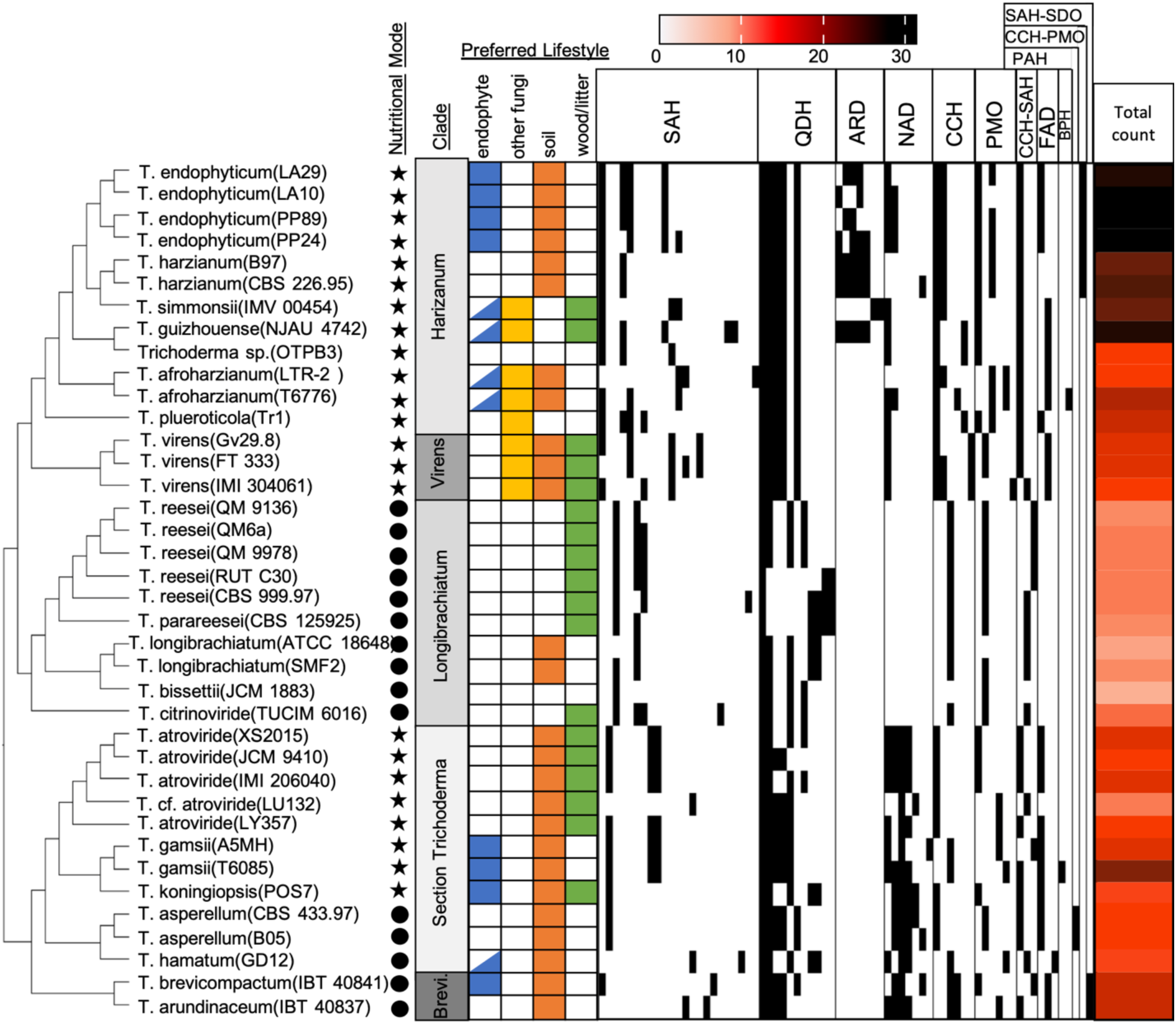
Distribution and total amount of DGC families per genome varies across isolates and *Trichoderma* clade. The presence of different DGC families in a particular genome are indicated by a black cell, absence of a particular family is indicated by a white cell. DGC families are separated by cluster class. *Trichoderma* species lifestyle is indicated by presence/absence of color-coordinated cell, nutritional mode is indicated by either a circle (saprotroph) or a star (mycotroph). *Trichoderma* species lifestyle is indicated by the presence or absence of a color-coordinated cell, a potential endophyte lifestyle is indicated by a half-filled cell. Nutritional mode is indicated by either a circle (saprotroph) or a star (mycotroph). The total amount of DGC families identified in each genome is indicated by the red heatmap. Clade “Brevicompactum” is abbreviated as “Brevi”.

*Trichoderma* endophytes have a significantly greater number of total DGCs compared to non-endophytes (Fig. S7A). Additionally, we compared the number of DGCs within each DGC class (defined by the cluster core gene) between endophyte genomes and non-endophyte genomes. Endophytic genomes contained greater numbers of ARD, NAD, and FAD DGCs (Fig. S8).

Results from Blomberg’s K and Pagel’s Lambda analyses suggest that the individual DGC class counts per genome as well as the overall DGC count per genome is consistent with the *Trichoderma* phylogeny (Blomberg’s K and Pagel’s Lambda: p ≤ 0.05) (Fig. S9A, B). According to Blomberg’s K analysis, the DGC classes CCH-PMO, ARD, and FAD are overdispersed in the *Trichoderma* phylogeny (K < 0.8) as well as being overrepresented in endophytic genomes (average count in endophyte genome:average count in non-endophyte genomes [E:NE] > 2). However, their counts and distributions are also consistent with Brownian motion evolution (Blomberg’s K p < 0.05). All other DGC classes, as well as the total number of DGCs per genome, are either not enriched in endophyte genomes (E:NE < 2) and/or are consistent with Brownian motion model evolution (Blomberg’s K/Pagel’s Lambda: p ≤ 0.05) (Fig. S9A,B).

Similarly, the calculation of Fritz and Purvis’ D-statistic for each individual DGC cluster family suggests that the distribution of most DGC families are consistent with Brownian motion evolution (D-statistic < 1, p > 0.05) (Fig. S10). Although CCH-PMO hybrid family 187 was found enriched in endophytes (E:NE > 2) and was found to be a DGC family enriched in a particular lineage (D < 0), the Fritz and Purvis’ analysis suggests that the distribution of this DGC family is consistent with the *Trichoderma* phylogeny (p > 0.05). The CCH-PMO family is only found in the four *T. endophyticum* genomes and the two *T. harzianum* genomes. NAD family 127 is overdispersed (D > 1) in the *Trichoderma* phylogeny and its distribution is not consistent with Brownian motion (p ≤ 0.05), however, this DGC family is not overrepresented in endophytic genomes (E:NE < 2). All other DGC families either are not enriched in endophytic genomes, have a distribution consistent with Brownian motion, or are not overdispersed in the *Trichoderma* phylogeny.

The distribution of DGC cluster families varies greatly among the *Trichoderma* dataset and primarily reflects the *Trichoderma* phylogeny (Fig. 1). Longibrachiatum has the fewest overall number of identified DGCs (Fig. S11A), as well as one of the lowest proportions of DGCs to total gene number (Fig. S11C). All Longibrachiatum genomes also lack FAD or NAD DGCs, despite these cluster classes being present in the other four clades (Fig. 1). Brevicompactum and Harzianum have the highest number of DGCs and proportion of DGCs to total gene number (Fig. S11A,C). Compared to Harzianum and Virens, there is greater NAD diversity within Brevicompactum and section *Trichoderma* (Fig. 1). However, there are no NAD DGCs in two genomes in Harzianum, despite all other Harzianum genomes containing at least one NAD DGC. Similarly, there are no FAD clusters in two Harzianum genomes and one section *Trichoderma* genome, despite this DGC class being identified in all other genomes in this analysis besides Longibrachiatum genomes. At least one ARD family is present in all the Harzianum clade genomes except the unknown *Trichoderma* species (OTPB3).

The *T. endophyticum* genomes are among the genomes with the highest diversity of DGCs (Fig. 1) and have a greater diversity of DGCs clusters than expected, given the association between total gene number and diversity of DGCs (Fig. S10). Each *T. endophyticum* genome contains at least two ARD DGCs and a CCH-PMO DGC, both exclusively identified in Harzianum. In each *T. endophyticum* genome, there is a least one NAD DGC also found in most other genomes, except for genomes in Longibrachiatum. The four *T. endophyticum* genomes have almost identical DGC repertoires, with minimal differences between their overall number of DGCs and individual DGC families between the *T. endophyticum* genomes (Fig. 1).

Every *Trichoderma* genome contains multiple BGCs, which we have grouped into families (homologous clusters across species), and further categorized into classes based on the “core” metabolic genes present in each cluster, as assigned by antiSMASH: nonribosomal peptide synthetase (NRPS) (384 total clusters identified in the dataset), type I polyketide synthase (T1PKS) (496 total), NRPS-like (309 total), terpene (304 total), T1PKS-NRPS hybrid clusters (240 total), fungal-associated ribosomally synthesized and posttranslationally modified peptides (fungal-RiPPs) (nine total), beta-lactone (12 total), and other “hybrid” clusters containing two or more of these key genes (51 total). BGCs containing fungal-RiPPs and beta-lactone are found in exclusively in Harzianum genomes (Fig. 2, Fig. S12). Additionally, a single indole BGC was identified in the *T. brevicompactum* (IBT40841) genome (not shown in Fig. 2). Similar to the diversity of DGCs in the dataset, there are correlations between different genome features and the number of BGCs in each *Trichoderma* genome. There is no correlation between genome N50 and BGCs identified per genome, or metabolic genes identified per genome (Fig. S2B,C). Larger *Trichoderma* genomes tend to have greater diversities of BGCs and metabolic genes (Fig. S4B, C), as well as a greater proportion of BGCs to total genes (Fig. S5B). Additionally, genomes with greater numbers of genes tended to have a greater diversity of BGCs (Fig. S6B).

**Fig 2.**
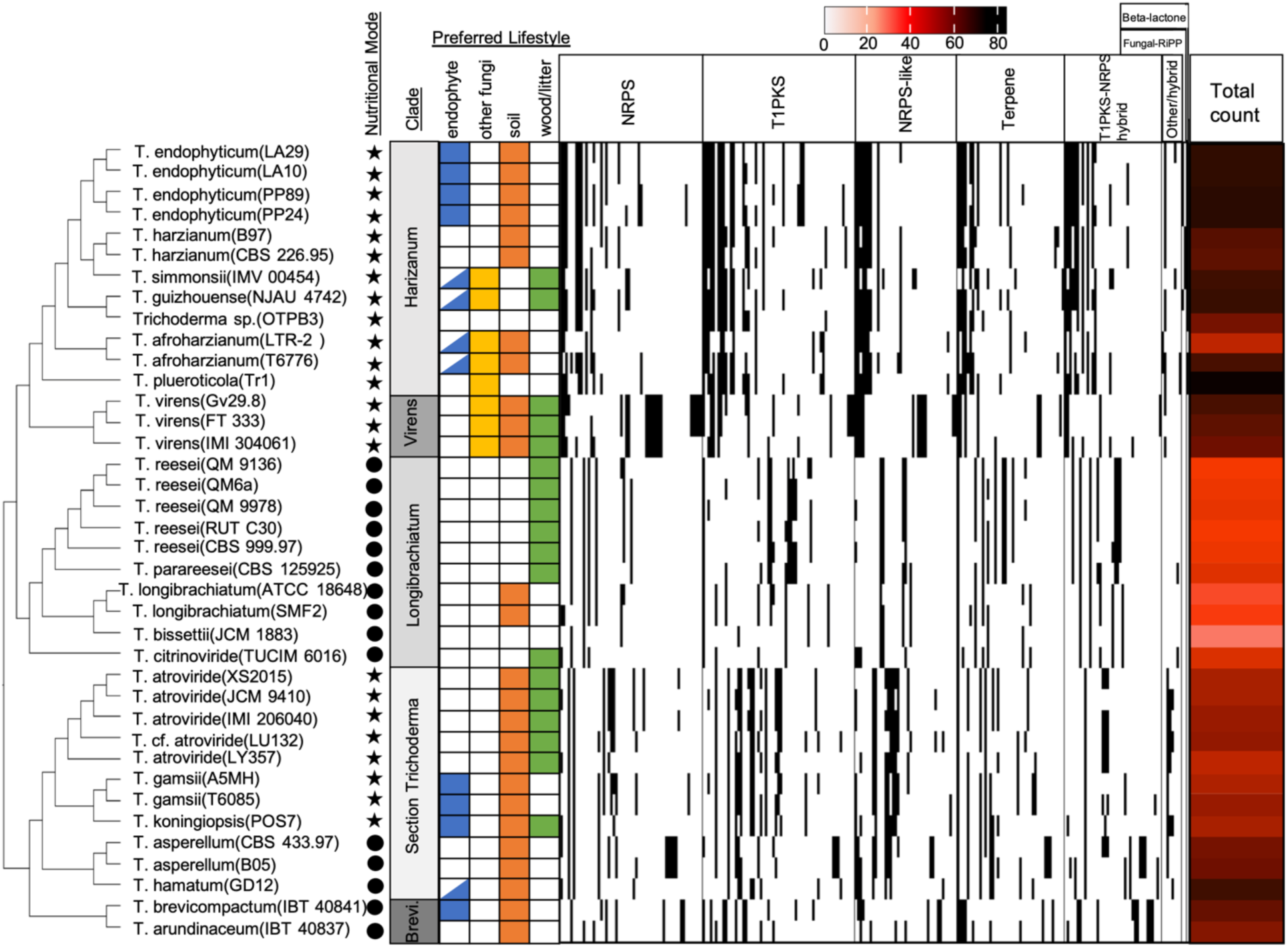
Distribution and total amount of BGC families per genome varies across isolates and *Trichoderma* clade. The presence of different BGC families are indicated by a black cell, BGC families are separated by class. The nutritional mode of each species is indicated by either a circle (saprotroph) or a star (mycotroph). BGC families present only in a single genome were removed for visualization purposes. *Trichoderma* species lifestyle is indicated by the presence or absence of a color-coordinated cell. Nutritional mode is indicated by either a circle (saprotroph) or a star (mycotroph). Endophyte cells half-filled indicated a potential endophytic lifestyle for that species. The total amount of DGC families identified in each genome are indicated by the red heatmap. Clade “Brevicompactum” is abbreviated as “Brevi”.

Similar to the pattern found in *Trichoderma* DGCs, *Trichoderma* endophyte genomes have a significantly greater number of BGCs when compared to non-endophyte genomes (Fig. S7B). Endophytic genomes were found to have significantly greater numbers of T1PKS, NRPS-like, beta-lactone, and fungal-RiPP BGCs (Fig. S13). Phylogenetic correction and assessment of the BGC class counts with Blomberg’s K and Pagel’s Lambda suggest that the counts of each separate BGC classes are consistent with the Brownian motion model of trait evolution (Blomberg’s K and Pagel’s Lambda > 0.8, p ≤ 0.05) (Fig. S9, A.B). Beta-lactone BGCs are found in greater abundance in endophytic genomes (E:NE > 2) and appear to be overdispersed in the *Trichoderma* genus (K < 0.8); however, the distribution of the beta-lactone is ultimately consistent with Brownian model trait evolution (Blomberg’s K: p ≤ 0.05). All other BGC classes we evaluated, as well as the total count of BGCs per genome, despite being overdispersed in the phylogeny (Blomberg’s K/Pagel’s Lambda < 0.08), are not overrepresented in endophytic genomes (E:NE < 2) and have a distribution consistent with the *Trichoderma* phylogeny (Blomberg’s K/Pagel’s Lambda: p ≤ 0.05) (Fig. S9A,B).

Similarly, the calculation of Fritz and Purvis’ D-statistic for each individual BGC family suggests that the distribution of each BGC family can also be explained by the *Trichoderma* phylogeny (D-statistic < 1, p > 0.05) (Fig. S14). Terpene family 1276 has a very high D-statistic (D > 4.5), indicating it is overdispersed in *Trichoderma*; however, it is not found overrepresented in endophytic genomes (E:NE < 2), and its distribution was consistent with Brownian motion (p > 0.05). Terpene family 1276 is found in all *Trichoderma* genomes besides *T. simmonsii* (IMV 00454). All other BGC families either are not enriched in endophytic genomes (E:NE < 2), have a distribution consistent with Brownian motion (D-statistic: p > 0.05), or were not overdispersed in the *Trichoderma* phylogeny (D-statistic < 1).

The distribution of BGC cluster families varies greatly across the *Trichoderma* dataset and primarily reflect the *Trichoderma* phylogeny (Fig. 2). There are significant differences in the total diversity of BGCs identified across the different clades (Fig. S11B), as well as the proportion of BGCs to total gene number (Fig. S11D). Longibrachiatum genomes contain the lowest diversity of BGCs, while Harzianum, Brevicompactum, and Virens contain the highest diversity of BGCs (Fig. S11B). The same pattern of BGC diversity is also observed in the proportion of BGCs to total gene number; Longibrachiatum genomes have the lowest proportions, while Harzianum, Brevicompactum, and Virens have the highest proportions of BGCs to total identified genes (Fig. S11D).

The *T. endophyticum* genomes have some of the highest diversities of BGCs (Fig. 2) and have a greater diversity of BGC clusters than expected, given the association between BCG diversity and genome length (Fig. S4B) and total gene number (Fig. S6B). The *T. endophyticum* genomes contain a similar number of NRPS (12-13), T1PKS (16-20), NRPS-like (8-9), T1PKS-NRPS (9-10), fungal RiPP (1), and beta-lactone (1) DGCs. All *T. endophyticum* genomes contain a terpene-NRPS (family 1020) hybrid cluster, which is also present in all other Harzianum genomes as well as *T. virens* (IMI 304061). Additionally, *T. endophyticum* LA29 contains an NRPS-indole (family 1016) hybrid cluster, only found in LA29 and *T. virens* (IMV 00454). Overall, the *T. endophyticum* genomes are very similar and have few differences between their total number of BGCs and individual BGC families.

### Several mycoparasitism genes are associated with the *Trichoderma* endophytic lifestyle

The *Trichoderma* genomes differ in their diversity and distribution of the 746 genes previously linked to fungal mycoparasitism [62–64]. Of the 746 mycoparasitism genes, 468 of these genes are present (at least one ortholog) in all 38 *Trichoderma* genomes, and some mycoparasitism genes have more than one ortholog present in a single genome (Suppl. Data file 1). The number of total mycoparasitism genes per genome ranges from 676 (*T. longibrachiatum* ATCC18648) to 991 (*T. virens* IMI304061) and on average, a *Trichoderma* genome contains 850.4 mycoparasitism genes. The total amount of genes per genome positively correlated with the amount of identified mycoparasitism genes per genome (Fig. S15A). Larger genomes tend to have greater numbers of mycoparasitism genes (Fig. S15B) and genome quality (N50) does not correlate with the number of mycoparasitism genes identified per genome (Fig. S15C).

The total diversity of mycoparasitism genes is significantly higher in endophyte genomes compared to non-endophyte genomes (Fig. S7D), and mycotroph genomes have greater amounts than saprotroph genomes (Fig. S16D). However, when phylogenetic signal was controlled for, the distribution and diversity of the total number of mycoparasitism genes as well the distribution of most individual mycoparasitism genes can be explained by *Trichoderma* phylogeny, as determined by Blomberg’s K and Pagel’s Lambda analyses (Fig. 3A, B). There are three total orthogroups whose distribution is not consistent with Brownian motion evolution according to the two separate phylogenetic signal metrics and are also found to be overrepresented in endophyte genomes (Table 2). The two mycoparasitism genes orthogroups of interest from the Blomberg’s K calculations are OG0009104 (function: unknown) and OG0008134 (function: unknown). The singular mycoparasitism gene of interest from the calculation of Pagel’s Lambda is OG0009627 (function: unknown). From this same set of calculations, OG0008500 (function: unknown) was nearly considered an orthogroup of interest but was discarded due to its E:NE value (E:NE < 2). Besides the three genes discussed above, the remaining genes are not overrepresented in *Trichoderma* and have distributions consistent with Brownian motion (K > 0.8 or Lambda > 0.8, p ≤ 0.05). Most of the mycoparasitism genes enriched in endophyte genomes do not have a known eggnog functional annotation. The genes found in the highest ratio in mycotroph genomes vs. saprotroph genomes, as well as genes only found in mycotroph genomes, are shown in Table S4 (full results in Fig. S17). Similar to the endophyte results, most mycoparasitism genes enriched in mycotroph genomes or found only in mycotroph genomes had an eggnog functional annotation of “unknown”. Mycoparasitism orthogroup OG16824 which does have an available functional annotation (function: Post-translational modification, protein turnover, and chaperones) was found only in mycotroph genomes and did not have a distribution consistent with Brownian motion (Table S4).

**Fig 3.**
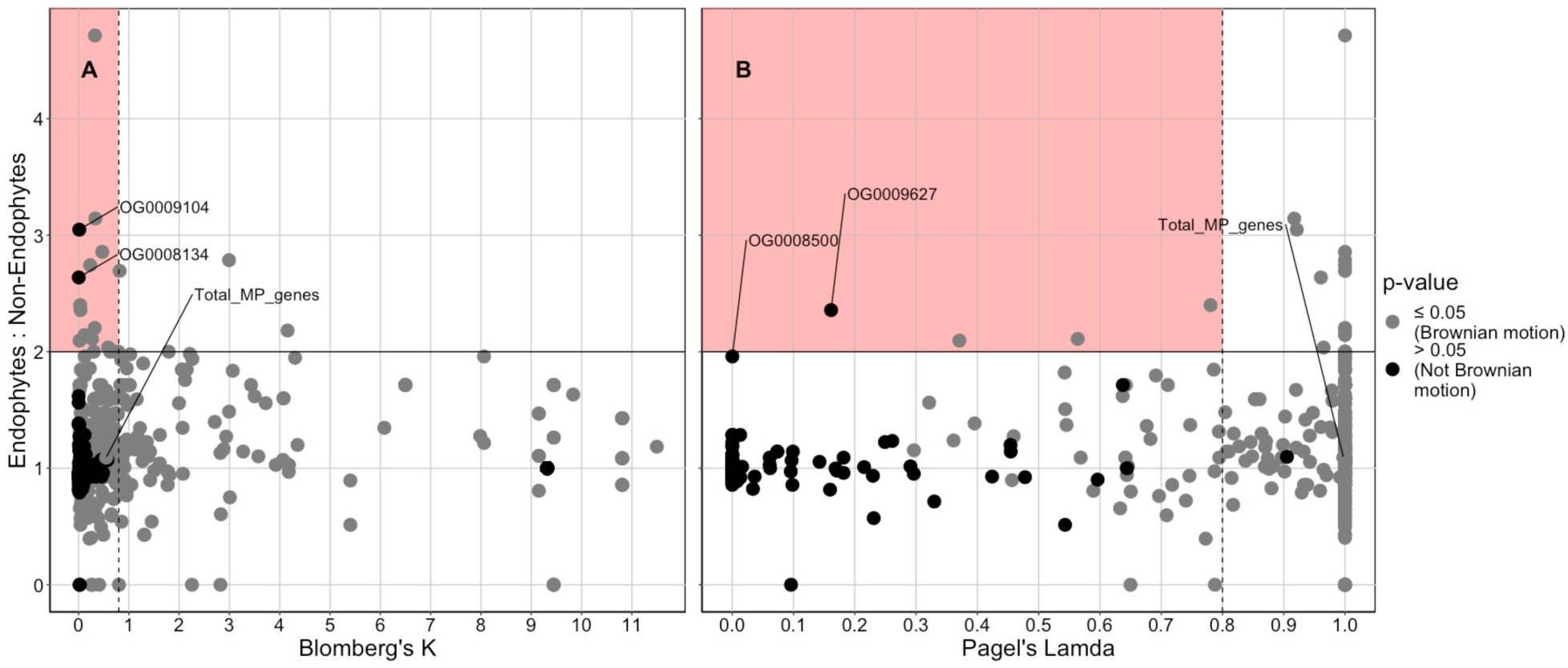
Several mycoparasitism genes potentially related to the *Trichoderma* endophytic lifestyle were identified. The phylogenetic and ecological (endophyte:non-endophyte ratio) signal of the distribution of the different mycoparasitism genes were determined by two methods. (A) Calculation of Blomberg’s K for each mycoparasitism gene distribution suggests that only two mycoparasitism genes are both overdispersed in the *Trichoderma* phylogeny (K < 0.8) and is not consistent with Brownian motion evolution (p > 0.05), while also being overrepresented in endophyte genomes (E:NE > 2). (B) Calculation of Pagel’s Lambda for each mycoparasitism gene distribution suggests that only a single mycoparasitism gene is both overdispersed in the *Trichoderma* phylogeny (Lambda < 0.8) and is not consistent with Brownian motion evolution (p > 0.05), while also being overrepresented in endophyte genomes (E:NE > 2).

**Table 2.**
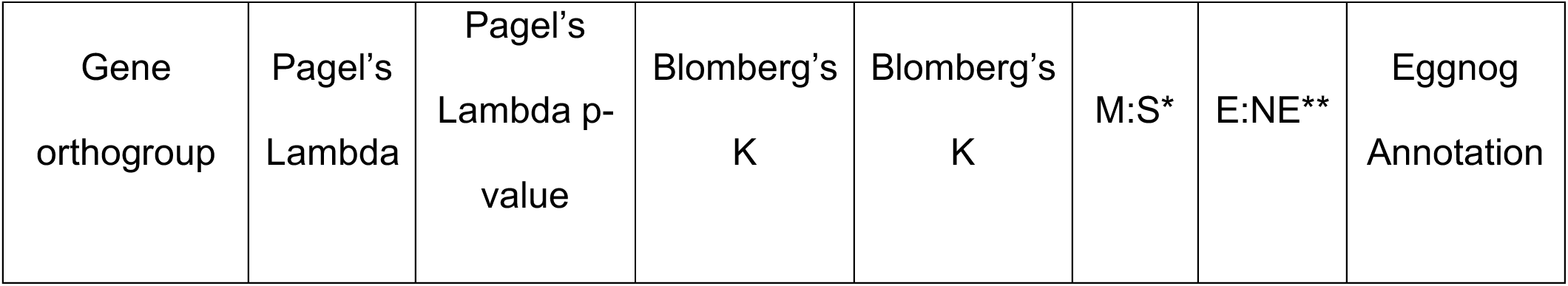
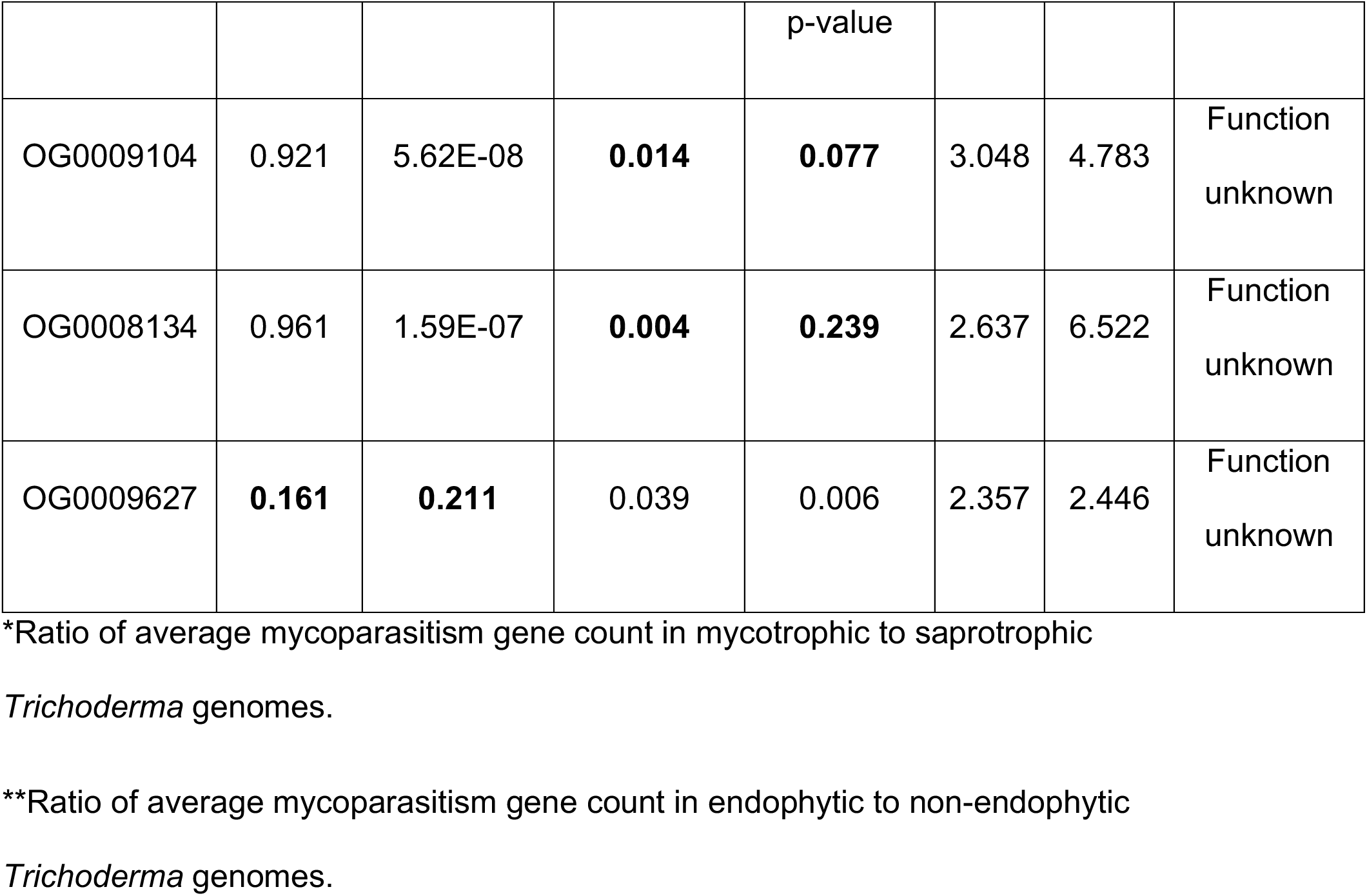
Mycoparasitism gene orthogroups of interest that are found in greater numbers in endophytic Trichoderma genomes.

|  |  |  |  | p-value |  |  |  |
| --- | --- | --- | --- | --- | --- | --- | --- |
| OG0009104 | 0.921 | 5.62E-08 | **0.014** | **0.077** | 3.048 | 4.783 | Function unknown |
| OG0008134 | 0.961 | 1.59E-07 | **0.004** | **0.239** | 2.637 | 6.522 | Function unknown |
| OG0009627 | **0.161** | **0.211** | 0.039 | 0.006 | 2.357 | 2.446 | Function unknown |

\*Ratio of average mycoparasitism gene count in mycotrophic to saprotrophic
*Trichoderma* genomes.
\*\*Ratio of average mycoparasitism gene count in endophytic to non-endophytic

Based on the total number of mycoparasitism genes per genome, the distribution of mycoparasitism genes varies greatly across the *Trichoderma* genus (Fig. S18). Longibrachiatum has the fewest of our targeted mycoparasitism-related genes, while Virens, section *Trichoderma*, and Harzianum have the most (Fig. 18A). However, Brevicompactum and section *Trichoderma* have the highest percent of their genomes composed of mycoparasitism genes (Fig. S18B). We ordinated the mycoparasitism gene count data after correcting for the phylogenetic signal inherently present the *Trichoderma* dataset using phylogenetic principal components analysis (pPCA), the results of which suggest that the number and distribution of the mycoparasitism genes primarily reflect *Trichoderma* phylogeny (Fig. S19A, S20). The differences in mycoparasitism gene distributions between the *Trichoderma* clades can clearly be seen in a visualization of the global positive and negative principal component values calculated from the pPCA (Fig. S19A).

The *T. endophyticum* isolates have almost identical distribution and diversity of mycoparasitism genes compared to each other, with the total count ranging from 891 to 901. The average number of *T. endophyticum* mycoparasitism genes (897.0 genes) was comparable to the combined average in all other endophyte genomes (898.6 genes) and below the combined average in all other mycotroph genomes (910 genes). The phylogenetically adjusted PCA (pPCA) groups *T. endophyticum* genomes with their relatives in Harzianum (Fig. S20). No mycoparasitism genes were exclusively present or absent in the *T. endophyticum* genomes.

### Chitinase gene repertoires reflect vertical inheritance and nutritional mode, but not lifestyle in *Trichoderma*

Chitinases are a complex group of carbohydrate-active enzymes important for mycoparasitism, so we profiled chitinase repertoires across all genomes separate from the mycoparasitism genes analysis. The chitinase genes in our dataset are grouped into nine chitinase classes as delimited in [65], including classes AII (76), AIV (79), AV (107), BI (215), BII (105), BV (77), CI (107), CII (194), and ChitD (38), which collectively bear 153 lysin domains (LysM) and 374 chitin-binding domains (CBD) (Fig. 4). Each *Trichoderma* genome contains two AII chitinases and one ChitD chitinase gene each. We found no class AIII, BIII, or BIV chitinase genes in any *Trichoderma* genomes. The distribution of chitinase genes and overall chitinase diversity varies greatly over our dataset and reflects the *Trichoderma* phylogeny (Fig.4). Chitinase diversity (total number of chitinase genes) within a genome ranges from 18 to 40 genes. Larger genomes tend to have a greater diversity of chitinase genes as well as a greater proportion of chitinase genes compared to total gene counts (Fig. S21A,B). There was no correlation between genome quality (N50) and the total number of chitinase genes identified (Fig. S21C).

**Fig 4.**
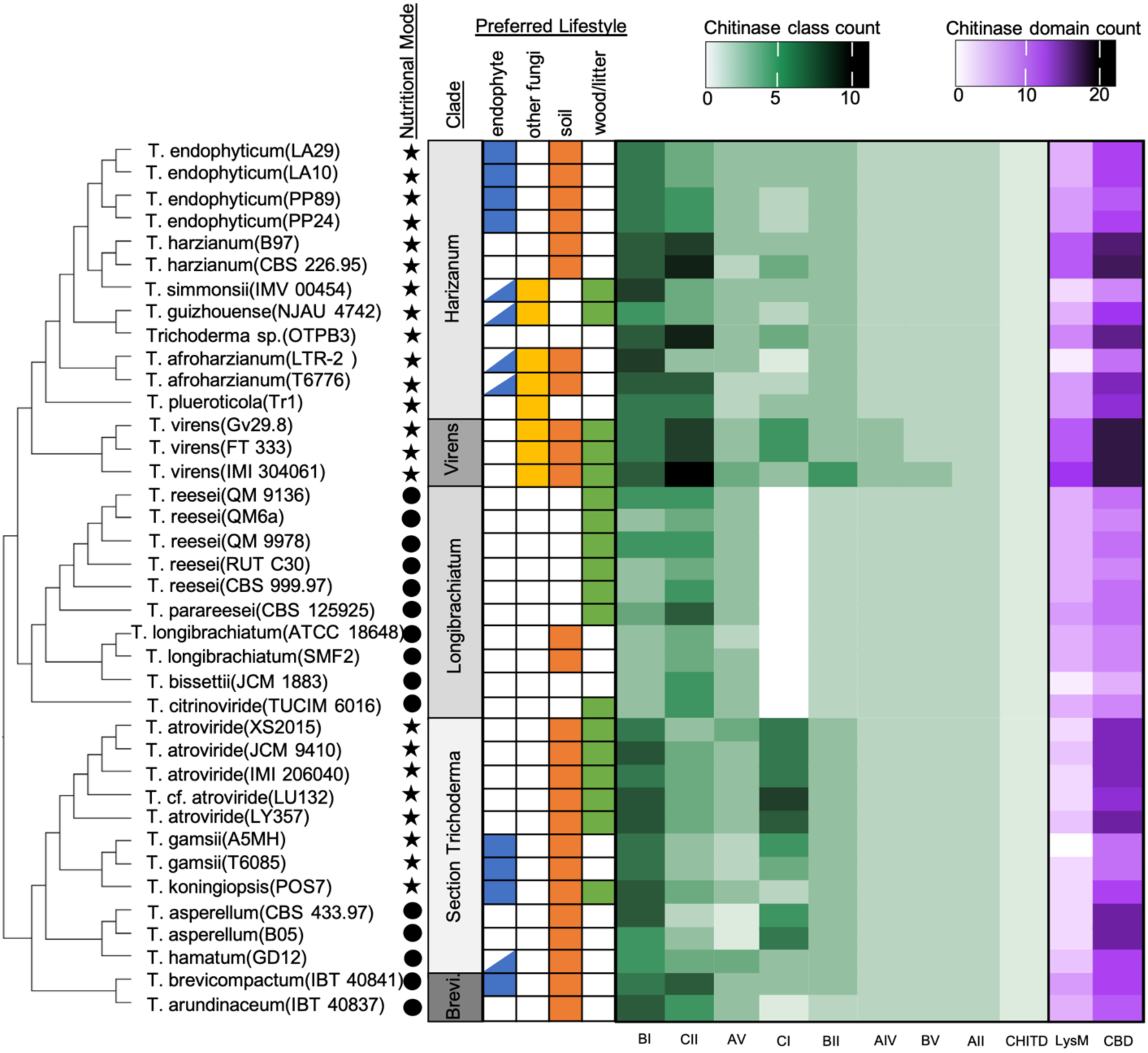
Distribution and amount of chitinase gene classes per genome varies across isolates and *Trichoderma* clade. Heatmap indicating the relative numbers of chitinase classes (green heatmaps) and chitinase domains (purple heatmaps) found in the genomes of 38 different *Trichoderma* isolates. Half-filled cells indicated a potential endophytic lifestyle for that species based on sequence identity and isolation in GenBank. Species lifestyle is indicated by color shading in cells, nutritional mode is indicated by either a circle (saprotroph) or a star (mycotroph). Clade “Brevicompactum” is abbreviated as “Brevi”.

Endophytic genomes did not contain a greater diversity of total chitinase genes compared to non-endophyte genomes (Fig. S7C), or have a different total number of individual chitinases classes (Fig. S22A). However, endophytic genomes did contain a higher percentage of class BII chitinase genes in their genomes as compared to non-endophytic genomes (Fig. S22B). Blomberg’s K and Pagel’s Lambda analysis suggests that the distribution of all chitinase gene classes, including BII, are either consistent with the *Trichoderma* phylogeny, not overdispersed in the phylogeny, or not overrepresented in endophyte genomes (Fig. S23A, B). Mycotrophic genomes contain a significantly greater amount of chitinase genes overall (Fig. S16C) as compared to saprotrophic genome, and mycotroph genomes contain greater numbers of BI, BII, and CI chitinase genes (Fig. S24A). Calculation of Blomberg’s K and Pagel’s lambda for the chitinase class counts and total chitinase gene count suggests that, similar to the endophyte:non-endophyte genome comparison, the number of total chitinase genes per genome is consistent with the *Trichoderma* phylogeny (Fig. S25).

There are significant differences in the diversity and proportions of chitinase genes to the total number of genes among the different *Trichoderma* clades (Fig. S26A,B). Virens, Brevicompactum, and section *Trichoderma* contain the highest proportions of chitinase genes, and Longibrachiatum and Harzianum contain the lowest (Fig. S26B). CI chitinase genes are completely absent in Longibrachiatum, while all other *Trichoderma* genomes contain at least one CI gene (Fig. 4). Chitinase repertoires are distinguished by the presence/absence of genes that differ in the number of lysin domains (LysM) or chitin-binding domains (CBDs); the percent of LysM domains per chitinase gene is lowest in section *Trichoderma* (Fig. S27C), and the percent of CBD per chitinase gene is highest in Virens (Fig. S27D). The variation in LysM domains corresponds to the distribution of LysM-containing CII chitinases, which are fewer in section *Trichoderma* (Fig. 4).

The *T. endophyticum* genomes have chitinase repertoires similar to other species in Harzianum (Fig. 4). The four *T. endophyticum* genomes have an average of 26 chitinase genes each, with a near-identical distribution of these genes between the four genomes. Isolates PP24 and PP89 contain two CI and five CII chitinases, while LA10 and LA29 contain three CI and four CII chitinases, respectively (Fig. 4). Domain architecture among the 26 chitinases in *T. endophyticum* strain PP24 is diverse, even among closely related genes (Fig. S28). The number of LysMs and CBDs are variable within chitinase classes, and a single member of the BV chitinase clade was inferred to contain an RTA1 Superfamily domain. LysM domains are present in all CII chitinase genes, with either one or two copies per gene. The *T. endophyticum* chitinases collectively contain fewer LysM domains than their closest relatives, the two *T. harzianum* genomes, primarily due to the higher number of LysM-containing CII chitinases in the *T. harzianum* genomes (Fig. 4). CBD is present as a single copy in all *T. endophyticum* PP24 CII genes, and a single copy per gene is present in some but not all BI and BII genes (Fig. S28).

### Xylariales-like ergot alkaloid BGCs are found in *Trichoderma* species

In the course of identifying known BGCs across *Trichoderma* using antiSMASH’s knownclusterblast [66] module, we incidentally noted a cluster of 11 homologs of ergot alkaloid BGC genes (cluster average bit score 1027.5, average amino acid identity 55.75%) in *T. brevicompactum* IBT40841 and *T. arundinaceum* IBT40837 (Brevicompactum clade, Table S5). These clusters include a hypothetical protein and 11 ergot alkaloid biosynthesis homologs (UniProt): elymoclavine monooxygenase (*cloA*), chanoclavine synthase catalase (*easC*), lysergyl peptide synthetase subunits 1 and 2 (*lps1, lps2*), chanoclavine-I dehydrogenase (*easD*), chanoclavine-I aldehyde oxidoreductase (*easA*), argoclavine dehydrogenase (*easG*), chanoclavine-I synthase oxidoreductase *(easE*), dimethylallyltryptophan N-methyltransferase (*easF*), dimethylallytryptophan synthase (*dmaW*), and a dioxygenase/oxygenase (*easH*). The hypothetical protein is not reported from any other ergot alkaloid BGC and is annotated as a nuclear transport factor 2 in InterPro or small polyketide cyclase (SnoaL) in UniProt [67].

To determine the evolutionary origin of the unexpected ergot alkaloid BGCs, we conducted a phylogenetic analysis of each gene in *T. brevicompactum’s* BGC, including homologs recovered from Eurotiales (Eurotiomycetes, Ascomycota), Helotiales (Leotiomycetes, Ascomycota), Hypocreales (Sordariomycetes, Ascomycota), Onygenales (Eurotiomycetes, Ascomycota), and Xylariales (Sordariomycetes, Ascomycota) genomes (Fig. 5). Eight of twelve ergot alkaloid BGC gene phylogenies highly support (> 95% IQ-TREE ultrafast BP) two Hypocreales clades: Hypocreales I, comprised of Clavicipitaceae species, and Hypocreales II, the Brevicompactum clade of *Trichoderma*, which is within a clade of predominantly Xylariales species. Only *cloA* supports (100% ultrafast BP) a single Hypocreales clade. *Aspergillus coremiiformis* CBS555.377 (Eurotiales) and *Pseudotulostoma volvatum* 6Q3QHG6JMA (Eurotiales) homologs are recovered with Brevicompactum clade and Xylariales species in 9 and 3 gene phylogenies respectively. All other homologs from Eurotiales species are placed on different branches in each gene phylogeny. We only recovered homologs of *easC* and a hypothetical protein (Tribre_123779) from other *Trichoderma* species. The *easC* homologs in other *Trichoderma* are part of a clade of non-ergot alkaloid BGC *easC* homologs and a *T. brevicompactum easC* paralog (100% ultrafast BP) that is distal to the cluster. The Tribre1_123779 phylogeny places the Brevicompactum clade and other *Trichoderma* homologs in two distinct subclades nested within Xylariales species homologs.

**Fig 5.**
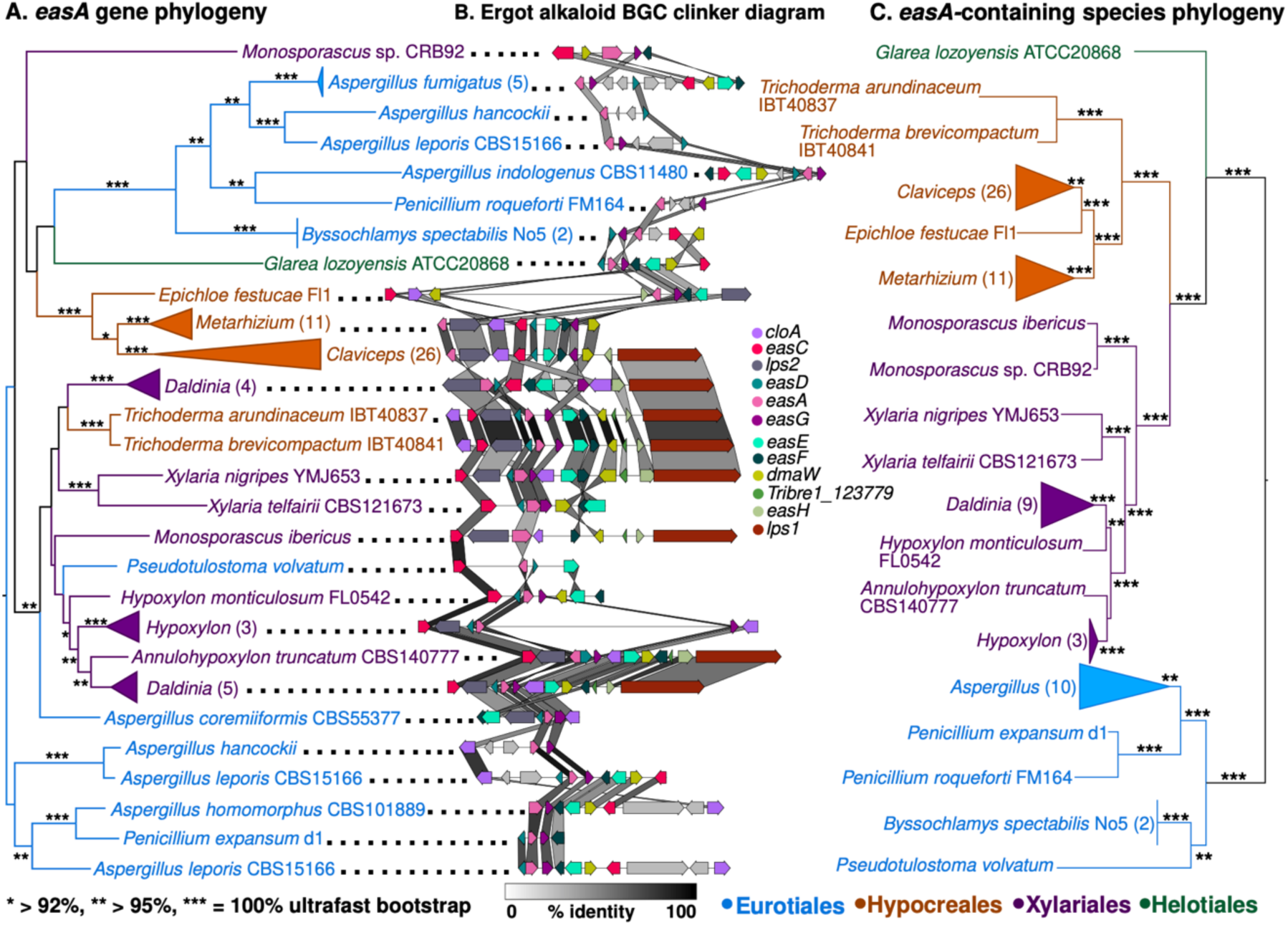
Horizontal transfer of ergot alkaloid BGCs from Xylariales to Brevicompactum and Eurotiales. (A) IQ-TREE maximum likelihood gene phylogeny of *easA* with ultrafast bootstrap branch support. (B) Shared synteny and sequence similarity of ergot alkaloid BGCs. Gene-gene percent identity is highlighted in connecting shade. (C) Species phylogeny of genomes in *easA* gene phylogeny. Tip labels in (A) and (C) are colored by taxonomic order.

The gene order in the *Xylaria nigripes* YMJ653 (Xylariales) ergot alkaloid BGC is most similar to those in Brevicompactum genomes (mean protein amino acid identity 58.5%). Eleven of twelve syntenic homologs are in the same relative position in *Xylaria nigripes* YMJ653 and *T. brevicompactum*, whereas Clavicipitaceae (Hypocreales) species BGCs have a largely different gene order (mean protein amino acid identity 53.5%). Additionally, Tribre1_123779 is located in a clade of Xylariales homologs, but we did not recover homologs in Clavicipitaceae species. The homologs of Tribre1_123779 in other *Trichoderma*, which lack ergot alkaloid BGCs, are part of a two-gene NRPS cluster they share with *Annulohypoxylon maetangense* CBS123835 (Xylariales).

## DISCUSSION

The growing resource of publicly available *Trichoderma* genomes makes it possible to search for genes and genomic features associated with particular *Trichoderma* lifestyles, including the endophytic lifestyle [68]. Yet despite the prevalence of *Trichoderma* as endophytes [12,23,69,70], few sequenced *Trichoderma* genomes were originally isolated as endophytes.

### *Trichoderma* species with a recorded endophytic lifestyle have a greater metabolic potential than non-endophytic species

Using the diversity of DGCs and BGCs in a genome as a proxy for the metabolic potential of *Trichoderma* species, we found that endophytic *Trichoderma* species have greater metabolic potential than their soil-or litter/wood-dwelling relatives. However, these results are difficult to disentangle due to the relationship between lifestyle and phylogeny within the *Trichoderma* genus, as well as the complexity inherent in the endophytic fungal lifestyle. The relative enrichment of metabolic potential in the endophytic *Trichoderma* species could result from unique selection pressures on endophytic fungi, a consequence of the natural complexity of the endophytic lifestyle or some combination thereof. Our findings, along with previous reports [26], contribute to the hypothesis that a general trend of increased metabolic capabilities is associated with the fungal endophytic lifestyle.

One potential explanation for the relative enrichment of DGCs in endophytes is the expanse of host plant chemical defenses that endophytic fungi must resist to persist inside the host plant. There is conflicting evidence about the response of the host plant to the colonization by an endophyte. Some evidence suggests that certain endophytic fungi elicit different defense responses compared to those triggered by pathogenic fungi [71, 72]. Other evidence suggests that upon the initial colonization, plant receptors indiscriminately identify both pathogenic fungi and endophytic fungi as intruders [73–76], triggering plant immune responses which include the production of defensive secondary metabolites. These chemical defenses include aromatic compounds and phenolics [77], which can negatively impact fungal growth and survival [78, 79]. Despite being a potential benefit to the host plant, the endophyte must contend with the same profile of defensive metabolites that restrict potential pathogens [80]. Although plant-produced secondary metabolites are present in soil, litter, and wood and therefore influence microbial growth and activity [81, 82], higher concentrations of induced defensive metabolites in the plant may provide focused selection for their degradation by endophytic fungi, as previous research suggest that metabolite-degrading gene clusters distribution is shaped by both fungal phylogeny and ecology [83]. Additionally, endophyte species that are horizontally transmitted and are potentially exposed to defensive metabolites from diverse hosts may benefit from a greater diversity of DGCs. A more diverse repertoire of DGCs may promote endophyte survival in the host plant by improving the breadth of host plant defensive compounds the endophyte can degrade.

The unique pressures presented by the host plant may explain in part the pattern of DGC presence and absence in the *Trichoderma* dataset. The elevated DGC diversity in endophytic *Trichoderma* species is underpinned by increased numbers of ARD, FAD, and NAD DGCs, which is consistent with previous research suggesting that fungal endophyte genomes are enriched in ferulic acid decarboxylase (FAD) DGCs, and plant symbiotic fungal genomes are enriched in DGCs containing aromatic ring-opening dioxygenase (ARD) or naringenin 3-dioxygenase (NAD) (Gluck-Thaler and Slot, 2018). Living plants contain a variety of aromatics and flavonoids, which are either constitutively expressed or induced as plant defenses against pathogens, pests, and herbivores [77,84,85]. Given that these metabolites are the likely targets of ARD and NAD DGCs, a higher diversity of these DGCs may contribute to fungal endophytes’ success in persisting within the host plant. However, endophytic species not in Harzianum did not contain ARD (Fig. 1), suggesting that ARD may not be essential for an endophytic lifestyle. The Longibrachiatum clade, which contains only species found on plant litter/soil and/or wood, was the only *Trichoderma* clade to completely lack NAD DGCs (Fig. 1), consistent with these fungi not often being exposed to these compounds in their habitat. No DGCs containing vanillyl alcohol oxidases were detected in any of the *Trichoderma* genomes (Fig. 1). This class of DGC was previously found to be enriched in soil-dwelling, saprotrophic genomes, which may benefit from lignin decay enzymes (Gluck-Thaler and Slot 2018), suggesting that *Trichoderma* species may avoid or utilize other mechanisms to tolerate lignin byproducts.

The higher diversity of BGCs found in endophytes may allow them to overcome the unique pressures of an endophytic lifestyle. For example, metabolites that mimic plant hormones may facilitate communion with the host plant or produce plant growth-promoting compounds [17]. Some plant growth-regulating compounds produced by *Trichoderma* species are harzianolide [86–88], harzianic acid [89], and 6-pentyl-2H-pyran-2-one [90, 91]. BGCs in *Trichoderma* species may also produce siderophores that sequester iron [50, 92] or metabolites that inhibit antagonistic microbes [93–95], or in the case of harzianic acid, both iron sequestration and antifungal ability [96]. Although the majority of *Trichoderma* secondary metabolite products have not been linked to their respective BGCs [68], a greater diversity of BGCs is expected to facilitate a wider variety of interactions between a fungus and its surrounding environment, potential hosts, and other microorganisms.

Due to “cross-talk” between plant and endophyte metabolic pathways, a diverse metabolic gene cluster repertoire may lead to even further diversification of the metabolite profile in the holobiont [97]. *Cola acuminata* produces camptothecin, and a common endophyte of this plant (*Fusarium solani)* produces camptothecin precursors [98, 99]. In this system, the fungus is missing a key enzyme that is present in the plant that allows the production of camptothecin. However, since the plant can transform the fungal metabolites, this increases the availability of camptothecin in the plant. The opposite of this case is also true, as endophytic fungi are known to modify terpene metabolites produced by their host plant [100]. These observations are consistent with the “mosaic effect” theory, which states that endophytes create a variable metabolite composition within and between plant organs that would otherwise contain a uniform profile of metabolites, thereby increasing the defense ability of the plant [101].

Increased diversity of both DGCs and BGCs in endophytes may ultimately improve the fitness of the holobiont (a plant and all closely-associated microorganisms) by contributing to the overall collection of defensive metabolites available to the plant and acting as the host plant’s “acquired immune system” [93]. Endophytes increase host availability in the ecosystem by increasing host protection from pathogens and herbivores. Endophytes can produce novel metabolites in the host plant or produce similar metabolites already synthesized by the host plant [98,102,103]. Fungal endophytes of several grass species (e.g., *Epichloë* spp.) produce a variety of novel defense alkaloids which can directly protect the host plant by reducing feeding by both invertebrate and vertebrate herbivores [104, 105]. *Trichoderma* species specifically are known to produce a multitude of fungal defensive secondary metabolites beneficial to the host plant [94, 106], such as the antifungals trichodermin and harzianopyridone [86, 107]. Fungal endophytes may also influence the blend of volatile organic compounds exuded by the plant, which attract predators of the herbivorous aphid species, thereby decreasing herbivory of the host plant [108]. Taken together, our findings suggest that the selective pressure for increased overall fitness of the holobiont via metabolite diversity may select for more metabolically diverse pathways in endophytic fungal genomes.

Alternatively, elevated metabolic diversity in endophytic genomes may be associated with these fungi being able to exploit multiple lifestyles throughout their lifecycle [26,109–111], including utilizing decaying plant biomass [112, 113]. Indeed, each of the endophytic *Trichoderma* isolates in our dataset are more or less “ecological generalists” that have been associated with multiple lifestyles (Table S1). Fungi with more complex interactions with other organisms, such as plants or other fungi, tend to produce a greater diversity of bioactive compounds [115]. For example, *T. virens*, capable of multiple lifestyles (wood/litter-, soil-, and fungi-dwelling), is well-known for producing a variety of bioactive metabolites [50,62,116]. In this study we also observed that genomes of *Trichoderma* species with multiple potential lifestyles have greater numbers of DGCs and BGCs than genomes of *Trichoderma* species only exhibiting a single lifestyle (Fig. S29A,B). In contrast, metabolic gene cluster diversity is reduced in Longibrachiatum, which contains exclusively single-lifestyle species (Fig S11A,B). Rather than an increase in metabolic diversity in multi-lifestyle genomes, there may have been a past reduction in diversity in single-lifestyle genomes. Species in Longibrachiatum were shown to be able to degrade comparatively fewer metabolites in laboratory metabolite digestion challenges compared to species in Harzianum or other plant-related *Trichoderma* species [5], consistent with our findings of their metabolic potential (Fig. S11A, Fig. S29A). Specifically, all species in Longibrachiatum completely lack ARD and NAD DGCs (Fig. 1), despite these species living exclusively in wood or plant litter where aromatic compounds are present as products of wood decay. Previous studies on the succession of fungal species in leaf litter and dead wood indicate that *Trichoderma* species are more abundantly found as late-term colonizers, active only after other microbe community members have degraded or modified complex compounds such as lignin or complex metabolites [117–121]. By depending on other members of the substrate microbiome to degrade aromatic compounds [77, 122], litter- and soil-dwelling *Trichoderma* species would not be under the same selective pressure as endophytic species to retain DGCs, resulting in the loss of these DGCs. Metabolic diversity may reinforce ecological generalism in some lineages, while specialized lifecycles in others favor metabolic streamlining.

The trends observed in the metabolic capabilities of endophytic *Trichoderma* are exemplified by newly sequenced *T. endophyticum* genomes isolated from tropical trees. This study shows that *T. endophyticum* is an extreme example of the characteristics typically exhibited by endophytic *Trichoderma* species. The four *T. endophyticum* genomes are among the genomes with the greatest numbers of DGCs, BGCs, and metabolic genes as compared to their close relatives (Fig. S2). Previous comparative analyses suggest that endophyte genomes tend to have comparatively larger genomes [26], a trait that was also observed in our four *T. endophyticum* genomes (Fig. SS4). The four *T. endophyticum* genomes were above the trend comparing number of DGCs and BGCs to genome length (Fig. S4A,B) and the proportion of DGCs and BGCs to the total gene number (Fig. 5A,B) and exceeded their closest relatives in the Harzianum clade in DGC measures (Fig. S4A, S5A). *Trichoderma endophyticum* may be of interest in natural product discovery due to the comparative richness of its metabolic diversity [123, 124]. However, this study only evaluated four *T. endophyticum* genomes from two geographic locations and two host plant species. Given that *T. endophyticum* isolates are more similar based on geographic location and host plant species [3], more representatives of this species should be sequenced from different host tree species and from a wider geographical range in order to understand more about the intra-species diversity of *T. endophyticum*.

### Mycoparasitism-related genes

Each of the 38 *Trichoderma* genomes contains multiple genes in orthogroups made from our list of 746 key mycoparasitism genes. The distributions of the mycoparasitism gene repertoires are consistent with phylogenetic relationships. There was no indication of grouping based on fungal lifestyle, even after adjusting for phylogenetic relationships within the dataset (Fig. S20). Although the main indicator of mycoparasitism gene diversity and distribution was phylogeny, mycotrophic genomes contained significantly greater numbers of the mycoparasitism-related genes in mycotrophic genomes as compared to saprotroph genomes (Fig. S16D).

Overall, endophytic *Trichoderma* genomes had greater abundances of mycoparasitism-related genes than non-endophytic genomes (Fig. S7D). However, this difference was not as distinct as the difference in abundances between mycotrophic and saprotrophic genomes (Fig. S16D). Potentially, the relative abundance of mycoparasitism genes could be due to some of the mycoparasitism genes being beneficial for creating or maintaining a host-endophyte symbiosis. For instance, a gene encoding an aspartyl protease (OG0009580) was overrepresented in both mycotrophic and endophytic genomes (Table S3); aspartyl proteases are cell wall degrading enzymes known to be produced upon competition of a fungal target [125] as well as upon colonization of plant host [126]. The ability to digest a plant cell wall would be positively selected for horizontally-transmitted endophyte species, as this ability would increase the chances of reaching the inside of the plant. Our results support similar findings, which suggest that certain cell-wall degrading enzymes are overrepresented or expanded in fungal endophytes, also theorized to be useful for the initial infection and colonization of the host plant [127]. Alternatively, the relative abundance of mycoparasitism genes in endophytic genomes may simply be due to the evolutionary history of *Trichoderma*, which is hypothesized to have a mycoparasitic origin [4], as the mycoparasitic nutritional mode is widespread throughout this genus and most species with a recorded endophytic lifestyle also have a mycotrophic nutritional mode.

### Chitinase prevalence and other key mycoparasitism-related genes are not correlated with an endophytic lifestyle in *Trichoderma*

Each *Trichoderma* genome contained one or more chitinase genes of different classes (Fig. 4). Mycotrophic genomes contained a significantly greater amount of total chitinase genes as compared to saprotrophic genomes (Fig. S16C). Mycotrophic genomes contained greater numbers of BI, BII, and CI chitinases (Fig. S24A). The relative abundance of chitinases in mycotrophic genomes is not unexpected since chitinase production is key to mycoparasitism, many mycoparasitic *Trichoderma* species are well-known chitinases producers [128], and as previously stated, mycoparasitism is the ancestral state of *Trichoderma* [50]. Similar to the results for metabolite gene clusters and mycoparasitism gene repertoires, we also determined that the overall pattern of chitinase gene distribution was consistent with *Trichoderma* phylogeny (Fig. S25A,B).

One of the benefits endophyte confer to their host plant is direct competition of invading fungi via mycoparasitism [129, 130]. Due to this selective pressure for mycoparasitic ability in endophyte genomes, we investigated if endophytic *Trichoderma* species have expanded chitinase repertoires. However, we did not find any difference in the overall number of chitinase genes (Fig. S7D) or individual number of chitinase gene classes (Fig. S22A) between endophytic genomes and non-endophytic genomes. Only when we looked at percent of total gene number did we identify any differences in endophytic and non-endophytic chitinases content; the percent of chitinase BII genes in each genome is significantly greater in endophytes (Fig. S22B). This relative increase in BII chitinases could simply be due to the mycoparasitic history of *Trichoderma* [50], because we did not identify a significant signal associated with an endophytic or mycoparasitic lifestyle when we controlled for phylogenetic signal (Fig. S23A,B).

### Horizontal transfer of an ergot alkaloid cluster is consistent with a defensive role in endophytes

BGC repertoire diversification in *Trichoderma* is in part shaped by horizontal gene transfer between Eurotiales, Hypocreales, and Xylariales. Given the precedent of horizontal transfer in Hypocreales and Brevicompactum clade [131–133], we were not entirely surprised to identify BGC acquisitions among *Trichoderma* species, including the ergot alkaloid BGC [134]. The species phylogeny places Clavicipitaceae and *Trichoderma* within Hypocreales. However, eight of twelve ergot alkaloid gene phylogenies support Brevicompactum’s placement within a Xylariales-dominant clade (> 95% ultrafast BP per gene) apart from the Clavicipitaceae clusters. HGT is supported by the shared gene order between *Xylaria nigripes* YMJ653 and *T. brevicompactum,* which differs from the gene order in Clavicipitaceae. Additionally, we recovered a novel ergot alkaloid gene family (Tribre1_123779) that is shared between *T. brevicompactum* and Xylariales ergot alkaloid clusters but absent in Clavicipitaceae, further supporting the HGT. The ergot alkaloid BGC was likely obtained after Brevicompactum diverged from other *Trichoderma* because the only homologs we recovered in other *Trichoderma* are distant *easC* paralogs and independently transferred xenologs of Tribre1_123779. Interestingly, the independent transfers of Tribre1_123779 to other *Trichoderma* species are part of a different NRPS cluster acquired from Xylariales species that also lack the ergot alkaloid BGC (100% ultrafast BP). We further identified ergot alkaloid BGCs in two Eurotiales species that are more closely related to the Xylariales clusters and Brevicompactum clusters than other Eurotiales’, suggesting there have also been inter-class HGTs involving Xylariales. These findings add to previous work that suggested the origin of the Clavicipitaceae (Hypocreales) ergot alkaloid BGC was by HGT from Eurotiales species [135]. HGT of metabolic gene clusters to *Trichoderma* has been previously reported [136], including a recent report of HGT of a PKS BGC to Brevicompactum from Eurotiales [134]. Our finding of multiple novel HGTs between Eurotiales, Xylariales, and *Trichoderma* (Hypocreales) provides evidence of a potential horizontal transfer highway between these orders.

HGT is thought to indicate shared selection pressures in the donor and recipient lineages and can suggest possible ecological roles of the products of the transferred BGC [34]. Ergot alkaloids are generally considered to have roles in animal-associated fungal niches [137]. They can act as anti-herbivory compounds in the plant endophytic/pathogenic *Claviceps* and *Epichloë* species [138]. They are also upregulated during insect colonization by *Metarhizium* insect pathogens [139]. The most closely related clusters to Brevicompactum’s ergot alkaloid BGCs are also in animal-associated, plant-pathogenic, or endophytic Xylariales: *Daldinia childiae* (Xylariales)*, D. decipiens* CBS113046, *D. loculata* AZ0526, *D. loculata* CBS113971, *Xylaria nigripes,* and *X. telfairii* CBS121673 [140–149]. All of these species have been isolated as endophytes or from decaying wood, while *D. decipiens* is also a symbiont of *Xiphydria* woodwasps [142], and *X. nigripes* is one of several termite-associated *Xylaria* species [150]. Xylanigripones are the first described ergot alkaloids from Xylariales and are produced by *X. nigripes* [151], and these compounds are the likely product of the *X. nigripes* cluster we identified. *X. nigripes* has an extensive history of medicinal consumption in humans attributed to its diverse pharmacological activity, including effects on the nervous system, which is typical of ergot alkaloids. Given the high similarity between Brevicompactum’s ergot alkaloid BGCs and *X. nigripes’,* it is likely that Brevicompactum species produce similarly structured compounds. More broadly, the presence of novel ergot alkaloid BGCs throughout Xylariales suggests there may be undescribed ergot alkaloids from Brevicompactum and Xylariales.

The ecological function of xylanigripones is not known, nor are any shared ecological pressures that could drive HGT from Xylariales to Brevicompactum. This HGT may simply represent the potential for mycotrophs like *Trichoderma* species to acquire genetic material from hosts [37, 136]. Additionally, endophytic ecologies in Brevicompactum and Xylariales may facilitate HGT through shared habitat [152]. Indeed, the majority of Xylariales species with ergot alkaloid BGCs have been isolated as endophytes. Furthermore, the two Eurotiales clusters within the Xylariales ergot BGC donor clade are potential plant associates, including *Pseudotulostoma volvatum* [153], which is associated with tree roots [154] and *Aspergillus coremiiformis* CBS55377, an anomalous *Aspergillus* with a reduced genome, but maintenance of plant cell wall degradation enzymes [155]. We speculate that the putative products of ergot alkaloid BGCs might contribute to plant host defense in Brevicompactum endophytes, given their documented role in anti-herbivory and the widespread distribution of ergot alkaloid BGCs in Xylariales endophytes and animal-associated species.

### Limitations in analysis

It is important to note that there may be some error in assigning the gene clusters to particular cluster families due to fragmentation of the clusters or cluster core genes not being present in annotations. Fragmented secondary metabolite clusters are likely still recovered by antiSMASH if the core genes are present, and BiG-SCAPE’s model attempts to account for cluster fragmentation in cluster family grouping. However, core genes that were not called in the annotation are not detected by antiSMASH, which is an ongoing problem in the field. For DGCs, we adjusted the weights of BiG-SCAPE’s cluster relationship network, so these weights may not optimally group DGCs into families. To better account for fragmented and DGC grouping, we opted to use BiG-SCAPE clan designations, which are more crude.

Other limitations of this study include difficulties assigning lifestyles to the different *Trichoderma* species due to its ubiquitous nature and oftentimes cryptic nutritional modes and/or lifestyles [3, 4]. For these analyses, *Trichoderma* lifestyles were determined by finding two or more reports of confirmed *Trichoderma* species being isolated from the internal tissues of a living plant.

## METHODS

### *T. endophyticum* DNA extraction and sequencing

Four isolates of *T. endophyticum* were isolated from the sapwood of living rubber trees (*Hevea* spp.) from Peru and identified by Sanger sequencing the ITS region. Isolates were individually grown in Potato Dextrose Broth (PDB) in sterile flasks on a shaker plate for several days in darkness. Mycelium was filtered from the PDB and rinsed twice with 50m of sterile MiliQ water. The mycelium was frozen using liquid nitrogen and ground into fine dust using a mortar and pestle. We extracted DNA from the pulverized mycelium using the DNeasy Plant Minikit (Qiagen), which was then sequenced with both short (Illumina) and long-read (Oxford Nanopore Technology) technologies. We prepared the DNA for long-read sequencing using the Oxford Nanopore Technologies library preparation kit SQK-LSK108, using the standard kit instruction and G-Tube fragmentation. The prepared DNA was individually sequenced using the Nanopore MinION Mk1B and flow cell with R9.4.1 chemistry, resulting in approximately 10x coverage of each *T. endophyticum* genome. Nanopore raw sequence data was basecalled with Albacore v.2.3.1. The Illumina PE150 sequencing was performed using a NovaSeq 6000 sequencer.

### Genome assembly

We retrieved publicly available raw SRA data from NCBI and JGI for 35 *Trichoderma* strains. For both the publicly available read data and our own *T. endophyticum* reads, we removed adapters and low-quality reads from the short-read data with Trimmomatic v0.36 [156] and assessed overall read quality with FastQC v0.11.7 [157]. We used SPAdes v3.12.0 [158] to assemble each *Trichoderma* genome, using both the Illumina and Nanopore reads to produce hybrid assemblies for the four *T. endophyticum* isolates. All assemblies had their quality metrics (e.g., N50) assessed with QUAST v4.6.3 [159] and coverage assessed with BBmap v37.93 [160].

### Genome annotation

All genomes were annotated using a MAKER v2.31.8 [161] annotation pipeline. Genome annotation for the entire 39-genome dataset was performed identically for each separate genome. For each genome, de novo repeat libraries were created with RepeatModeler [162]. Our MAKER2 pipeline was run with three iterations in order to fine-tune the genome annotations. In the first iteration, protein and EST data from the same or closely related *Trichoderma* species were retrieved from NCBI for each genome. The first iteration of MAKER used the repeat library and provided EST and protein data to generate individual preliminary gene annotations for each genome. For the second iteration of the MAKER2 pipeline, we provided *ab initio* gene predictions from SNAP v.2013-02-16 [163] (produced from preliminary gene predictions from the first iteration) and Augustus v.3.3 [164] (trained using BUSCO v.3.0.1 [165]), and GeneMark v4.32 [166]. SNAP and Augustus were both retrained using the high-quality predictions from iteration 2, and this refined dataset was supplied to MAKER2 for the final iteration to produce the final genome annotations for each isolate (option keep_preds = 1).

### Genome dataset curation

The complete dataset of 39 *Trichoderma* genomes includes ecologically and geographically diverse isolates. The genes encoding translation-elongation factor 1 alpha (TEF1) and RNA polymerase II gene (RPB2) sequence were identified in each genome and compared to their respective GenBank type specimen sequences using BLASTn. In the case of several *Trichoderma* isolates, the original species identification was determined to be incorrect, and the species was re-labeled. Available in Table S1 are details of each isolate’s revised species identification, assembly source (public or re-assembled), nutritional mode, species lifestyle, and country of origin.

### Assignment of ecological data to isolates

Lifestyle and nutritional mode data for each species was collected using the same approach used in Chaverri and Samuels (2013). Briefly, lifestyle information (i.e., endophyte, on decaying plant material, in soil, and on other fungi) was based on published research, and nutritional mode (i.e., mycotrophy vs. saprotrophy) was based on ancestral character reconstructions, phylogenetic affinities, and confirmed antagonism experiments. Two *Trichoderma* strains (*T. bissettii* [JCM 1883] and unknown *Trichoderma* species [OTPB3]) were left as “unknown” because no ecological information was available. For each *Trichoderma* species, we assigned a potential endophytic lifestyle by identifying isolates reportedly isolated from living plant tissue, documented on NCBI. To be a potential endophyte, we required at least two isolates from separate sources with a blastn match for one of the following: TEF1, 100% coverage and >99.5% identity, and RPB2, 100% coverage and >99.0% identity. For *T. arundinaceum*, there was no RPB2 sequence from type material available on NCBI, so we looked for matches with TEF1, 100% coverage and >99.5% identity and ITS 100% coverage and >99.5% identity.

### Ortholog supplementation and identification

To find orthologous sequences across the 39 genome set, we provided the final protein datasets for each *Trichoderma* genome to OrthoFinder v2.2.6 [167]. The resulting orthology datafiles and the MAKER protein files were processed with Orthofiller [168] to improve the genome annotations and identification of orthologs. All subsequent analyses used the post-Orthofiller protein datasets.

### Phylogenomic tree creation

Using the protein sequences of the single-copy orthogroups produced from OrthoFinder, we placed each orthogroup into an automated pipeline that built alignments for each orthogroup using mafft v7.407 [169], and alignments were automatically curated using the automated1 algorithm in TrimAI v1.4 [170]. RAxML was used for phylogenetic analyses [mapping percentage of 100 rapid bootstraps to the best-scoring ML tree], which resulted in 154 SCO trees with a median BP ≥98%. These 154 SCO trees were used to create the majority rule extended consensus tree using RAxML v8.2.11 (-m GTRCAT) [171]. We produced an alignment consensus phylogenomic tree from the 154 SCO IQ-TREEs with >98% BP support in IQ-TREE v2.0.3 (-bb 1000 -bsam GENESITE -m TEST) [172].

### Degradative gene cluster identification

We used a custom pipeline (https://github.com/egluckthaler/cluster_retrieve) [30] identify putative gene clusters associated with secondary metabolite degradation in the 39 annotated *Trichoderma* genomes. This custom script searched each genome for multiple gene cluster models containing particular “core” genes, or genes known to be important for processing secondary metabolites. We searched for gene clusters containing the following 13 core genes: aromatic ring-opening dioxygenase (ARD), benzoate 4-monooxygenase (BPH), ferulic acid esterase 7 (CAE), catechol dioxygenase (CCH), epicatechin laccase (ECL), ferulic acid decarboxylase (FAD), pterocarpan hydroxylase (PAH), naringenin 3-dioxygenase (NAD), phenol 2-monooxygenase (PMO), quinate 5-dehydrogenase (QDH), salicylate hydroxylase (SAH), stilbene dioxygenase (SDO), and vanillyl alcohol oxidase (VAO) (Gluck-Thaler and Slot 2018). Homologous genes at each locus were defined by a minimum BLASTp (v2.2.25+) bitscore of 50, 30% amino acid identity, and a target sequence alignment 50-150% of the query sequence length. Homologs of the query genes were considered clustered if a maximum of 6 intervening genes separated them; gene clusters on the same contig were consolidated if they were separated by less than 30kb. DGCs containing the same core gene are referred to as belonging to the same cluster class. When comparing DGCs across different genomes, DGCs exhibiting similar organization with a high degree of sequence similarity will be referred to as the same DGC family.

### Biosynthetic gene cluster identification

We predicted secondary metabolism clusters by submitting our *Trichoderma* annotations through the antiSMASH v5.0b [173] pipeline with glimmerhmm gene prediction enabled. Clusters were analyzed via the ‘knownclusterblast’ module to identify potential homologs of MIBiG v2.0 [66] database entries. As additional quality filters, we only referenced MIBiG accessions with 3 or more genes and removed knownclusterblast genes with bit scores lower than 100, redundant hits within each cluster, clusters with aggregate bit scores less than 100 x cluster gene quantity, and any clusters with less than 60% of the respective MIBiG cluster’s gene quantity. We parsed and compiled data from the antiSMASH and knownclusterblast output using *smashStats.py* (gitlab.com/xonq/mycotools). BGCs are grouped in different classes respective to their association with different secondary metabolite compound classes, according to antiSMASH nomenclature (e.g. polyketides, non-ribosomal peptides, terpenes, indole etc.).

### Cluster family classification

We grouped gene clusters into homologous cluster families using BiG-SCAPE v1.0.1 [174]. Secondary metabolism clusters were classified into cluster families using default parameters while referencing MIBiG v2.0. For degradative clusters, we modified BiG-SCAPE’s default model weights to 63% protein domain sequence similarity, 35% protein domain presence-absence, 2% conserved synteny, and a core gene boost set to 2x. *Trichoderma* cf*. atroviride* (LU140) produced a very fragmented genome which skewed the identification of MGCs, and as such, this genome was removed from the subsequent BGC and DGC cluster family designations. The differences in the overall counts of DGCs, BGCs, and their respective gene cluster classes between *Trichoderma* clades were determined using ANOVA or Kruskal-Wallis, and differences between both endophyte vs non-endophyte and mycotrophic vs saprotrophic were determined using a 2-sample t-test or Wilcoxon rank sum test.

### Mycoparasitism-related genes analysis

Using previously published results of differential gene expression analyses from RNA-seq and oligonucleotide tiling array data, we compiled a list of 746 genes from *T. atroviride, T. virens, T. reesei,* and *T. harzianum* that were upregulated during at least one time point or treatment during mycoparasitism assays compared with control time points or treatments [62–64]. For each of these 746 mycoparasitism-related genes, we identified the highest scoring hit in each *Trichoderma* genome using BLASTp (E value threshold = 0.001). Any orthogroup containing a highest scoring hit to a mycoparasitism-related gene was considered to be a mycoparasitism-related orthogroup. The differences in the overall counts of mycoparasitism-related genes between *Trichoderma* clades were determined using ANOVA or Kruskal-Wallis, and differences between both endophyte vs non-endophyte and mycotrophic vs saprotrophic were determined using a 2-sample t-test or Wilcoxon rank sum test.

### Chitinase analysis

We used an updated kingdom-level GH-18 chitinase ontology [65] to classify chitinases across the dataset. Chitinase sequences were first identified by hmmsearch for proteins containing Glyco_hydro_18 (PF00704) domain in hmmer-3.1b2 [175]. Chitinase genes from all *Trichoderma* species were then incorporated into a multifasta file with diverse representatives of each chitinase class in Goughenour et al. (2020) [65]. The sequences were then aligned in mafft v7.487 [169] and analyzed in FastTree v2.1.10 with default settings [176]. Sequence chitinase classes were then manually assigned based on placement in the FastTree phylogeny. Domain architecture for *T. endophyticum* PP24 chitinases (Figure S12) was determined using a CDD search [177] and mapped to an IQ-TREE (v. 1.6.11) phylogeny inferred under the LG+F+R3 model, which was determined by Model Test to be the best fit according to the BIC, with support computed by 1000 ultrafast BS replicates. The differences in the overall counts of chitinase gene classes between *Trichoderma* clades were determined using ANOVA or Kruskal-Wallis, and differences between both endophyte vs non-endophyte and mycotrophic vs saprotrophic were determined using a 2-sample t-test or Wilcoxon rank sum test.

### Phylogenetic correction: Fritz and Purvis’ D, Blomberg’s K, and Pagel’s Lambda

To determine how the distribution and diversity of binary traits, such as the presence/absence of individual DGC and BGC families, are consistent with the phylogeny of *Trichoderma*, we calculated Fritz and Purvis’ D statistic [178] for each individual gene cluster family using the “phylo.d” function from the “caper” v1.0.1 package [179] in R. Fritz and Purvis’ D indicates the level to which species’ shared phylogenetic history impact the presence or absence of any given trait. Fritz and Purvis’ D Statistic is a measure of “phylogenetic signal” in the distribution of a trait. A D of zero indicates that a binary trait follows the Brownian motion model of evolution, where differences in distribution can be entirely explained by the phylogenetic history. When D = 1, this indicates that a binary trait is randomly distributed. D values above 1 indicate that traits are more overdispersed than expected by the Brownian motion model, and together with ecological distribution bias (see below) could suggest a trait is associated with particular lifestyle or nutritional mode. As D falls below 0, a trait is considered underdispersed (more conserved than the Brownian motion model). For the Fritz and Purvis’s D-statistic, a significant p-value indicates that a particular gene cluster’s distribution is significantly different compared to a simulated random distribution (a phylogeny under Brownian motion), therefore its distribution is not entirely explained by its evolutionary history.

To determine how the distribution and diversity of continuous traits are consistent with the phylogeny of *Trichoderma*, we calculated Blomberg’s K [180] and Pagel’s Lambda [181, 182], two different quantitative measures of the “phylogenetic signal”. Using the “phylosig” function in “phytools” v1.3-5 [183] in we determined both the Blomberg’s K (method = ‘K’) and Pagel’s Lambda (method = ‘lambda’) for continuous traits such as total number of gene clusters, total number of separate gene cluster classes, total chitinase genes, total number of separate chitinase gene classes, and total number of mycoparasitism genes. Similar to Fritz and Purvis’ D statistic, these analyses also incorporate the branch length and structure of a phylogenetic tree to determine the measure of how much the evolutionary history of taxa history drive the distribution and pattern of a particular trait.

Blomberg’s K is a scaled ratio of (variance among the trait counts between the different species/contrasts variance); when phylogenetic signal is strong, the contrasts variance is low. When Blomberg’s K is equal to 1, this indicates that the model is completely equal to Brownian motion, or that the evolutionary history completely explains the distribution and pattern of a particular trait. When K < 1, there is less strength of phylogenetic signal than expected under Brownian model evolution. When K > 1, there is a greater signal than expected. Pagel’s Lambda is a parameter that scales between 0 and 1, indicating the correlations between species relative to the relationship expected when the trait is distributed according to the Brownian motion model of evolution. When Pagel’s Lambda is equal to 1, this indicates that the model is completely equal to Brownian motion. When Pagel’s Lambda is equal to 0, this indicates that there is no phylogenetic influence on the distribution and pattern of a particular trait. As Pagel’s Lambda decreases from 1 to 0, this is indicating a progressively lesser influence of the “phylogenetic signal” on the count of a particular trait. Pagel’s Lambda and Blomberg’s K test for the null hypothesis that there is no phylogenetic signal, therefore a significant p-value indicates that there is significant phylogenetic signal for a particular genomic feature. We used an arbitrary cutoff value of 0.8 for Blomberg’s K and Pagel’s Lambda; values equal to or lower than 0.8 are considered to have the potential of indicating gene clusters with selection in their evolutionary history. A significant p-value indicates that a particular genomic feature’s distribution is consistent with its evolutionary history (follows Brownian motion model evolution).

We determined the ecological distribution bias of each trait by calculating the ratios for the mycotroph:saprotroph genomes (average count of each trait in mycotroph genomes/average count of each trait in saprotroph genomes) and endophyte:non-endophyte genomes (average count of each trait in endophyte genomes/average count of each trait in saprotroph genomes). We considered a comparative 2-fold increase (ratio value > 2) in the endophyte:non-endophyte (E:NE) and mycotroph:saprotroph (M:S) ratio to indicate genomic traits that were significantly enhanced in a particular *Trichoderma* ecology. The resulting ratios and respective phylogenetic signal statistic for each genomic feature was plotted on a scatter plot. Genomic features which had a significance level indicating that their distributions were consistent with *Trichoderma* phylogeny (consistent with Brownian motion evolution, or neutral evolution) were colored grey, points which were not entirely consistent with *Trichoderma* phylogeny and therefore were of potential interest for underpinning a particular trait were colored black.

### Ergot alkaloid BGC phylogenetic and synteny analyses

To study the evolution of the Brevicompactum clade ergot alkaloid BGC, we implemented the Cluster Reconstruction and Phylogenetic Analysis Pipeline (CRAP) across a database of 2,297 GenBank and MycoCosm [184] published fungal genomes using Mycotools v0.27.15 (gitlab.com/xonq/mycotools). We obtained the complete Brevicompactum clades’ clusters by synthesizing non-overlapping gene coordinates from our annotations with MycoCosm’s (Portal Tribre1, Triaru1). *easE* and *easF* were originally fused in Tribre1’s annotation, so to construct robust phylogenies of each gene, we split the gene via exonerate [185] v2.2.0 (maxintron 100, percent 10, protein2genome:bestfit model) referencing *X. nigripes easE* and *easF*.

We queried the refined *T. brevicompactum* cluster using CRAP, which first obtained homologous genes via BLASTp [186] with a minimum bitscore threshold 30. To increase phylogenetic reconstruction throughput, we set CRAP to implement sequence similarity clustering to truncate groups of greater than 250 homologous sequences. The sequence clustering module iteratively implements *mmseqs2* v14.7e284 [187] *easy-cluster* with a minimum query coverage +/-40% and adjusts the minimum sequence identity of each iteration until a group of fewer than 250 sequences was retrieved. Each query’s truncated homolog group was subsequently aligned via mafft v7.487 (Katoh and Standley 2013) with default settings, trimmed using ClipKIT [188] v.1.3.0 with default settings, and initial phylogenies constructed via FastTree [176] v2.1.10 with default settings. We used the full 379 homologs retrieved by BLAST for Tribre1_123779 because the sequence similarity clusters were too granular (12 genes) to contextualize its evolution. FastTree phylogenies were rooted upon an outgroup branch identified by a larger module of homologs detected via *mmseqs easy-cluster.* Alternatively, these phylogenies were midpoint rooted if the initial BLASTp retrieved homologs were under 250 sequences.

To examine shared synteny with the Brevicompactum clade ergot alkaloid BGCs, CRAP extracted 50 kB up-/downstream of each homolog, truncated the loci boundaries to homologs of query genes, and mapped the resulting synteny diagrams on each query phylogeny. To generate robust IQ-TREE phylogenies, we extracted highly supported clades (> 0.98 Shimodaira-Hasegawa test) from FastTree phylogenies that contained known ergot alkaloid BGC homologs. Alternatively, all sequences from the FastTree phylogeny were used in IQ-TREE when the FastTree was rooted with known ergot alkaloid BGC homologs [135]. We inferred robust maximum likelihood phylogenies via implementing IQ-TREE v2.0.3 with 1000 ultrafast bootstrap iterations after aligning and trimming as described above.

To investigate the origin of *Trichoderma* ergot alkaloid BGCs, we compared the topology of the IQ-TREE gene phylogenies with a species phylogeny of genomes that contain chanoclavine cyclase oxidoreductase positional homologs. We constructed the species phylogeny via IQ-TREE using a multigene partition phylogeny of 13 profile hidden markov protein models [189] that were recovered from 2,296 / 2,297 fungal genomes. We obtained the top hit of these protein models from each genome as described in [190] using *db2search.py* with an e-value cutoff of 0.01 and minimum query coverage of 50%. In preparation for phylogenetic reconstruction, the retrieved proteins were aligned via mafft, trimmed via ClipKIT (--gappy 0.7), and submitted to ModelFinder [191] via *fa2tree.py* (gitlab.com/xonq/mycotools). The best model for each protein was used to build the multigene partition phylogeny from the concatenated trimmed alignments. To visualize shared synteny with respect to the species’ phylogeny, we submitted loci obtained from the *easA* homologs to clinker v0.0.24 [192] with minimum alignment identity set to 50%.

The genome assembly, annotations, and other resource-intensive analyses were performed using the Ohio Supercomputer Center [193] services. Genome assemblies and annotations, and results from the DGC, BGC, and mycoparasitism-related gene analyses are available as supplemental information files at Dryad (https://doi.org/10.5061/dryad.r2280gbhn).

## ACKNOWLEDGEMENTS

This work was supported by a grant from the National Science Foundation (DEB-1638999) to JCS and by a Marie Skłodowska-Curie Individual Fellowship (SEP-210615880-TRADEOFF) to EGT.

## REFERENCES

1. Bailey BA, Bae H, Strem MD, Crozier J, Thomas SE, Samuels GJ, et al. Antibiosis, mycoparasitism, and colonization success for endophytic Trichoderma isolates with biological control potential in Theobroma cacao. Biological Control. 2008;46: 24–35. doi:10.1016/j.biocontrol.2008.01.003

2. Bailey BA, Strem MD, Wood D. Trichoderma species form endophytic associations within Theobroma cacao trichomes. Mycological Research. 2009;113: 1365–1376. doi:10.1016/j.mycres.2009.09.004

3. Chaverri P, Branco-Rocha F, Jaklitsch W, Gazis R, Degenkolb T, Samuels GJ. Systematics of the Trichoderma harzianum species complex and the re-identification of commercial biocontrol strains. Mycologia. 2015;107: 558–590. doi:10.3852/14-147

4. Chaverri P, Samuels GJ. Evolution of Habitat Preference and Nutrition Mode in a Cosmopolitan Fungal Genus with Evidence of Interkingdom Host Jumps and Major Shifts in Ecology. Evolution. 2013;67: 2823–2837. doi:10.1111/evo.12169

5. Hoyos-Carvajal L, Orduz S, Bissett J. Genetic and metabolic biodiversity of Trichoderma from Colombia and adjacent neotropic regions. Fungal Genetics and Biology. 2009;46: 615–631. doi:10.1016/j.fgb.2009.04.006

6. Barrera VA, Iannone L, Romero AI, Chaverri P. Expanding the Trichoderma harzianum species complex: Three new species from Argentine natural and cultivated ecosystems. Mycologia. 2021;0: 1–20. doi:10.1080/00275514.2021.1947641

7. Tan RX, Zou WX. Endophytes: a rich source of functional metabolites. Nat Prod Rep. 2001;18: 448–459. doi:10.1039/B100918O

8. Strobel GA. Endophytes as sources of bioactive products. Microbes and Infection. 2003;5: 535–544. doi:10.1016/S1286-4579(03)00073-X

9. Hanada RE, de Jorge Souza T, Pomella AWV, Hebbar KP, Pereira JO, Ismaiel A, et al. Trichoderma martiale sp. nov., a new endophyte from sapwood of Theobroma cacao with a potential for biological control. Mycological Research. 2008;112: 1335–1343. doi:10.1016/j.mycres.2008.06.022

10. Yu H, Zhang L, Li L, Zheng C, Guo L, Li W, et al. Recent developments and future prospects of antimicrobial metabolites produced by endophytes. Microbiological Research. 2010;165: 437–449. doi:10.1016/j.micres.2009.11.009

11. Krauss U, Hidalgo E, Bateman R, Adonijah V, Arroyo C, García J, et al. Improving the formulation and timing of application of endophytic biocontrol and chemical agents against frosty pod rot (Moniliophthora roreri) in cocoa (Theobroma cacao). Biological Control. 2010;54: 230–240. doi:10.1016/j.biocontrol.2010.05.011

12. Hanada RE, Pomella AWV, Costa HS, Bezerra JL, Loguercio LL, Pereira JO. Endophytic fungal diversity in Theobroma cacao (cacao) and T. grandiflorum (cupuaçu) trees and their potential for growth promotion and biocontrol of black-pod disease. Fungal Biology. 2010;114: 901–910. doi:10.1016/j.funbio.2010.08.006

13. Mousa WK, Raizada MN. The Diversity of Anti-Microbial Secondary Metabolites Produced by Fungal Endophytes: An Interdisciplinary Perspective. Front Microbiol. 2013;4. doi:10.3389/fmicb.2013.00065

14. Alvin A, Miller KI, Neilan BA. Exploring the potential of endophytes from medicinal plants as sources of antimycobacterial compounds. Microbiological Research. 2014;169: 483–495. doi:10.1016/j.micres.2013.12.009

15. Pujade-Renaud V, Déon M, Gazis R, Ribeiro S, Dessailly F, Granet F, et al. Endophytes from Wild Rubber Trees as Antagonists of the Pathogen Corynespora cassiicola. Phytopathology®. 2019;109: 1888–1899. doi:10.1094/PHYTO-03-19-0093-R

16. Martínez-Medina A, Del Mar Alguacil M, Pascual JA, Van Wees SCM. Phytohormone Profiles Induced by Trichoderma Isolates Correspond with Their Biocontrol and Plant Growth-Promoting Activity on Melon Plants. J Chem Ecol. 2014;40: 804–815. doi:10.1007/s10886-014-0478-1

17. Pusztahelyi T, Holb I, Pócsi I. Secondary metabolites in fungus-plant interactions. Frontiers in Plant Science. 2015;6. Available: https://www.frontiersin.org/articles/10.3389/fpls.2015.00573

18. Sarrocco S, Matarese F, Baroncelli R, Vannacci G, Seidl-Seiboth V, Kubicek CP, et al. The Constitutive Endopolygalacturonase TvPG2 Regulates the Induction of Plant Systemic Resistance by Trichoderma virens. Phytopathology®. 2017;107: 537–544. doi:10.1094/PHYTO-03-16-0139-R

19. Rai S, Solanki MK, Solanki AC, Surapathrudu K. Biocontrol Potential of Trichoderma spp.: Current Understandings and Future Outlooks on Molecular Techniques. In: Ansari RA, Mahmood I, editors. Plant Health Under Biotic Stress: Volume 2: Microbial Interactions. Singapore: Springer; 2019. pp. 129–160. doi:10.1007/978-981-13-6040-4_7

20. Jaroszuk-Ściseł J, Tyśkiewicz R, Nowak A, Ozimek E, Majewska M, Hanaka A, et al. Phytohormones (Auxin, Gibberellin) and ACC Deaminase In Vitro Synthesized by the Mycoparasitic Trichoderma DEMTkZ3A0 Strain and Changes in the Level of Auxin and Plant Resistance Markers in Wheat Seedlings Inoculated with this Strain Conidia. International Journal of Molecular Sciences. 2019;20: 4923. doi:10.3390/ijms20194923

21. Singh BN, Dwivedi P, Sarma BK, Singh GS, Singh HB. A novel function of N-signaling in plants with special reference to Trichoderma interaction influencing plant growth, nitrogen use efficiency, and cross talk with plant hormones. 3 Biotech. 2019;9: 109. doi:10.1007/s13205-019-1638-3

22. Harman GE, Kubicek CP, editors. Trichoderma And Gliocladium, Volume 2: Enzymes, Biological Control and commercial applications. London: CRC Press; 2014. doi:10.1201/9781482267945

23. Gazis R, Chaverri P. Wild trees in the Amazon basin harbor a great diversity of beneficial endosymbiotic fungi: is this evidence of protective mutualism? Fungal Ecology. 2015;17: 18–29. doi:10.1016/j.funeco.2015.04.001

24. Chong J, Poutaraud A, Hugueney P. Metabolism and roles of stilbenes in plants. Plant Science. 2009;177: 143–155. doi:10.1016/j.plantsci.2009.05.012

25. Jeandet P, Delaunois B, Conreux A, Donnez D, Nuzzo V, Cordelier S, et al. Biosynthesis, metabolism, molecular engineering, and biological functions of stilbene phytoalexins in plants. BioFactors. 2010;36: 331–341. doi:10.1002/biof.108

26. Franco MEE, Wisecaver JH, Arnold AE, Ju Y-M, Slot JC, Ahrendt S, et al. Ecological generalism drives hyperdiversity of secondary metabolite gene clusters in xylarialean endophytes. New Phytologist. 2022;233: 1317–1330. doi:10.1111/nph.17873

27. Hammerbacher A, Schmidt A, Wadke N, Wright LP, Schneider B, Bohlmann J, et al. A Common Fungal Associate of the Spruce Bark Beetle Metabolizes the Stilbene Defenses of Norway Spruce. Plant Physiology. 2013;162: 1324–1336. doi:10.1104/pp.113.218610

28. Wang Y, Lim L, Madilao L, Lah L, Bohlmann J, Breuil C. Gene Discovery for Enzymes Involved in Limonene Modification or Utilization by the Mountain Pine Beetle-Associated Pathogen Grosmannia clavigera. Applied and Environmental Microbiology. 2014;80: 4566–4576. doi:10.1128/AEM.00670-14

29. Wadke N, Kandasamy D, Vogel H, Lah L, Wingfield BD, Paetz C, et al. The Bark-Beetle-Associated Fungus, Endoconidiophora polonica, Utilizes the Phenolic Defense Compounds of Its Host as a Carbon Source. Plant Physiology. 2016;171: 914–931. doi:10.1104/pp.15.01916

30. Gluck-Thaler E, Slot JC. Specialized plant biochemistry drives gene clustering in fungi. The ISME Journal. 2018;12: 1694–1705. doi:10.1038/s41396-018-0075-3

31. Zhao T, Kandasamy D, Krokene P, Chen J, Gershenzon J, Hammerbacher A. Fungal associates of the tree-killing bark beetle, Ips typographus, vary in virulence, ability to degrade conifer phenolics and influence bark beetle tunneling behavior. Fungal Ecology. 2019;38: 71–79. doi:10.1016/j.funeco.2018.06.003

32. Wisecaver JH, Slot JC, Rokas A. The evolution of fungal metabolic pathways. PLoS Genet. 2014;10: e1004816–e1004816. doi:10.1371/journal.pgen.1004816

33. Adamek M, Spohn M, Stegmann E, Ziemert N. Mining Bacterial Genomes for Secondary Metabolite Gene Clusters. In: Sass P, editor. Antibiotics: Methods and Protocols. New York, NY: Springer; 2017. pp. 23–47. doi:10.1007/978-1-4939-6634-9_2

34. Slot JC. Chapter Four - Fungal Gene Cluster Diversity and Evolution. In: Townsend JP, Wang Z, editors. Advances in Genetics. Academic Press; 2017. pp. 141–178. doi:10.1016/bs.adgen.2017.09.005

35. Rokas A, Wisecaver JH, Lind AL. The birth, evolution and death of metabolic gene clusters in fungi. Nature Reviews Microbiology. 2018;16: 731–744.

36. Tran PN, Yen M-R, Chiang C-Y, Lin H-C, Chen P-Y. Detecting and prioritizing biosynthetic gene clusters for bioactive compounds in bacteria and fungi. Appl Microbiol Biotechnol. 2019;103: 3277–3287. doi:10.1007/s00253-019-09708-z

37. Slot JC, Hallstrom KN, Matheny PB, Hibbett DS. Diversification of NRT2 and the Origin of Its Fungal Homolog. Molecular Biology and Evolution. 2007;24: 1731–1743. doi:10.1093/molbev/msm098

38. Slot JC, Gluck-Thaler E. Metabolic gene clusters, fungal diversity, and the generation of accessory functions. Current Opinion in Genetics & Development. 2019;58–59: 17–24. doi:10.1016/j.gde.2019.07.006

39. Vargas WA, Mukherjee PK, Laughlin D, Wiest A, Moran-Diez ME, Kenerley CMY 2014. Role of gliotoxin in the symbiotic and pathogenic interactions of Trichoderma virens. Microbiology. 2014;160: 2319–2330. doi:10.1099/mic.0.079210-0

40. Tomah AA, Abd Alamer IS, Li B, Zhang J-Z. A new species of Trichoderma and gliotoxin role: A new observation in enhancing biocontrol potential of T. virens against Phytophthora capsici on chili pepper. Biological Control. 2020;145: 104261. doi:10.1016/j.biocontrol.2020.104261

41. Elmore MH, McGary KL, Wisecaver JH, Slot JC, Geiser DM, Sink S, et al. Clustering of Two Genes Putatively Involved in Cyanate Detoxification Evolved Recently and Independently in Multiple Fungal Lineages. Genome Biology and Evolution. 2015;7: 789–800. doi:10.1093/gbe/evv025

42. Glenn AE, Davis CB, Gao M, Gold SE, Mitchell TR, Proctor RH, et al. Two Horizontally Transferred Xenobiotic Resistance Gene Clusters Associated with Detoxification of Benzoxazolinones by Fusarium Species. PLOS ONE. 2016;11: e0147486. doi:10.1371/journal.pone.0147486

43. Gluck-Thaler E, Vijayakumar V, Slot JC. Fungal adaptation to plant defences through convergent assembly of metabolic modules. Mol Ecol. 2018; mec.14943. doi:10.1111/mec.14943

44. Cohen-Kupiec R, Broglie KE, Friesem D, Broglie RM, Chet I. Molecular characterization of a novel β-1,3-exoglucanase related to mycoparasitism of Trichoderma harzianum. Gene. 1999;226: 147–154. doi:10.1016/S0378-1119(98)00583-6

45. Viterbo A, Ramot O, Chernin L, Chet I. Significance of lytic enzymes from Trichoderma spp. in the biocontrol of fungal plant pathogens. Antonie Van Leeuwenhoek. 2002;81: 549–556. doi:10.1023/A:1020553421740

46. Iqbal M, Dubey M, Gudmundsson M, Viketoft M, Jensen DF, Karlsson M. Comparative evolutionary histories of fungal proteases reveal gene gains in the mycoparasitic and nematode-parasitic fungus Clonostachys rosea. BMC Evol Biol. 2018;18: 171. doi:10.1186/s12862-018-1291-1

47. Duo-Chuan L. Review of Fungal Chitinases. Mycopathologia. 2006;161: 345–360. doi:10.1007/s11046-006-0024-y

48. Seidl V. Chitinases of filamentous fungi: a large group of diverse proteins with multiple physiological functions. Fungal Biology Reviews. 2008;22: 36–42. doi:10.1016/j.fbr.2008.03.002

49. Karlsson M, Stenlid J. Comparative Evolutionary Histories of the Fungal Chitinase Gene Family Reveal Non-Random Size Expansions and Contractions due to Adaptive Natural Selection. Evol Bioinform Online. 2008;4: EBO.S604. doi:10.4137/EBO.S604

50. Kubicek CP, Herrera-Estrella A, Seidl-Seiboth V, Martinez DA, Druzhinina IS, Thon M, et al. Comparative genome sequence analysis underscores mycoparasitism as the ancestral life style of Trichoderma. Genome Biol. 2011;12: R40. doi:10.1186/gb-2011-12-4-r40

51. Ihrmark K, Asmail N, Ubhayasekera W, Melin P, Stenlid J, Karlsson M. Comparative Molecular Evolution of Trichoderma Chitinases in Response to Mycoparasitic Interactions. Evol Bioinform Online. 2010;6: EBO.S4198. doi:10.4137/EBO.S4198

52. Schardl CL, Young CA, Hesse U, Amyotte SG, Andreeva K, Calie PJ, et al. Plant- Symbiotic Fungi as Chemical Engineers: Multi-Genome Analysis of the Clavicipitaceae Reveals Dynamics of Alkaloid Loci. PLOS Genetics. 2013;9: e1003323. doi:10.1371/journal.pgen.1003323

53. Aranda-Martinez A, Lenfant N, Escudero N, Zavala-Gonzalez EA, Henrissat B, Lopez-Llorca LV. CAZyme content of Pochonia chlamydosporia reflects that chitin and chitosan modification are involved in nematode parasitism. Environmental Microbiology. 2016;18: 4200–4215. doi:10.1111/1462-2920.13544

54. Lopez-Moya F, Escudero N, Lopez-Llorca LV. Pochonia chlamydosporia: Multitrophic Lifestyles Explained by a Versatile Genome. In: Manzanilla-López RH, Lopez-Llorca LV, editors. Perspectives in Sustainable Nematode Management Through Pochonia chlamydosporia Applications for Root and Rhizosphere Health. Cham: Springer International Publishing; 2017. pp. 197–207. doi:10.1007/978-3-319-59224-4_10

55. Carbone I, Jakobek JL, Ramirez-Prado JH, Horn BW. Recombination, balancing selection and adaptive evolution in the aflatoxin gene cluster of Aspergillus parasiticus. Molecular Ecology. 2007;16: 4401–4417. doi:10.1111/j.1365-294X.2007.03464.x

56. Gluck-Thaler E, Slot JC. Dimensions of Horizontal Gene Transfer in Eukaryotic Microbial Pathogens. PLOS Pathogens. 2015;11: e1005156. doi:10.1371/journal.ppat.1005156

57. Patron NJ, Waller RF, Cozijnsen AJ, Straney DC, Gardiner DM, Nierman WC, et al. Origin and distribution of epipolythiodioxopiperazine (ETP) gene clusters in filamentous ascomycetes. BMC Evolutionary Biology. 2007;7: 174. doi:10.1186/1471-2148-7-174

58. Matsuzawa T, Fujita Y, Tanaka N, Tohda H, Itadani A, Takegawa K. New insights into galactose metabolism by Schizosaccharomyces pombe: Isolation and characterization of a galactose-assimilating mutant. Journal of Bioscience and Bioengineering. 2011;111: 158–166. doi:10.1016/j.jbiosc.2010.10.007

59. Proctor RH, McCormick SP, Kim H-S, Cardoza RE, Stanley AM, Lindo L, et al. Evolution of structural diversity of trichothecenes, a family of toxins produced by plant pathogenic and entomopathogenic fungi. PLOS Pathogens. 2018;14: e1006946. doi:10.1371/journal.ppat.1006946

60. Druzhinina I, Kubicek CP. Species concepts and biodiversity in Trichoderma and Hypocrea: from aggregate species to species clusters? J Zhejiang Univ Sci B. 2005;6: 100–112. doi:10.1631/jzus.2005.B0100

61. Jaklitsch WM. European species of Hypocrea Part I. The green-spored species. Studies in Mycology. 2009;63: 1–91. doi:10.3114/sim.2009.63.01

62. Atanasova L, Crom SL, Gruber S, Coulpier F, Seidl-Seiboth V, Kubicek CP, et al. Comparative transcriptomics reveals different strategies of Trichoderma mycoparasitism. BMC Genomics. 2013;14: 121. doi:10.1186/1471-2164-14-121

63. Steindorff AS, Ramada MHS, Coelho ASG, Miller RNG, Pappas GJ, Ulhoa CJ, et al. Identification of mycoparasitism-related genes against the phytopathogen Sclerotinia sclerotiorum through transcriptome and expression profile analysis in Trichoderma harzianum. BMC Genomics. 2014;15: 204. doi:10.1186/1471-2164-15-204

64. Morán-Diez ME, Carrero-Carrón I, Rubio MB, Jiménez-Díaz RM, Monte E, Hermosa R. Transcriptomic Analysis of Trichoderma atroviride Overgrowing Plant-Wilting Verticillium dahliae Reveals the Role of a New M14 Metallocarboxypeptidase CPA1 in Biocontrol. Frontiers in Microbiology. 2019;10: 1120. doi:10.3389/fmicb.2019.01120

65. Goughenour K. Histoplasma capsulatum: Drugs and Sugars. The Ohio State University. 2020. Available: http://rave.ohiolink.edu/etdc/view?acc_num=osu1584377509624302

66. Kautsar SA, Blin K, Shaw S, Navarro-Muñoz JC, Terlouw BR, van der Hooft JJJ, et al. MIBiG 2.0: a repository for biosynthetic gene clusters of known function. Nucleic Acids Research. 2020;48: D454–D458. doi:10.1093/nar/gkz882

67. Sultana A, Kallio P, Jansson A, Wang J-S, Niemi J, Mäntsälä P, et al. Structure of the polyketide cyclase SnoaL reveals a novel mechanism for enzymatic aldol condensation. EMBO J. 2004;23: 1911–1921. doi:10.1038/sj.emboj.7600201

68. Shenouda M, Cox R. Molecular methods unravel the biosynthetic potential of Trichoderma species. RSC Advances. 2021;11: 3622–3635. doi:10.1039/D0RA09627J

69. Evans HC, Holmes KA, Thomas SE. Endophytes and mycoparasites associated with an indigenous forest tree, Theobroma gileri, in Ecuador and a preliminary assessment of their potential as biocontrol agents of cocoa diseases. Mycol Progress. 2003;2: 149–160. doi:10.1007/s11557-006-0053-4

70. Verma VC, Gond SK, Kumar A, Kharwar RN, Strobel G. The Endophytic Mycoflora of Bark, Leaf, and Stem Tissues of Azadirachta indica A. Juss (Neem) from Varanasi (India). Microb Ecol. 2007;54: 119–125. doi:10.1007/s00248-006-9179-9

71. Zamioudis C, Pieterse CMJ. Modulation of Host Immunity by Beneficial Microbes. MPMI. 2012;25: 139–150. doi:10.1094/MPMI-06-11-0179

72. Xu X-H, Wang C, Li S-X, Su Z-Z, Zhou H-N, Mao L-J, et al. Friend or foe: differential responses of rice to invasion by mutualistic or pathogenic fungi revealed by RNAseq and metabolite profiling. Scientific Reports. 2015;5: 13624. doi:10.1038/srep13624

73. Kapulnik Y, Volpin H, Itzhaki H, Ganon D, Galili S, David R, et al. Suppression of defence responses in mycorrhizal alfalfa and tobacco roots. New Phytologist. 1996;133: 59–64. doi:10.1111/j.1469-8137.1996.tb04341.x

74. Liu J, Blaylock LA, Endre G, Cho J, Town CD, VandenBosch KA, et al. Transcript Profiling Coupled with Spatial Expression Analyses Reveals Genes Involved in Distinct Developmental Stages of an Arbuscular Mycorrhizal Symbiosis [W]. The Plant Cell. 2003;15: 2106–2123. doi:10.1105/tpc.014183

75. Paszkowski U. Mutualism and parasitism: the yin and yang of plant symbioses. Current Opinion in Plant Biology. 2006;9: 364–370. doi:10.1016/j.pbi.2006.05.008

76. Heller G, Adomas A, Li G, Osborne J, van Zyl L, Sederoff R, et al. Transcriptional analysis of Pinus sylvestris roots challenged with the ectomycorrhizal fungus Laccaria bicolor. BMC Plant Biology. 2008;8: 19. doi:10.1186/1471-2229-8-19

77. Mäkelä MR, Marinović M, Nousiainen P, Liwanag AJM, Benoit I, Sipilä J, et al. Chapter Two - Aromatic Metabolism of Filamentous Fungi in Relation to the Presence of Aromatic Compounds in Plant Biomass. In: Sariaslani S, Gadd GM, editors. Advances in Applied Microbiology. Academic Press; 2015. pp. 63–137. doi:10.1016/bs.aambs.2014.12.001

78. Strobel G, Daisy B. Bioprospecting for Microbial Endophytes and Their Natural Products. Microbiology and Molecular Biology Reviews. 2003;67: 491–502. doi:10.1128/MMBR.67.4.491-502.2003

79. Shalaby S, Horwitz BA. Plant phenolic compounds and oxidative stress: integrated signals in fungal–plant interactions. Curr Genet. 2015;61: 347–357. doi:10.1007/s00294-014-0458-6

80. Hartley SE, Eschen R, Horwood JM, Gange AC, Hill EM. Infection by a foliar endophyte elicits novel arabidopside-based plant defence reactions in its host, Cirsium arvense. New Phytologist. 2015;205: 816–827. doi:10.1111/nph.13067

81. Ormeño E, Baldy V, Ballini C, Larcheveque MG-, Perissol C, Fernandez C. Effects of environmental factors and leaf chemistry on leaf litter colonization by fungi in a Mediterranean shrubland. Pedobiologia. 2006;50: 1. doi:10.1016/j.pedobi.2005.07.005

82. Chomel M, Fernandez C, Bousquet-Mélou A, Gers C, Monnier Y, Santonja M, et al. Secondary metabolites of Pinus halepensis alter decomposer organisms and litter decomposition during afforestation of abandoned agricultural zones. Journal of Ecology. 2014;102: 411–424. doi:10.1111/1365-2745.12205

83. Greene GH, McGary KL, Rokas A, Slot JC. Ecology Drives the Distribution of Specialized Tyrosine Metabolism Modules in Fungi. Genome Biol Evol. 2014;6: 121–132. doi:10.1093/gbe/evt208

84. Lattanzio V, Lattanzio V, Cardinali A. Role of phenolics in the resistance mechanisms of plants against fungal pathogens and insects. Phytochemistry: Advances in Research. 2006;661: 23–67.

85. Bors W, Michel C. Chemistry of the Antioxidant Effect of Polyphenols. Annals of the New York Academy of Sciences. 2002;957: 57–69. doi:10.1111/j.1749-6632.2002.tb02905.x

86. Vinale F, Marra R, Scala F, Ghisalberti E I., Lorito M, Sivasithamparam K. Major secondary metabolites produced by two commercial Trichoderma strains active against different phytopathogens. Letters in Applied Microbiology. 2006;43: 143–148. doi:10.1111/j.1472-765X.2006.01939.x

87. Vinale F, Sivasithamparam K, Ghisalberti EL, Marra R, Barbetti MJ, Li H, et al. A novel role for Trichoderma secondary metabolites in the interactions with plants. Physiological and Molecular Plant Pathology. 2008;72: 80–86. doi:10.1016/j.pmpp.2008.05.005

88. Cai F, Yu G, Wang P, Wei Z, Fu L, Shen Q, et al. Harzianolide, a novel plant growth regulator and systemic resistance elicitor from Trichoderma harzianum. Plant Physiology and Biochemistry. 2013;73: 106–113. doi:10.1016/j.plaphy.2013.08.011

89. Vinale F, Flematti G, Sivasithamparam K, Lorito M, Marra R, Skelton BW, et al. Harzianic Acid, an Antifungal and Plant Growth Promoting Metabolite from Trichoderma harzianum. J Nat Prod. 2009;72: 2032–2035. doi:10.1021/np900548p

90. Claydon N, Allan M, Hanson JR, Avent AG. Antifungal alkyl pyrones of Trichoderma harzianum. Transactions of the British Mycological Society. 1987;88: 503–513. doi:10.1016/S0007-1536(87)80034-7

91. Parker SR, Cutler HG, Jacyno JM, Hill RA. Biological Activity of 6-Pentyl-2H-pyran-2-one and Its Analogs. J Agric Food Chem. 1997;45: 2774–2776. doi:10.1021/jf960681a

92. Barry SM, Challis GL. Recent advances in siderophore biosynthesis. Current Opinion in Chemical Biology. 2009;13: 205–215. doi:10.1016/j.cbpa.2009.03.008

93. Arnold AE, Mejía LC, Kyllo D, Rojas EI, Maynard Z, Robbins N, et al. Fungal endophytes limit pathogen damage in a tropical tree. PNAS. 2003;100: 15649–15654. doi:10.1073/pnas.2533483100

94. Keswani C, Mishra S, Sarma BK, Singh SP, Singh HB. Unraveling the efficient applications of secondary metabolites of various Trichoderma spp. Appl Microbiol Biotechnol. 2014;98: 533–544. doi:10.1007/s00253-013-5344-5

95. Keswani C, Bisen K, Chitara MK, Sarma BK, Singh HB. Exploring the Role of Secondary Metabolites of Trichoderma in Tripartite Interaction with Plant and Pathogens. In: Singh JS, Seneviratne G, editors. Agro-Environmental Sustainability: Volume 1: Managing Crop Health. Cham: Springer International Publishing; 2017. pp. 63–79. doi:10.1007/978-3-319-49724-2_4

96. Vinale F, Nigro M, Sivasithamparam K, Flematti G, Ghisalberti EL, Ruocco M, et al. Harzianic acid: a novel siderophore from Trichoderma harzianum. FEMS Microbiology Letters. 2013;347: 123–129. doi:10.1111/1574-6968.12231

97. Scherlach K, Hertweck C. Triggering cryptic natural product biosynthesis in microorganisms. Org Biomol Chem. 2009;7: 1753–1760. doi:10.1039/b821578b

98. Kusari S, Zühlke S, Spiteller M. An Endophytic Fungus from Camptotheca acuminata That Produces Camptothecin and Analogues. J Nat Prod. 2009;72: 2–7. doi:10.1021/np800455b

99. Kusari S, Zühlke S, Spiteller M. Effect of Artificial Reconstitution of the Interaction between the Plant Camptotheca acuminata and the Fungal Endophyte Fusarium solani on Camptothecin Biosynthesis. J Nat Prod. 2011;74: 764–775. doi:10.1021/np1008398

100. Demyttenaere J, De Kimpe N. Biotransformation of terpenes by fungi: Study of the pathways involved. Journal of Molecular Catalysis B: Enzymatic. 2001;11: 265–270. doi:10.1016/S1381-1177(00)00040-0

101. Carroll GC. Beyond Pest Deterrence—Alternative Strategies and Hidden Costs of Endophytic Mutualisms in Vascular Plants. In: Andrews JH, Hirano SS, editors. Microbial Ecology of Leaves. New York, NY: Springer; 1991. pp. 358–375. doi:10.1007/978-1-4612-3168-4_18

102. Stierle A, Strobel G, Stierle D. Taxol and Taxane Production by Taxomyces andreanae, an Endophytic Fungus of Pacific Yew. Science. 1993;260: 214–216. doi:10.1126/science.8097061

103. Eyberger AL, Dondapati R, Porter JR. Endophyte Fungal Isolates from Podophyllum peltatum Produce Podophyllotoxin. J Nat Prod. 2006;69: 1121–1124. doi:10.1021/np060174f

104. Leuchtmann A, Schmidt D, Bush LP. Different Levels of Protective Alkaloids in Grasses with Stroma-forming and Seed-transmitted Epichloë/Neotyphodium Endophytes. J Chem Ecol. 2000;26: 1025–1036. doi:10.1023/A:1005489032025

105. Panaccione DG, Beaulieu WT, Cook D. Bioactive alkaloids in vertically transmitted fungal endophytes. Functional Ecology. 2014;28: 299–314. doi:10.1111/1365-2435.12076

106. Almassi F, Ghisalberti EL, Narbey MJ, Sivasithamparam K. New antibiotics from strains of Trichoderma harzianum. Journal of natural products (USA). 1991 [cited 2 Dec 2022]. Available: https://scholar.google.com/scholar_lookup?title=New+antibiotics+from+strains+of+Trichoderma+harzianum&author=Almassi%2C+F.+%28The+University+of+Western+Australia%2C+Nedlands%2C+Australia%29&publication_year=1991

107. Dickinson JM, Hanson JR, Hitchcock PB, Claydon N. Structure and biosynthesis of harzianopyridone, an antifungal metabolite of Trichoderma harzianum. J Chem Soc, Perkin Trans 1. 1989; 1885–1887. doi:10.1039/P19890001885

108. Fuchs B, Krauss J. Can Epichloë endophytes enhance direct and indirect plant defence? Fungal Ecology. 2019;38: 98–103. doi:10.1016/j.funeco.2018.07.002

109. Koide K, Osono T, Takeda H. Colonization and lignin decomposition of Camellia japonica leaf litter by endophytic fungi. Mycoscience. 2005;46: 280–286. doi:10.1007/S10267-005-0247-7

110. Promputtha I, Lumyong S, Dhanasekaran V, McKenzie EHC, Hyde KD, Jeewon R. A Phylogenetic Evaluation of Whether Endophytes Become Saprotrophs at Host Senescence. Microb Ecol. 2007;53: 579–590. doi:10.1007/s00248-006-9117-x

111. Parfitt D, Hunt J, Dockrell D, Rogers HJ, Boddy L. Do all trees carry the seeds of their own destruction? PCR reveals numerous wood decay fungi latently present in sapwood of a wide range of angiosperm trees. Fungal Ecology. 2010;3: 338–346. doi:10.1016/j.funeco.2010.02.001

112. U’Ren JM, Arnold AE. Diversity, taxonomic composition, and functional aspects of fungal communities in living, senesced, and fallen leaves at five sites across North America. PeerJ. 2016;4: e2768. doi:10.7717/peerj.2768

113. Chen K-H, Liao H-L, Arnold AE, Bonito G, Lutzoni F. RNA-based analyses reveal fungal communities structured by a senescence gradient in the moss Dicranum scoparium and the presence of putative multi-trophic fungi. New Phytologist. 2018;218: 1597–1611. doi:10.1111/nph.15092

114. U’Ren JM, Miadlikowska J, Zimmerman NB, Lutzoni F, Stajich JE, Arnold AE. Contributions of North American endophytes to the phylogeny, ecology, and taxonomy of Xylariaceae (Sordariomycetes, Ascomycota). Molecular Phylogenetics and Evolution. 2016;98: 210–232. doi:10.1016/j.ympev.2016.02.010

115. Vicente I, Baroncelli R, Hermosa R, Monte E, Vannacci G, Sarrocco S. Role and genetic basis of specialised secondary metabolites in Trichoderma ecophysiology. Fungal Biology Reviews. 2022;39: 83–99. doi:10.1016/j.fbr.2021.12.004

116. Herrera-Estrella A. Chapter 33 - Genome-Wide Approaches toward Understanding Mycotrophic Trichoderma Species. In: Gupta VK, Schmoll M, Herrera-Estrella A, Upadhyay RS, Druzhinina I, Tuohy MG, editors. Biotechnology and Biology of Trichoderma. Amsterdam: Elsevier; 2014. pp. 455–464. doi:10.1016/B978-0-444-59576-8.00033-3

117. Osono T, Hirose D, Fujimaki R. Fungal colonization as affected by litter depth and decomposition stage of needle litter. Soil Biology and Biochemistry. 2006;38: 2743–2752. doi:10.1016/j.soilbio.2006.04.028

118. Osono T, Takeda H. Organic chemical and nutrient dynamics in decomposing beech leaf litter in relation to fungal ingrowth and succession during 3-year decomposition processes in a cool temperate deciduous forest in Japan. Ecological Research. 2001;16: 649–670. doi:10.1046/j.1440-1703.2001.00426.x

119. Osono T. Colonization and succession of fungi during decomposition of Swida controversa leaf litter. Mycologia. 2005;97: 589–597. doi:10.1080/15572536.2006.11832789

120. Frankland JC. Fungal succession — unravelling the unpredictable. Mycological Research. 1998;102: 1–15. doi:10.1017/S0953756297005364

121. Osono T. Phyllosphere fungi on leaf litter of Fagus crenata: occurrence, colonization, and succession. Can J Bot. 2002;80: 460–469. doi:10.1139/b02-028

122. del Cerro C, Erickson E, Dong T, Wong AR, Eder EK, Purvine SO, et al. Intracellular pathways for lignin catabolism in white-rot fungi. Proceedings of the National Academy of Sciences. 2021;118: e2017381118. doi:10.1073/pnas.2017381118

123. Suryanarayanan TS, Thirunavukkarasu N, Govindarajulu MB, Sasse F, Jansen R, Murali TS. Fungal endophytes and bioprospecting. Fungal Biology Reviews. 2009;23: 9–19. doi:10.1016/j.fbr.2009.07.001

124. Kusari S, Pandey S, Spiteller M. Untapped mutualistic paradigms linking host plant and endophytic fungal production of similar bioactive secondary metabolites - ScienceDirect. Phytochemistry. 2013;91: 81–87.

125. Liu Y, Yang Q. Cloning and heterologous expression of aspartic protease SA76 related to biocontrol in Trichoderma harzianum. FEMS Microbiology Letters. 2007;277: 173–181. doi:10.1111/j.1574-6968.2007.00952.x

126. Viterbo A, Harel M, Chet I. Isolation of two aspartyl proteases from Trichoderma asperellum expressed during colonization of cucumber roots⋆. FEMS Microbiology Letters. 2004;238: 151–158. doi:10.1111/j.1574-6968.2004.tb09750.x

127. Lahrmann U, Zuccaro A. Opprimo ergo sum—Evasion and Suppression in the Root Endophytic Fungus Piriformospora indica. MPMI. 2012;25: 727–737. doi:10.1094/MPMI-11-11-0291

128. Bruce A, Srinivasan U, Staines HJ, Highley TL. Chitinase and laminarinase production in liquid culture by Trichoderma spp. and their role in biocontrol of wood decay fungi. International Biodeterioration & Biodegradation. 1995;35: 337–353. doi:10.1016/0964-8305(95)00047-3

129. Silva LG, Camargo RC, Mascarin GM, Nunes PS de O, Dunlap C, Bettiol W. Dual functionality of Trichoderma: Biocontrol of Sclerotinia sclerotiorum and biostimulant of cotton plants. Frontiers in Plant Science. 2022;13. Available: https://www.frontiersin.org/articles/10.3389/fpls.2022.983127

130. Latz MAC, Jensen B, Collinge DB, Jørgensen HJL. Endophytic fungi as biocontrol agents: elucidating mechanisms in disease suppression. Plant Ecology & Diversity. 2018;11: 555–567. doi:10.1080/17550874.2018.1534146

131. Sieber CMK, Lee W, Wong P, Münsterkötter M, Mewes H-W, Schmeitzl C, et al. The Fusarium graminearum Genome Reveals More Secondary Metabolite Gene Clusters and Hints of Horizontal Gene Transfer. PLOS ONE. 2014;9: e110311. doi:10.1371/journal.pone.0110311

132. Zhang Q, Chen X, Xu C, Zhao H, Zhang X, Zeng G, et al. Horizontal gene transfer allowed the emergence of broad host range entomopathogens. Proceedings of the National Academy of Sciences. 2019;116: 7982–7989. doi:10.1073/pnas.1816430116

133. Wang H, Sun S, Ge W, Zhao L, Hou B, Wang K, et al. Horizontal gene transfer of Fhb7 from fungus underlies Fusarium head blight resistance in wheat. Science. 2020;368: eaba5435. doi:10.1126/science.aba5435

134. Cardoza RE, McCormick SP, Izquierdo-Bueno I, Martínez-Reyes N, Lindo L, Brown DW, et al. Identification of polyketide synthase genes required for aspinolide biosynthesis in Trichoderma arundinaceum. Appl Microbiol Biotechnol. 2022 [cited 7 Oct 2022]. doi:10.1007/s00253-022-12182-9

135. Marcet-Houben M, Gabaldón T. Horizontal acquisition of toxic alkaloid synthesis in a clade of plant associated fungi. Fungal Genet Biol. 2016;86: 71–80. doi:10.1016/j.fgb.2015.12.006

136. Slot JC, Hibbett DS. Horizontal Transfer of a Nitrate Assimilation Gene Cluster and Ecological Transitions in Fungi: A Phylogenetic Study. PLOS ONE. 2007;2: e1097. doi:10.1371/journal.pone.0001097

137. Clay K, Cheplick GP. Effect of ergot alkaloids from fungal endophyte-infected grasses on fall armyworm (Spodoptera frugiperda). J Chem Ecol. 1989;15: 169–182. doi:10.1007/bf02027781

138. Guerre P. Ergot Alkaloids Produced by Endophytic Fungi of the Genus Epichloë. Toxins. 2015;7: 773–790. doi:10.3390/toxins7030773

139. Leadmon CE, Sampson JK, Maust MD, Macias AM, Rehner SA, Kasson MT, et al. Several Metarhizium Species Produce Ergot Alkaloids in a Condition-Specific Manner. Applied and Environmental Microbiology. 2020;86: e00373–20. doi:10.1128/AEM.00373-20

140. Johannesson H, Gustafsson M, Stenlid J. Local population structure of the wood decay ascomycete Daldinia loculata. Mycologia. 2001;93: 440–446. doi:10.1080/00275514.2001.12063176

141. Collado J, González A, Platas G, Stchigel AM, Guarro J, Peláez F. Monosporascus ibericus sp. nov., an endophytic ascomycete from plants on saline soils, with observations on the position of the genus based on sequence analysis of the 18S rDNA. Mycological Research. 2002;106: 118–127. doi:10.1017/S0953756201005172

142. Pažoutová S, Šrůtka P, Holuša J, Chudíčková M, Kolařík M. Diversity of xylariaceous symbionts in Xiphydria woodwasps: role of vector and a host tree. Fungal Ecology. 2010;3: 392–401. doi:10.1016/j.funeco.2010.07.002

143. Pandey A, Debdulal B. Daldinia bambusicola Ch4/11 an endophytic fungus producing volatile organic compounds having antimicrobial and olio chemical potential. Journal of Advances in Microbiology. 2014;1: 330–337.

144. Da Silva Cruz K, Cortez V G. Xylaria (Xylariaceae, Ascomycota) in the Parque Estadual de São Camilo, Paraná, Brazil. Acta Biol Par. 2015;44: 1–4.

145. Leman-Loubière C, Le Goff G, Debitus C, Ouazzani J. Sporochartines A–E, A New Family of Natural Products from the Marine Fungus Hypoxylon monticulosum Isolated from a Sphaerocladina Sponge. Frontiers in Marine Science. 2017;4. Available: https://www.frontiersin.org/articles/10.3389/fmars.2017.00399

146. Li W, Lee C, Bang SH, Ma JY, Kim S, Koh Y-S, et al. Isochromans and Related Constituents from the Endophytic Fungus Annulohypoxylon truncatum of Zizania caduciflora and Their Anti-Inflammatory Effects. J Nat Prod. 2017;80: 205–209. doi:10.1021/acs.jnatprod.6b00698

147. Stoykov DY, Alvarado P. Daldinia vernicosa from the Eastern Forebalkan (Bulgaria). Phytologia Balcanica: International Journal of Balkan Flora and Vegetation. 2019;25: 153–155.

148. Mishra R, Kushveer JS, Khan MohdIK, Pagal S, Meena CK, Murali A, et al. 2,4-Di-Tert-Butylphenol Isolated From an Endophytic Fungus, Daldinia eschscholtzii, Reduces Virulence and Quorum Sensing in Pseudomonas aeruginosa. Frontiers in Microbiology. 2020;11. Available: https://www.frontiersin.org/articles/10.3389/fmicb.2020.01668

149. Kim JA, Jeon J, Park S-Y, Jeon MJ, Yeo J-H, Lee Y-H, et al. Draft Genome Sequence of Daldinia childiae JS-1345, an Endophytic Fungus Isolated from Stem Tissue of Korean Fir. Microbiology Resource Announcements. 2020;9: e01284–19. doi:10.1128/MRA.01284-19

150. Rogers JD, Ju Y-M, Lehmann J. Some Xylaria species on termite nests. Mycologia. 2005;97: 914–923. doi:10.1080/15572536.2006.11832783

151. Hu D, Li M. Three New Ergot Alkaloids from the Fruiting Bodies of Xylaria nigripes (Kl.) Sacc. Chemistry & Biodiversity. 2017;14: e1600173. doi:10.1002/cbdv.201600173

152. Tiwari P, Bae H. Horizontal Gene Transfer and Endophytes: An Implication for the Acquisition of Novel Traits. Plants. 2020;9: 305. doi:10.3390/plants9030305

153. Koch RA, Bach CE, Pirro S, Aime MC. Draft Genome Sequence of an Unusual Ectomycorrhizal Fungus, Pseudotulostoma volvatum. Microbiology Resource Announcements. 2022;11: e00801–21. doi:10.1128/mra.00801-21

154. Miller OK, Henkel TW, James TY, Miller SL. Pseudotulostoma, a remarkable new volvate genus in the Elaphomycetaceae from Guyana. Mycological Research. 2001;105: 1268–1272. doi:10.1017/S095375620100466X

155. Kjærbølling I, Vesth T, Frisvad JC, Nybo JL, Theobald S, Kildgaard S, et al. A comparative genomics study of 23 Aspergillus species from section Flavi. Nat Commun. 2020;11: 1106. doi:10.1038/s41467-019-14051-y

156. Bolger AM, Lohse M, Usadel B. Trimmomatic: a flexible trimmer for Illumina sequence data. Bioinformatics. 2014;30: 2114–2120. doi:10.1093/bioinformatics/btu170

157. Andrews S. FastQC: a quality control tool for high throughput sequence data. 2010. Available: http://www.bioinformatics.babraham.ac.uk/projects/fastqc/

158. Bankevich A, Nurk S, Antipov D, Gurevich AA, Dvorkin M, Kulikov AS, et al. SPAdes: A New Genome Assembly Algorithm and Its Applications to Single-Cell Sequencing. J Comput Biol. 2012;19: 455–477. doi:10.1089/cmb.2012.0021

159. Gurevich A, Saveliev V, Vyahhi N, Tesler G. QUAST: quality assessment tool for genome assemblies. Bioinformatics. 2013;29: 1072–1075. doi:10.1093/bioinformatics/btt086

160. Bushnell B. BBMap: A Fast, Accurate, Splice-Aware Aligner. Lawrence Berkeley National Lab. (LBNL), Berkeley, CA (United States); 2014 Mar. Report No.: LBNL-7065E. Available: https://www.osti.gov/biblio/1241166

161. Holt C, Yandell M. MAKER2: an annotation pipeline and genome-database management tool for second-generation genome projects. BMC Bioinformatics. 2011;12: 491. doi:10.1186/1471-2105-12-491

162. Smit AFA, Hubley R. RepeatModeler Open-1.0. 2008. Available: http://www.repeatmasker.org

163. Korf I. Gene finding in novel genomes. BMC Bioinformatics. 2004;5: 59. doi:10.1186/1471-2105-5-59

164. Stanke M, Diekhans M, Baertsch R, Haussler D. Using native and syntenically mapped cDNA alignments to improve de novo gene finding. Bioinformatics. 2008;24: 637–644. doi:10.1093/bioinformatics/btn013

165. Waterhouse RM, Seppey M, Simão FA, Manni M, Ioannidis P, Klioutchnikov G, et al. BUSCO Applications from Quality Assessments to Gene Prediction and Phylogenomics. Mol Biol Evol. 2018;35: 543–548. doi:10.1093/molbev/msx319

166. Besemer J, Borodovsky M. GeneMark: web software for gene finding in prokaryotes, eukaryotes and viruses. Nucleic Acids Res. 2005;33: W451–W454. doi:10.1093/nar/gki487

167. Emms DM, Kelly S. OrthoFinder: solving fundamental biases in whole genome comparisons dramatically improves orthogroup inference accuracy. Genome Biol. 2015;16: 157. doi:10.1186/s13059-015-0721-2

168. Dunne MP, Kelly S. OrthoFiller: utilising data from multiple species to improve the completeness of genome annotations. BMC Genomics. 2017;18: 390. doi:10.1186/s12864-017-3771-x

169. Katoh K, Standley DM. MAFFT Multiple Sequence Alignment Software Version 7: Improvements in Performance and Usability. Molecular Biology and Evolution. 2013;30: 772–780. doi:10.1093/molbev/mst010

170. Capella-Gutiérrez S, Silla-Martínez JM, Gabaldón T. trimAl: a tool for automated alignment trimming in large-scale phylogenetic analyses. Bioinformatics. 2009;25: 1972–1973. doi:10.1093/bioinformatics/btp348

171. Kozlov AM, Darriba D, Flouri T, Morel B, Stamatakis A. RAxML-NG: a fast, scalable and user-friendly tool for maximum likelihood phylogenetic inference. Bioinformatics. 2019;35: 4453–4455. doi:10.1093/bioinformatics/btz305

172. Minh BQ, Schmidt HA, Chernomor O, Schrempf D, Woodhams MD, von Haeseler A, et al. IQ-TREE 2: New Models and Efficient Methods for Phylogenetic Inference in the Genomic Era. Molecular Biology and Evolution. 2020;37: 1530–1534. doi:10.1093/molbev/msaa015

173. Blin K, Shaw S, Steinke K, Villebro R, Ziemert N, Lee SY, et al. antiSMASH 5.0: updates to the secondary metabolite genome mining pipeline. Nucleic Acids Res. 2019;47: W81–W87. doi:10.1093/nar/gkz310

174. Navarro-Muñoz JC, Selem-Mojica N, Mullowney MW, Kautsar SA, Tryon JH, Parkinson EI, et al. A computational framework to explore large-scale biosynthetic diversity. Nat Chem Biol. 2020;16: 60–68. doi:10.1038/s41589-019-0400-9

175. Eddy SR. A new generation of homology search tools based on probabilistic inference. Genome Inform. 2009;23: 205–211.

176. Price MN, Dehal PS, Arkin AP. FastTree 2 – Approximately Maximum-Likelihood Trees for Large Alignments. PLOS ONE. 2010;5: e9490. doi:10.1371/journal.pone.0009490

177. Marchler-Bauer A, Bo Y, Han L, He J, Lanczycki CJ, Lu S, et al. CDD/SPARCLE: functional classification of proteins via subfamily domain architectures. Nucleic Acids Research. 2017;45: D200–D203. doi:10.1093/nar/gkw1129

178. Fritz SA, Purvis A. Selectivity in Mammalian Extinction Risk and Threat Types: a New Measure of Phylogenetic Signal Strength in Binary Traits. Conservation Biology. 2010;24: 1042–1051. doi:10.1111/j.1523-1739.2010.01455.x

179. Orme D, Freckleton R, Thomas G, Petzoldt T, Fritz S, Isaac N, et al. caper: Comparative Analyses of Phylogenetics and Evolution in R. 2018. Available: <https://CRAN.R-project.org/package=caper>

180. Blomberg SP, Garland JR. T, Ives AR. Testing for Phylogenetic Signal in Comparative Data: Behavioral Traits Are More Labile. Evolution. 2003;57: 717– 745. doi:10.1111/j.0014-3820.2003.tb00285.x

181. Pagel M. Inferring evolutionary processes from phylogenies. Zoologica Scripta. 1997;26: 331–348. doi:10.1111/j.1463-6409.1997.tb00423.x

182. Pagel M. Inferring the historical patterns of biological evolution. Nature. 1999;401: 877–884. doi:10.1038/44766

183. Revell LJ. phytools: An R package for phylogenetic comparative biology (and other things). Methods in Ecology and Evolution. 2012;3: 217–223. doi:10.1111/j.2041-210X.2011.00169.x

184. Grigoriev IV, Nikitin R, Haridas S, Kuo A, Ohm R, Otillar R, et al. MycoCosm portal: gearing up for 1000 fungal genomes. Nucleic Acids Research. 2014;42: D699–D704. doi:10.1093/nar/gkt1183

185. Slater GSC, Birney E. Automated generation of heuristics for biological sequence comparison. BMC Bioinformatics. 2005;6: 31. doi:10.1186/1471-2105-6-31

186. Ye J, McGinnis S, Madden TL. BLAST: improvements for better sequence analysis. Nucleic Acids Research. 2006;34: W6–W9. doi:10.1093/nar/gkl164

187. Steinegger M, Söding J. MMseqs2 enables sensitive protein sequence searching for the analysis of massive data sets. Nat Biotechnol. 2017;35: 1026–1028. doi:10.1038/nbt.3988

188. Steenwyk JL, Link to external site this link will open in a new window, Buida TJ, Link to external site this link will open in a new window, Li Y, Link to external site this link will open in a new window, et al. ClipKIT: A multiple sequence alignment trimming software for accurate phylogenomic inference. PLoS Biology. 2020;18: e3001007. doi:10.1371/journal.pbio.3001007

189. James TY, Pelin A, Bonen L, Ahrendt S, Sain D, Corradi N, et al. Shared Signatures of Parasitism and Phylogenomics Unite Cryptomycota and Microsporidia. Current Biology. 2013;23: 1548–1553. doi:10.1016/j.cub.2013.06.057

190. Spatafora JW, Chang Y, Benny GL, Lazarus K, Smith ME, Berbee ML, et al. A phylum-level phylogenetic classification of zygomycete fungi based on genome-scale data. Mycologia. 2016;108: 1028–1046. doi:10.3852/16-042

191. Kalyaanamoorthy S, Minh BQ, Wong TKF, von Haeseler A, Jermiin LS. ModelFinder: fast model selection for accurate phylogenetic estimates. Nat Methods. 2017;14: 587–589. doi:10.1038/nmeth.4285

192. Gilchrist CLM, Chooi Y-H. clinker & clustermap.js: automatic generation of gene cluster comparison figures. Bioinformatics. 2021;37: 2473–2475. doi:10.1093/bioinformatics/btab007

193. Ohio Supercomputer Center. 1987. Available: http://osc.edu/ark:/19495/f5s1ph73

