## Supplemental Figures and Tables PDf for "Endophyte genomes support greater metabolic gene cluster diversity compared with non-endophytes in *Trichoderma*"

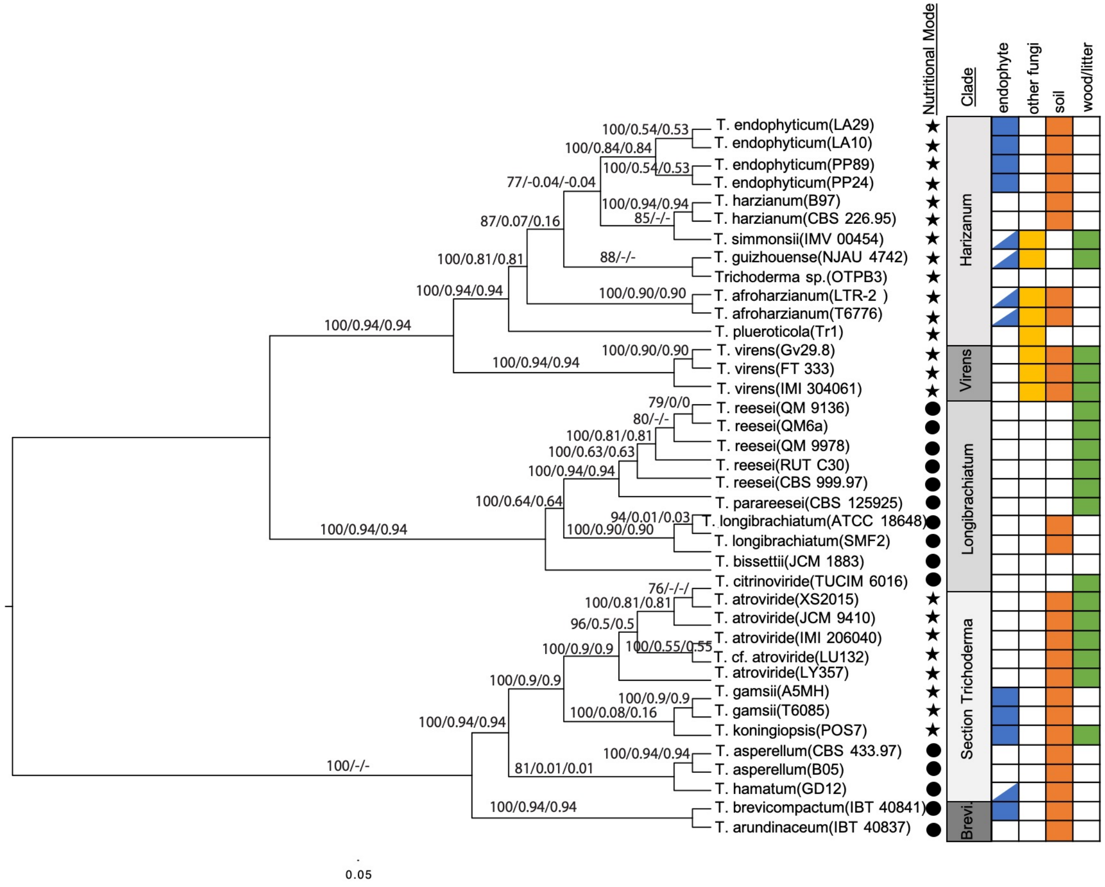

Nutritional Mode
Clade
endophyte
other fungi
soil
wood/litter
T. endophyticum(LA29)
T. endophyticum(LA10)
T. endophyticum(PP89)
T. endophyticum(PP24)
T. harzianum(B97)
T. harzianum(CBS 226.95)
T. simmonsii(IMV 00454)
T. guizhouense(NJAU 4742)
Trichoderma sp.(OTPB3)
T. afroharzianum(LTR-2 )
T. afroharzianum(T6776)
T. plueroticola(Tr1)
T. virens(Gv29.8)
T. virens(FT 333)
T. virens(IMI 304061)
T. reesei(QM 9136)
T. reesei(QM6a)
T. reesei(QM 9978)
T. reesei(RUT C30)
T. reesei(CBS 999.97)
T. parareesei(CBS 125925)
T. longibrachiatum(ATCC 18648)
T. longibrachiatum(SMF2)
T. bissettii(JCM 1883)
T. citrinoviride(TUCIM 6016)
T. atroviride(XS2015)
T. atroviride(JCM 9410)
T. atroviride(IMI 206040)
T. cf. atroviride(LU132)
T. atroviride(LY357)
T. gamsii(A5MH)
T. gamsii(T6085)
T. koningiopsis(POS7)
T. asperellum(CBS 433.97)
T. asperellum(B05)
T. hamatum(GD12)
T. brevicompactum(IBT 40841)
T. arundinaceum(IBT 40837)
Harizanum
Virens
Longibrachiatum
Section Trichoderma
Brevi
100/0.54/0.53
100/0.84/0.84
100/0.54/0.53
77/-0.04/-0.04
100/0.94/0.94
85/-/-
87/0.07/0.16
88/-/-
100/0.81/0.81
100/0.94/0.94
100/0.90/0.90
100/0.94/0.94
100/0.90/0.90
100/0.94/0.94
79/0/0
80/-/-
100/0.81/0.81
100/0.63/0.63
100/0.94/0.94
100/0.64/0.64
94/0.01/0.03
100/0.90/0.90
100/0.94/0.94
76/-/-/
100/0.81/0.81
96/0.5/0.5
100/0.55/0.55
100/0.9/0.9
100/0.9/0.9
100/0.9/0.9
100/0.08/0.16
100/0.94/0.94
100/0.94/0.94
81/0.01/0.01
100/-/-
100/0.94/0.94
0.05

**Fig S1. The dataset of 39 Trichoderma genomes contains 5 clades with different combinations of lifestyles and nutritional modes.**

This consensus phylogenomic tree was created from 154 SCO IQ-TREEs which had >98% BP, using 10,000 bootstraps. Values at nodes represent the IQ-TREE maximum likelihood BP, Internode Certainty (IC), and Internode Certainty All (ICA) (BP/IC/ICA). Internode Certainty (IC) and Internode Certainty All (ICA) values were obtained from the corresponding RAxML majority rule extended consensus tree. Species lifestyle is indicated by presence/absence of color-coordinated cell, nutritional mode is indicated by either a circle (saprotroph) or a star (mycotroph). Endophyte cells half-filled indicated a potential endophytic lifestyle for that species. Branch lengths are not proportional to sequence divergence.

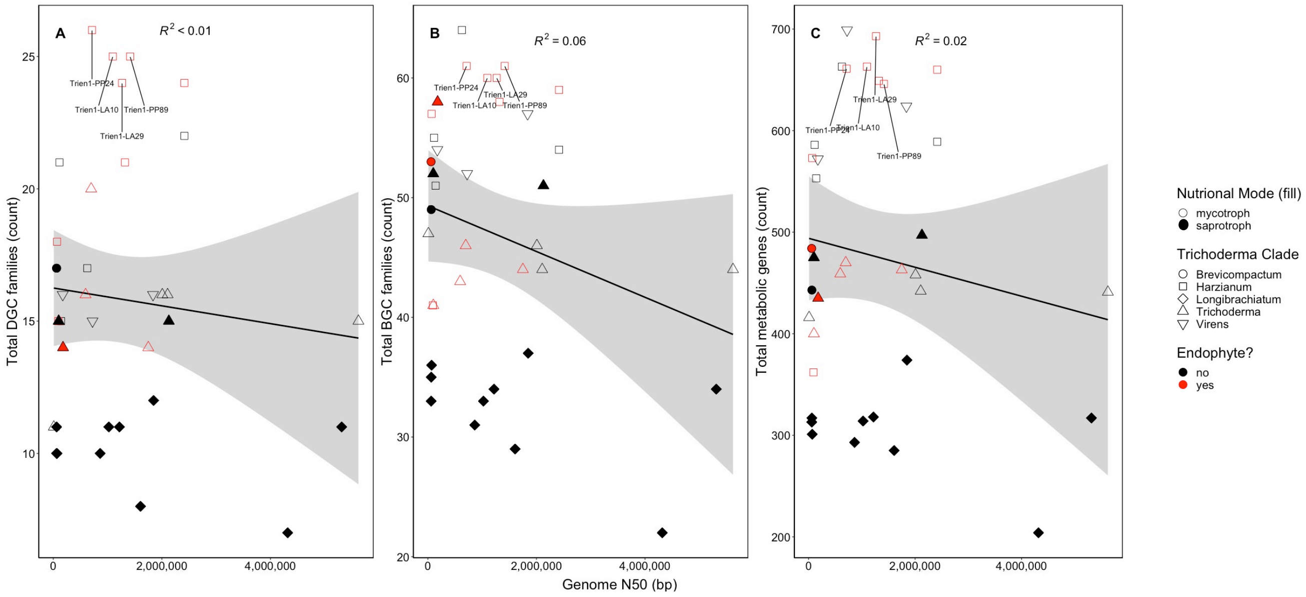


**Fig S2. Genome quality was not correlated with number of metabolic gene clusters identified.**
Genome quality is not correlated with the number of (A) DGC families (Pearson's, p > 0.05) (B) BGC families (Pearson's, p > 0.05), and (C) number of metabolic genes (Pearson's, p > 0.05) identified in the *Trichoderma* genomes. Each point represents a *Trichoderma* isolate. Point fill indicates nutritional mode, point shape indicates the clade to which the isolate belongs, and color indicates whether the *Trichoderma* species is recorded as having an endophytic lifestyle. Gray shaded areas indicate the 95% confidence interval.

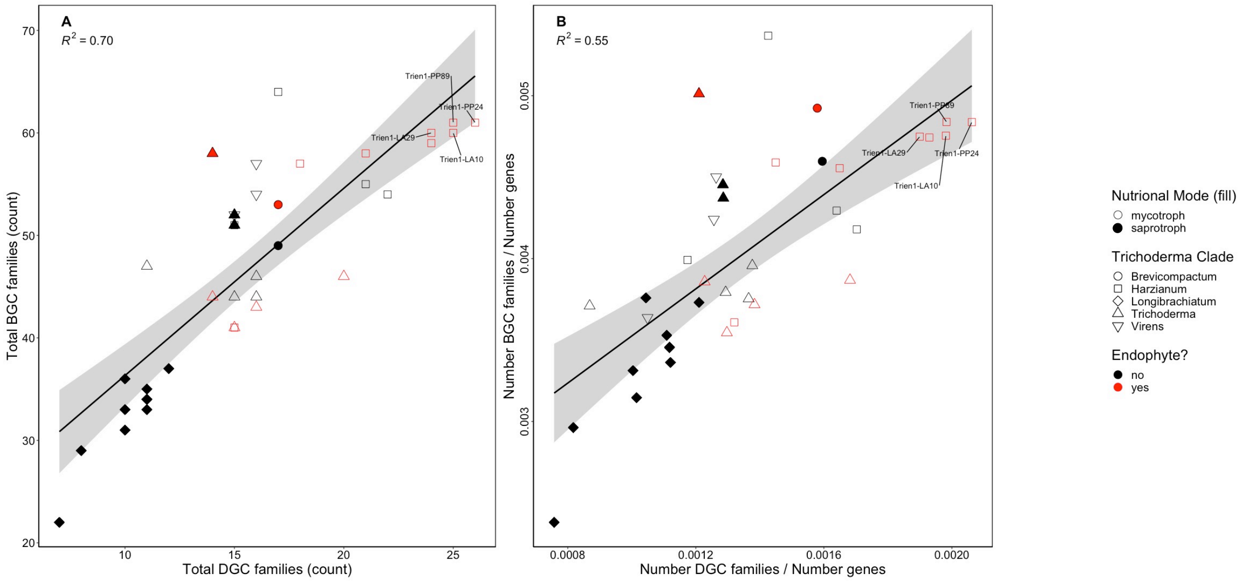


**Fig S3. Genomes with greater numbers of DGCs also contained greater numbers of BGCs.**
(A) The number of DGC families identified in each genome positively correlates with the number of identified biosynthetic gene cluster families (Pearson's, $p < 0.005$). (B) The proportion of DGC families to total number genes per genome positively correlates with the proportion of DGC families to total number genes per genome (Pearson's, $p < 0.005$). Each point represents a *Trichoderma* isolate. Point fill indicates nutritional mode, point shape indicates which clade to which the isolate belongs, and color indicates whether or not the isolate has a possible endophytic lifestyle. Gray shaded areas indicate the 95% confidence interval.

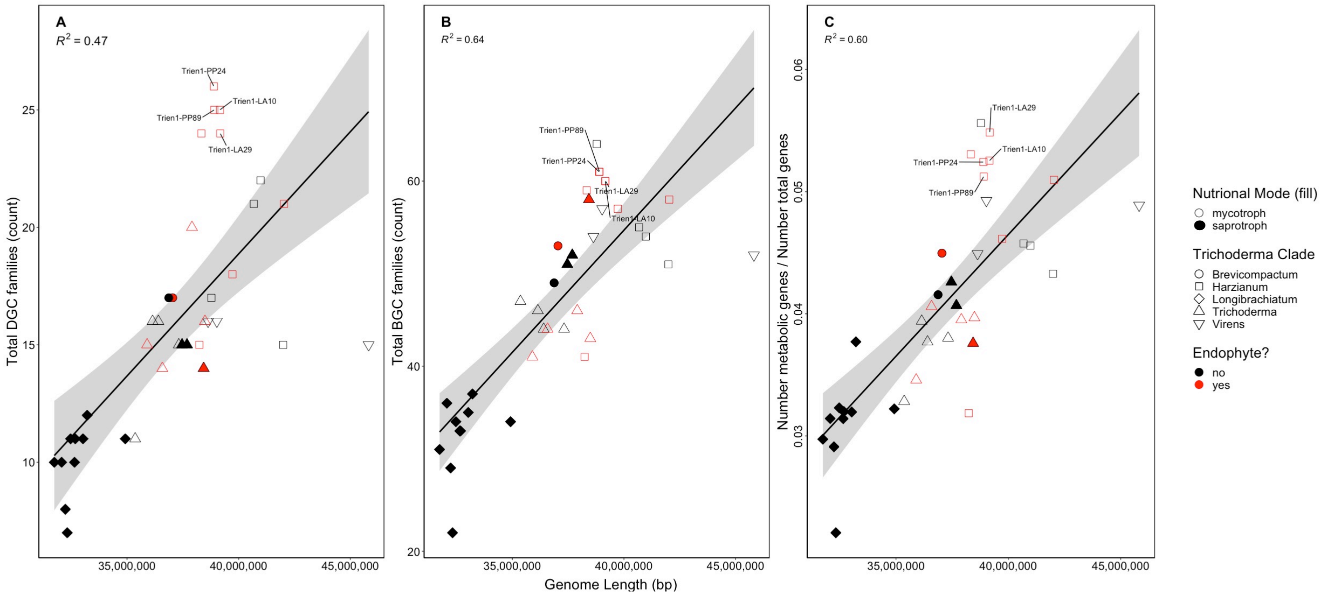


**Fig S4. Larger genomes contain more gene clusters and metabolic genes.**

There is a significant correlation between genome length (bp) of the 38 *Trichoderma* genomes and the diversity of (A) DGC families (Pearson's, $p \leq 0.05$), (B) BGC families (Pearson's, $p \leq 0.05$), and (C) proportion of metabolic genes to total genes (Pearson's, $p \leq 0.05$). Points highlighted in red are isolates identified to have an endophytic lifestyle, points in black did not have an identified endophytic lifestyle, and points in blue were species with a potential endophytic lifestyle. Mycotrophic genomes are represented by empty points, saprotrophic genomes are represented by filled points. Gray shaded areas indicate the 95% confidence interval.

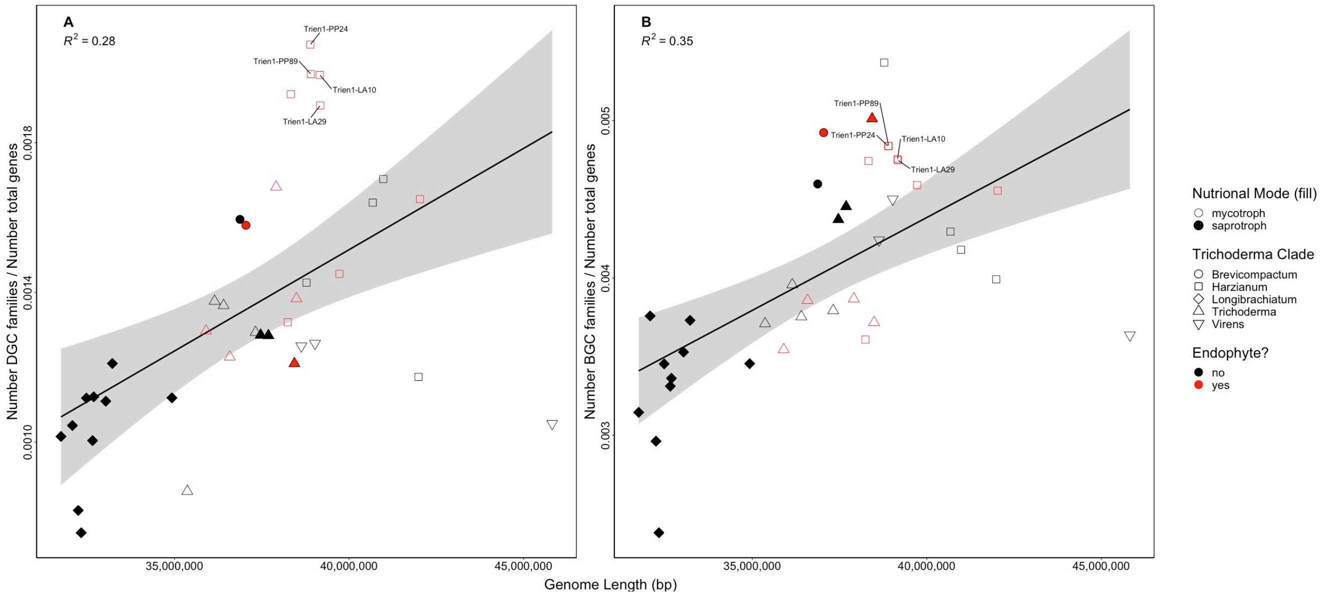


**Fig S5. Larger genomes contain a greater proportion of gene clusters to genes.**
There is a significant correlation between genome length (bp) of the 38 *Trichoderma* genomes and the diversity of (A) proportion of biosynthetic genes to BGC clusters (Pearson's, $p \leq 0.05$) and (B) proportion of DGC families to total genes (Pearson's, $p \leq 0.05$). Points highlighted in red are isolates identified to have an endophytic lifestyle, points in black did not have an identified endophytic lifestyle, and points in blue were species with a potential endophytic lifestyle. Mycotrophic genomes are represented by empty points, saprotrophic genomes are represented by filled points. Gray shaded areas indicate the 95% confidence interval.

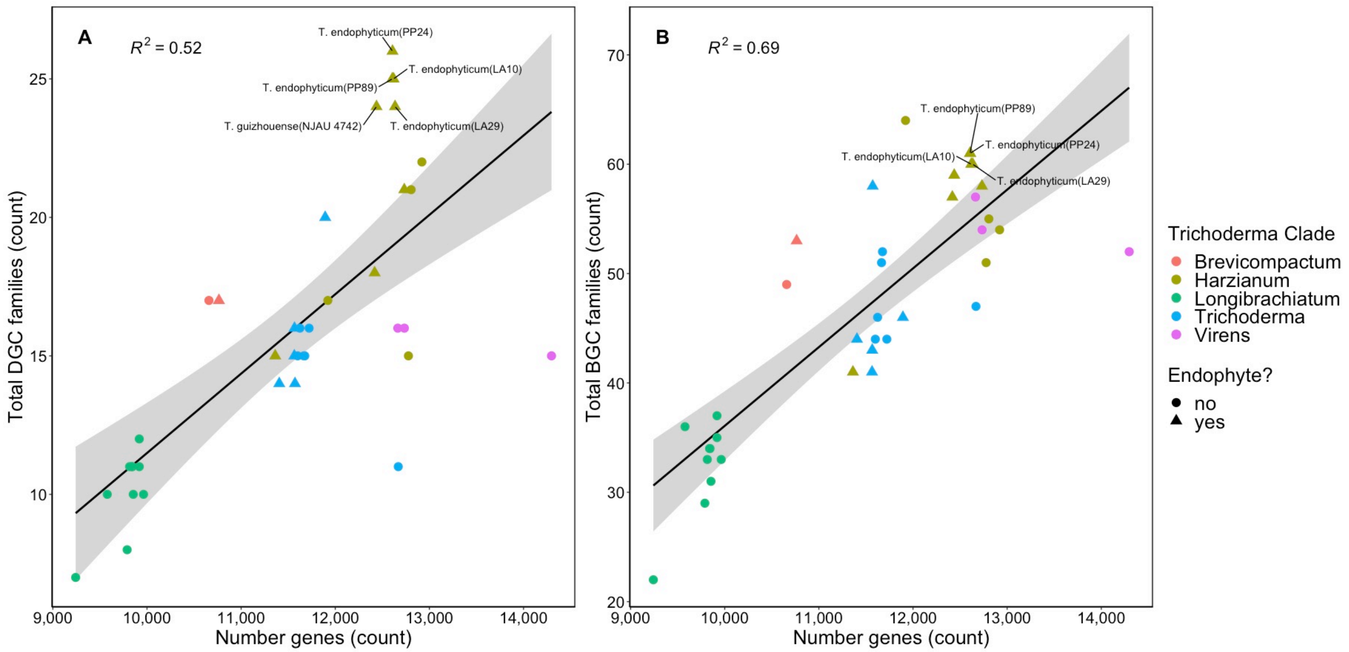


**Fig S6. Genomes with more genes contain greater numbers of metabolic gene clusters.**
The overall number of genes identified in a *Trichoderma* genome is correlated with the number of (A) identified DGCs (Pearson's, $p < 0.005$) and (B) BGCs (Pearson's, $p < 0.005$). Gray shaded areas indicate the 95% confidence interval.

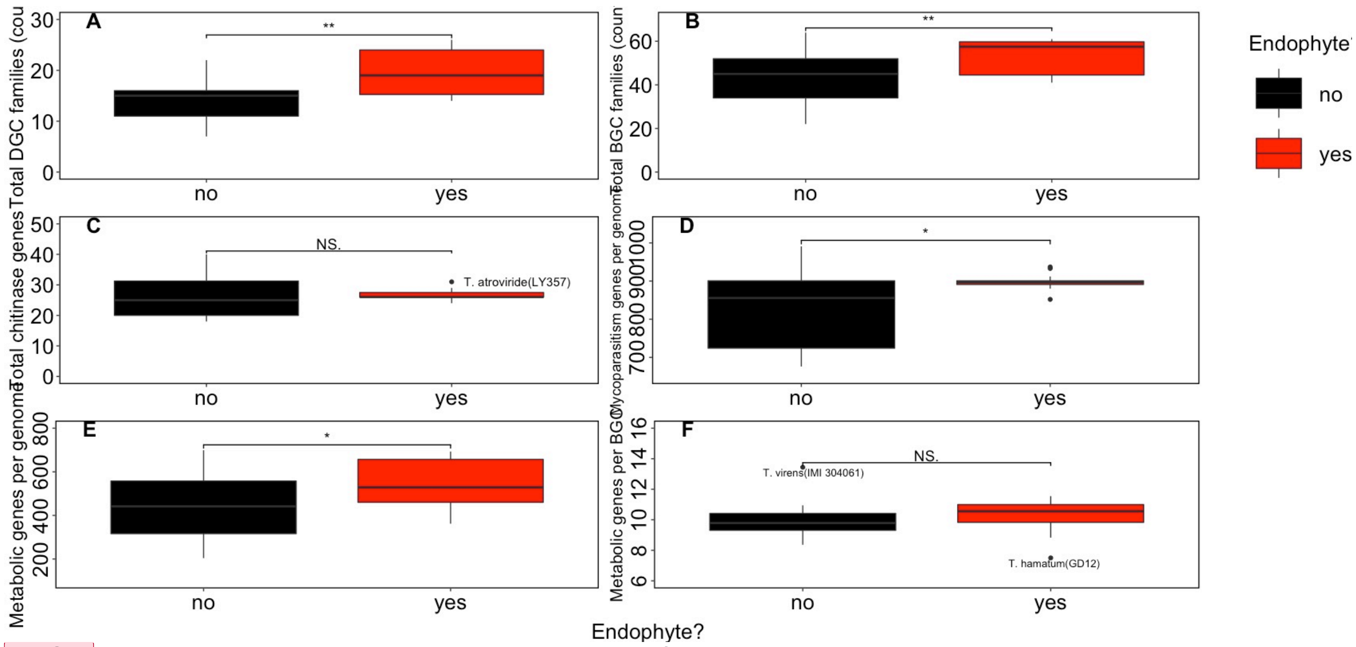


**Fig S7. Endophyte genomes contain greater numbers of metabolic gene clusters and mycoparasitism genes.** The average number metabolic gene clusters is significantly different between endophytic and non-endophytic *Trichoderma* genomes for both (A) DGCs (Wilcoxon, $p \leq 0.05$) and (B) BGCs (two-sample t-test, $p \leq 0.05$). There is no significant differences between the number of (C) chitinase genes for endophytic and non-endophytic *Trichoderma* genomes (Wilcoxon, $p > 0.05$). The number of mycoparasitism genes is significantly higher in (D) endophyte genomes (Wilcoxon, $p \leq 0.05$). (E) The number of metabolic genes per genome were higher in endophytes than in non-endophyte genomes (two-sample t-test, $p \leq 0.05$). The number of metabolic genes per BGC was not significantly different between endophyte and non-endophyte genomes (two-sample t-test, $p > 0.05$).

**Commented [MOU1]:** Double check in-text ref stat tests

**Commented [MOU2R1]:** Fix y-axes

**Commented [MOU3R1]:** Nudge down graph labels

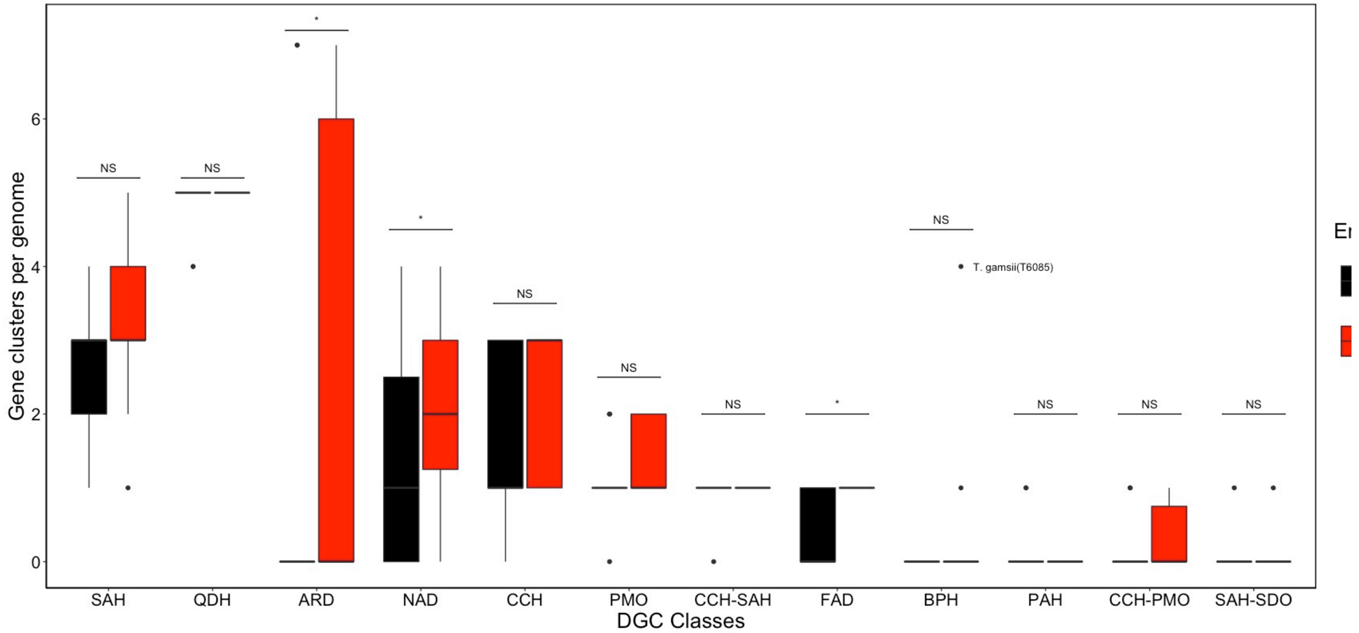


**Fig S8. Endophytic *Trichoderma* genomes contain significantly greater numbers of ARD, NAD, and FAD DGC.** Differences between endophytic and non-endophytic DGC class counts were calculated using Wilcoxon rank sum test corrected with Holm-Bonferroni method. Endophytic genomes have significantly greater ARD, NAD, and FAD (each $p \leq 0.05$), significance differences between groups are indicated with an asterisk. No significant difference between endophyte and non-endophyte DGC count is indicated with “NS” ($p > 0.05$).

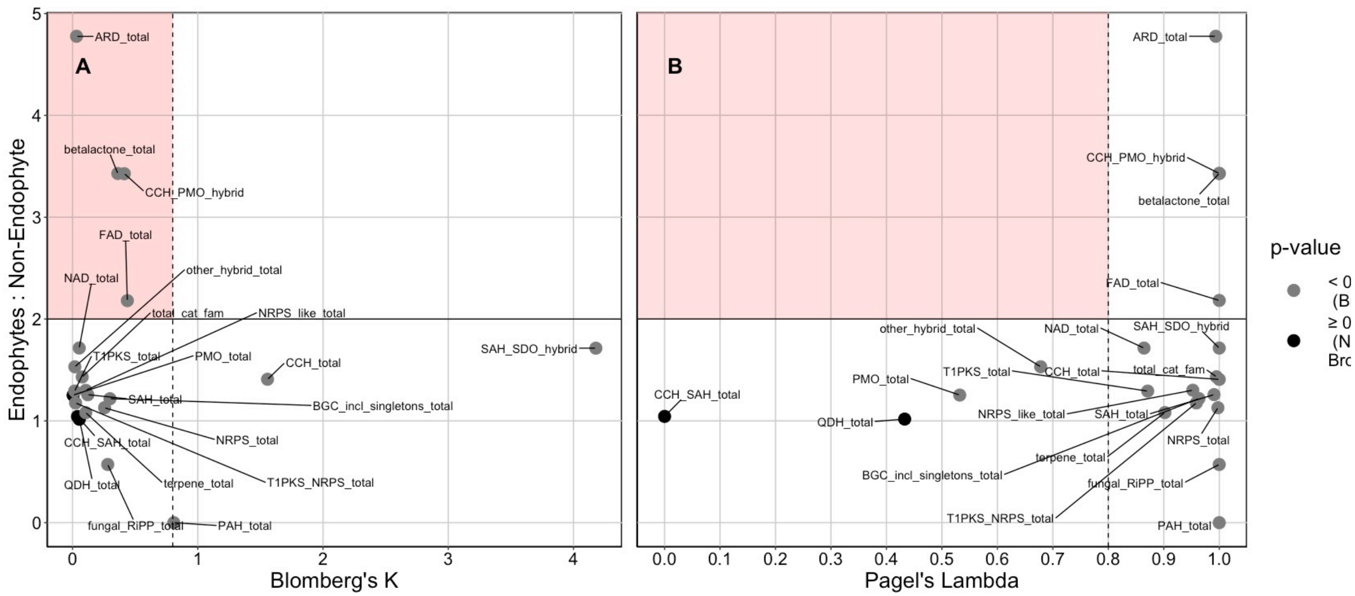


**Fig S9 Total metabolic cluster counts and individual metabolic cluster class distributions are reflected by *Trichoderma* phylogeny.**

Most evaluated DGC and BGC class distributions were determined to be consistent with Brownian motion evolution as determined by their (A) Blomberg's K value and significance (p-value ≤ 0.05) and (B) Pagel's Lambda value and significance (p ≤ 0.05). Black points in the shaded area of the graph would indicate DGC classes which exhibit overdispersal in the *Trichoderma* phylogeny as well as overrepresentation in endophytic genomes.

**Commented [MOU4]:** Edit legend, <= 0.05

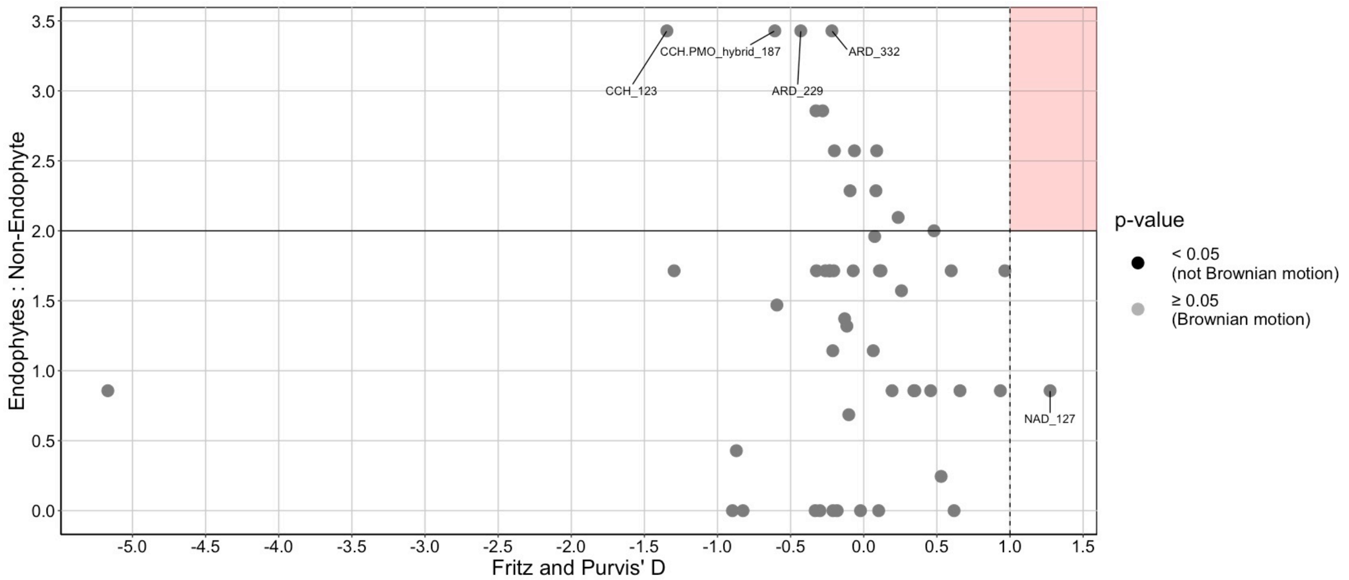


**Fig S10. The distribution of individual DGC families is consistent with *Trichoderma* phylogeny.**
All evaluated DGC families did not have a significant result when analyzing each DGC's Fritz and Purvis' D-statistic ($p > 0.05$), indicating that the distribution of DGC families followed Brownian motion evolution. The DGC families most overrepresented in endophyte genomes (E:NE > 3) are labeled. Black points in the shaded area of the graph would indicate DGC families which exhibit overdispersal in the *Trichoderma* phylogeny as well as overrepresentation in endophytic genomes.

**Commented [MOU5]:** Edit legend, <= 0.05

**Commented [MOU6R5]:** Non-endophyte -> non-endophytes

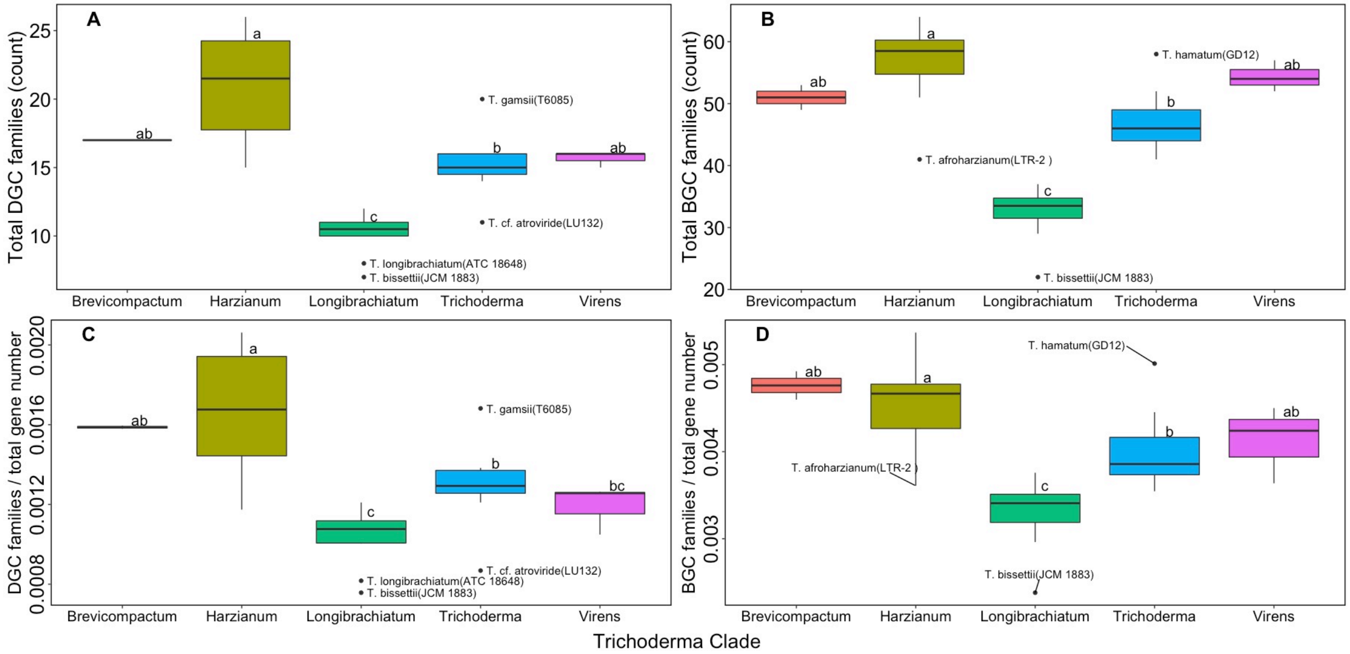


**Fig S11. Metabolic gene cluster content and proportions vary among *Trichoderma* clades.**
There are significant differences between the *Trichoderma* clades for total count of (A) DGCs per genome ($p \leq 0.05$) and (B) BGCs per genome ($p \leq 0.05$). There are also significant differences between clades for the (C) proportion of DGCs to total gene number in each genome ($p \leq 0.05$) and (D) proportion of BGCs to total gene number in each genome( $p \leq 0.05$). Significant differences were determined using Kruskal-Wallis test, post hoc differences were determined with Dunn’s test with Bonferroni correction.

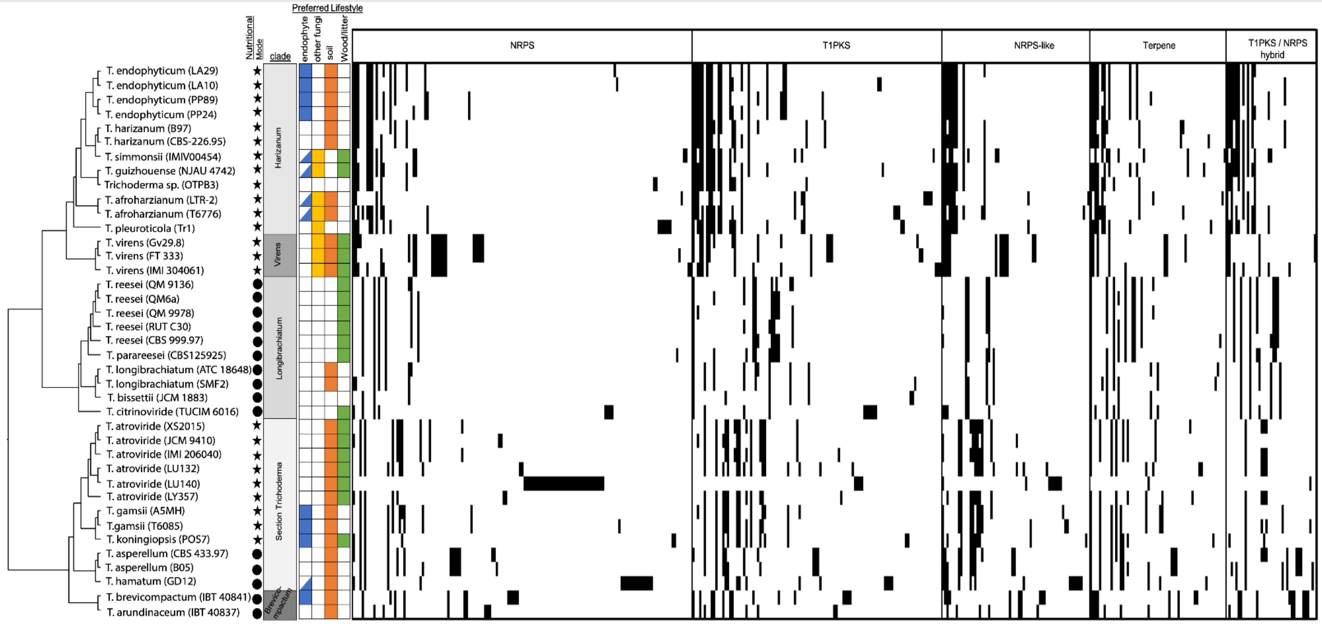


**Fig S12: The number and distribution of different BGC families varies across the Trichoderma phylogeny.** Heat map showing presence/absence of biosynthetic gene cluster families for each *Trichoderma* genome. Singleton BGCs, as well as the highly fragmented *T. cf. atroviride* (LU140) genome, are included in this figure. The total number of BGC per genome, as well as the distribution of fungal-RiPP, other hybrid, and beta-lactone clusters, are unchanged from those in Fig 1 and are thus excluded for visualization purposes. Species lifestyle is indicated by presence/absence of color-coordinated cell, nutritional mode is indicated by either a circle (saprotroph) or a star (mycotroph). Endophyte cells half-filled indicated a potential endophytic lifestyle for that species

.

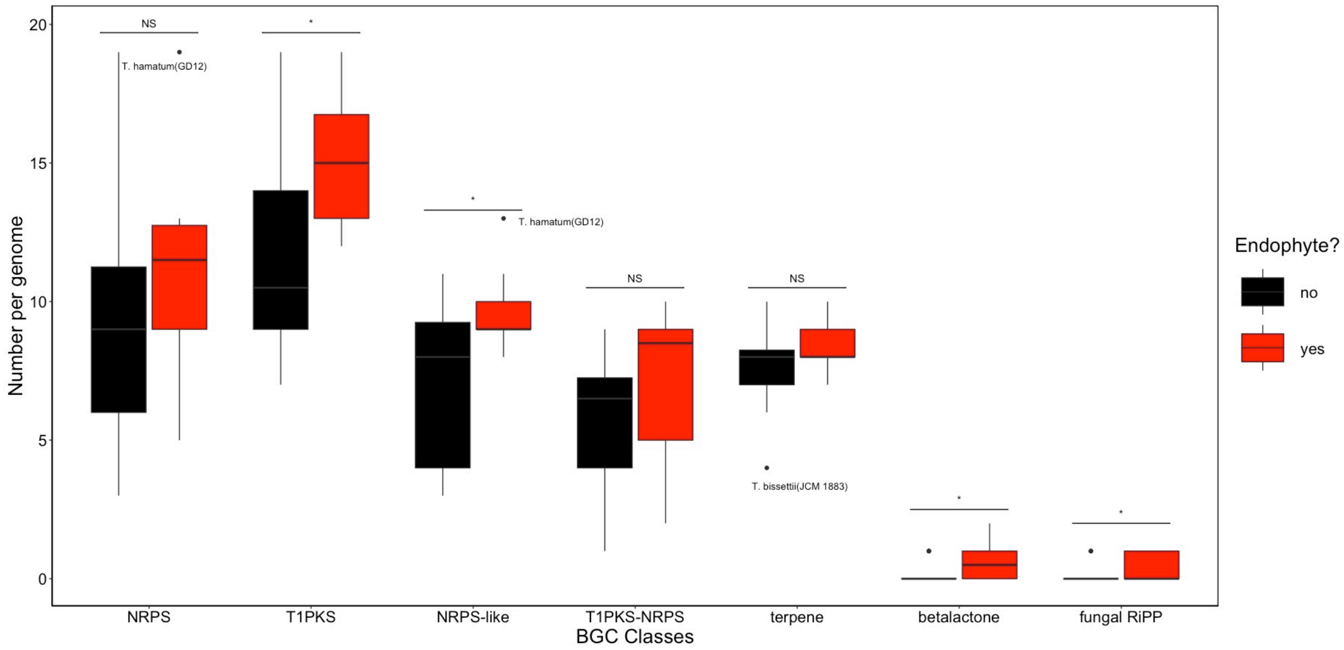


**Fig S13. Endophytic genomes are enriched in T1PKS, NRPS-like, beta-lactone, and fungal-RiPP BGCs.** Significant differences were determined by the Wilcoxon rank sum test corrected with Holm-Bonferroni method. No significant differences between groups are indicated by “NS”. Outliers within groups are labeled.

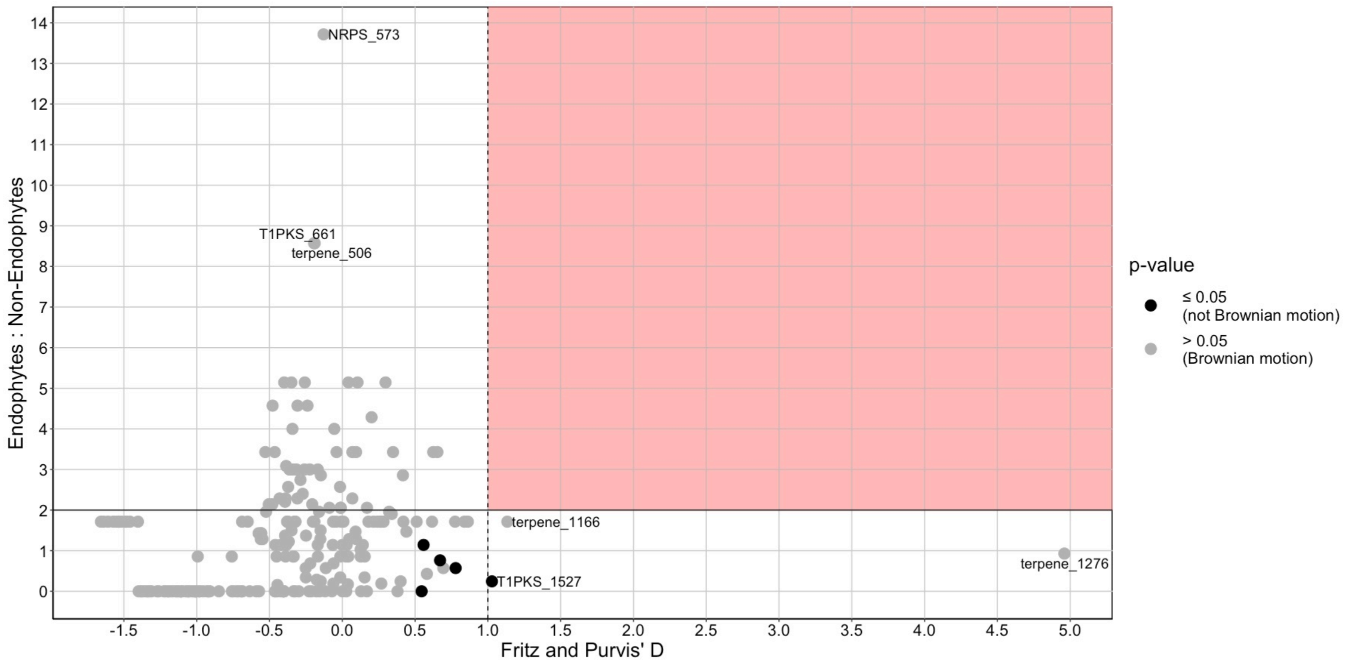


**Fig S14. The distribution of most individual BGC families is consistent with *Trichoderma* phylogeny.**
The majority of evaluated BGC families did not have a significant result when analyzing each's Fritz and Purvis' D-statistic ($p > 0.05$), which indicated Brownian motion evolution. The BGC families that did not exhibit Brownian motion model evolution were not found enriched in endophyte genomes (E:NE < 2). The BGC families most overrepresented in endophyte (E:NE value > 6) and DGCs overdispersed in the *Trichoderma* phylogeny (D-statistic > 1) are labeled. Black points in the shaded area of the graph would indicate BGC families which exhibit overdispersal in the *Trichoderma* phylogeny as well as overrepresentation in endophytic genomes.

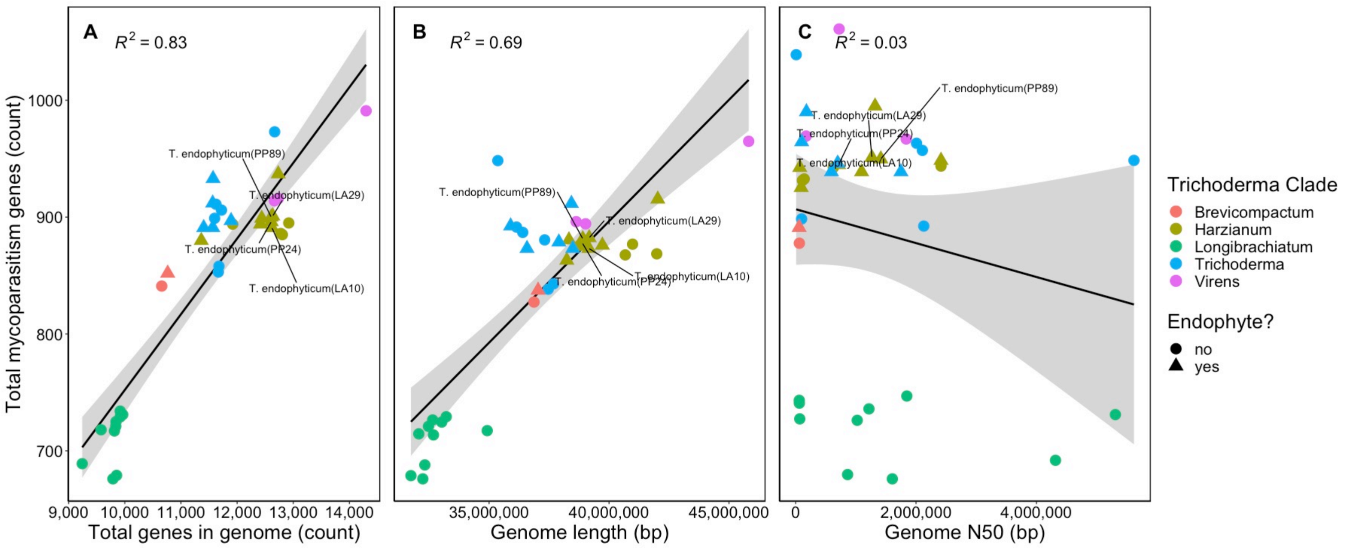


**Fig S15. Genomes with greater numbers of genes and larger genomes contain more mycoparasitism-related genes.**

The total number of mycoparasitism genes per genome is positively correlated with (A) total genes in a Trichoderma genome (Pearson's, $p \leq 0.05$) and (B) genome length (Pearson's, $p \leq 0.05$). (C) There is no relation to genome quality (N50) and number of identified mycoparasitism-related genes per genome (Pearson's, $p > 0.05$). Gray shaded areas indicate the 95% confidence interval.

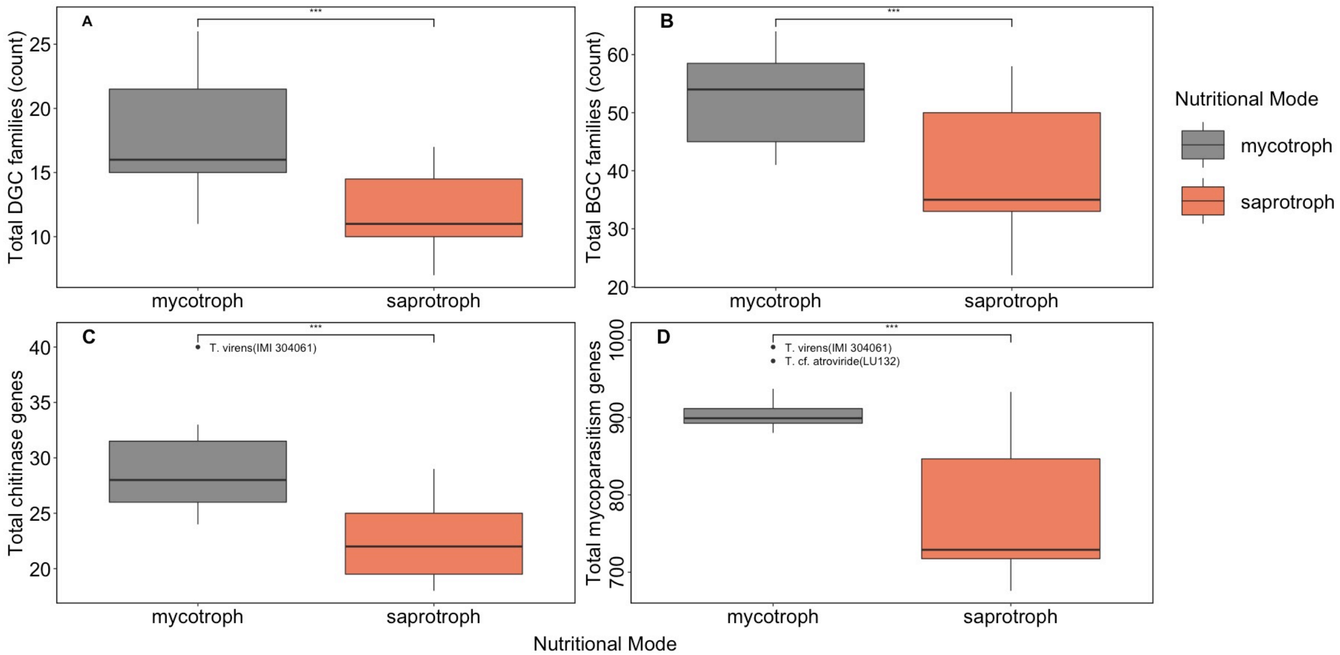


**Fig 16. Metabolic gene clusters and mycoparasitism genes are enriched in mycotrophic genomes.** The average number of metabolic gene clusters is significantly different between mycotrophic and saprotrophic Trichoderma genomes for both A) DGCs (2-sample t-test, $p \leq 0.0005$) and B) BGCs (2-sample t-test, $p \leq 0.0005$). The total number of C) chitinase genes is higher in mycotrophic genomes (2-sample t-test, $p \leq 0.0005$), as are the D) total number of mycoparasitism genes (2-sample t-test, $p \leq 0.0005$). Outliers are labeled and significant differences are indicated with asterisks.

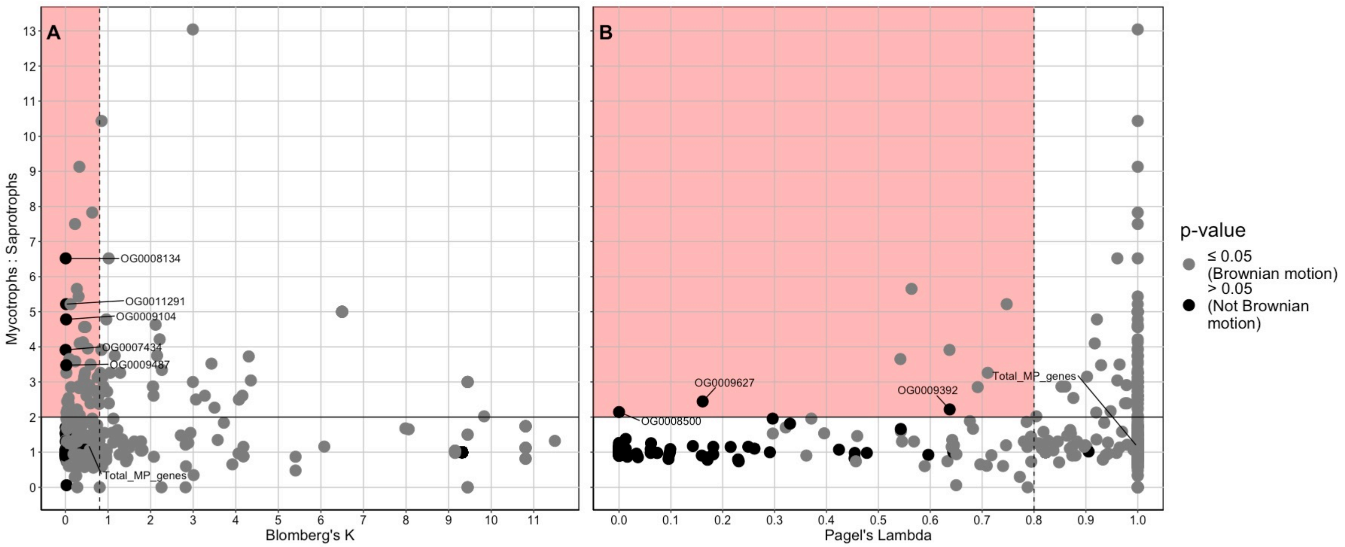


**Fig S17. Some mycoparasitism genes are overrepresented in mycotrophs and overdispersed in *Trichoderma*.**
A) The mycotroph to saprotroph gene ratio (M:S) and Blomberg's K value suggests that five mycoparasitism genes are overdispersed in *Trichoderma* (K < 0.8, p > 0.05) and enriched in genomes with a mycotrophic nutritional mode (M:S > 2). (B) The mycotroph to saprotroph gene ratio (M:S) and Blomberg's K value suggests that three mycoparasitism genes are overdispersed in *Trichoderma* (Lambda < 0.8, p > 0.05) and enriched in genomes with a mycotrophic nutritional mode (M:S > 2). Black points in the shaded area of the graph would indicate mycoparasite genes which exhibit overdispersal in the *Trichoderma* phylogeny as well as overrepresentation in mycotrophic genomes.

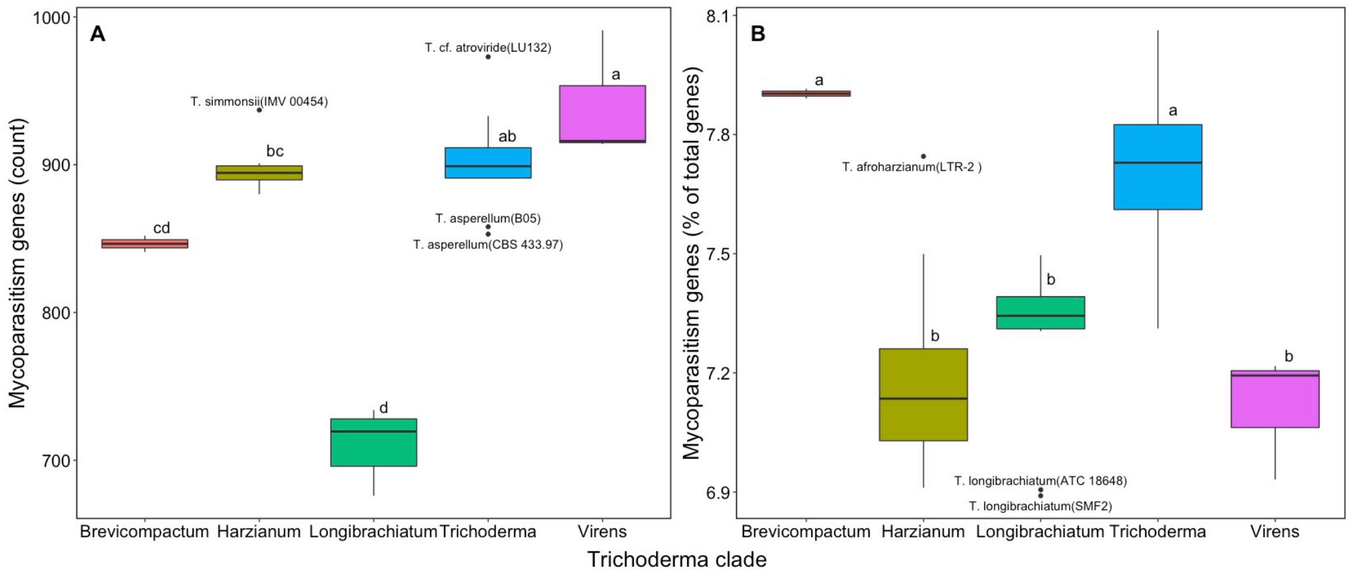


**Fig S18. Amount and percent of mycoparasitism-related genes differ between *Trichoderma* clades.**
(A) Longibrachiatum contains the fewest amount of mycoparasitism-related genes while Virens and section Trichoderma contain the highest amount (Kruskal-Wallis, correction with Bonferroni, $p \leq 0.05$). (B) Harzianum, Longibrachiatum, and Virens contain the lowest percent of mycoparasitism genes to the total number of genes per genome, and Brevicompactum and section Trichoderma contains the highest (ANOVA, correction with Bonferroni, $p \leq 0.05$).

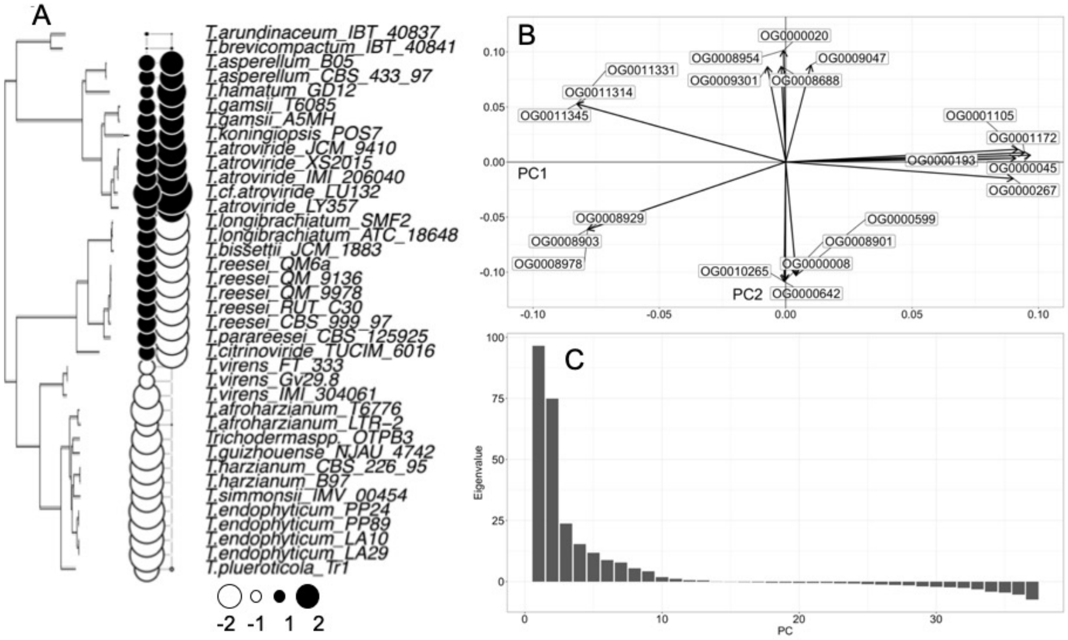


**Fig S19. Phylogenic correction of mycoparasitism data still indicate separation based on *Trichoderma* clade.** (A) Phylogenetic tree and relative signal of phylogenetically-corrected global principal component 1 (PC1) and global principal component 2 (PC2) for each *Trichoderma* genome. (B) Top 10 mycoparasitism genes contributing to PC1 signal and top 11 mycoparasitism genes contributing to PC2 signal. (C) Eigenvalues for the phylogenic correction PCA (pPCA) global principal components (PCs).

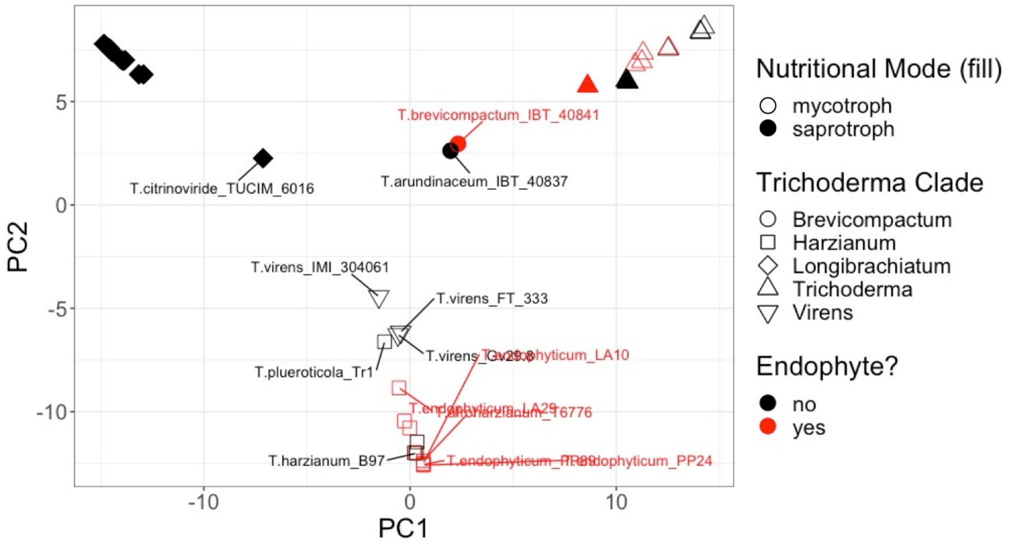


**Fig S20. *Trichoderma* genomes have similar mycoparasitism gene repertoires to other genomes within the same clade.**
Principal components analysis (PCA) of the distribution of the 746 mycoparasitism-related genes; the inherent phylogenetic bias within the *Trichoderma* dataset was corrected with phylogenetic principal components analysis (pPCA). Plotting of the global principal components values for each genome shows that *Trichoderma* genomes are grouping based on *Trichoderma* clade rather than nutritional mode or lifestyle.

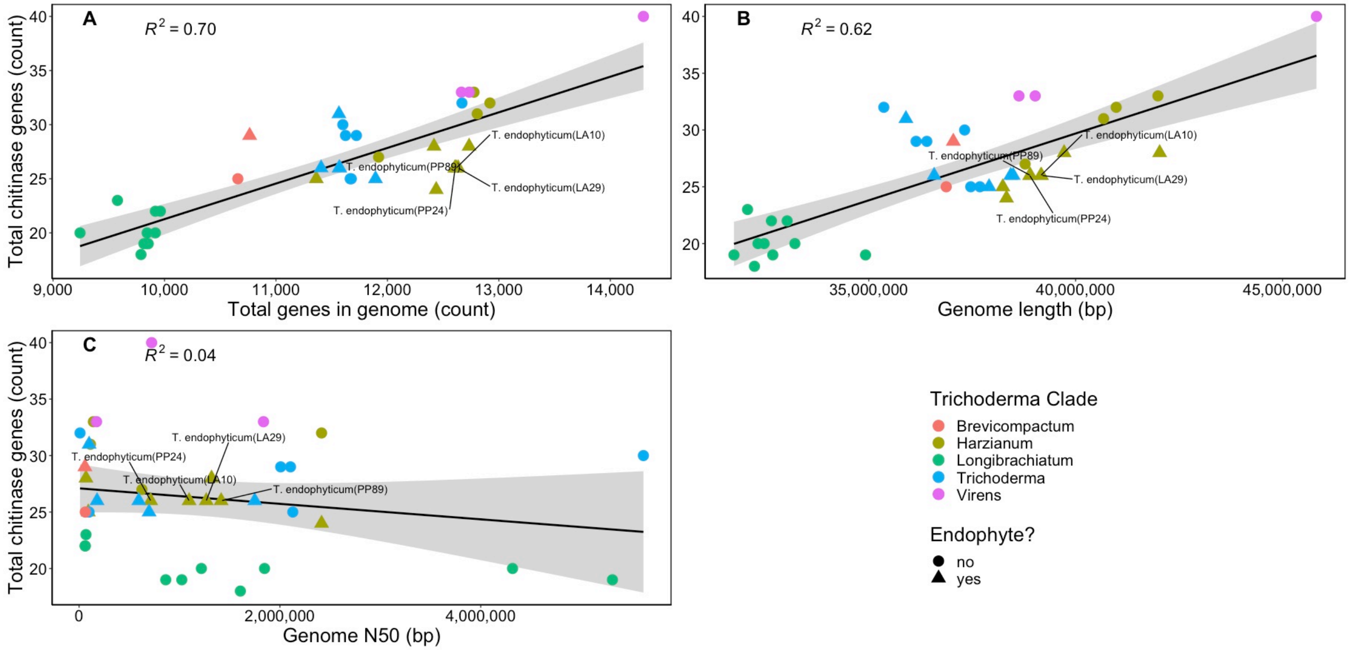


**Fig S21. Larger genomes and genomes that contain more genes contain greater numbers of chitinase genes.** The overall number of genes identified in a *Trichoderma* genome is correlated with the number of A) identified chitinase genes (Pearson's, $p \leq 0.05$) and B) genome length (Pearson's, $p \leq 0.05$), but not with C) genome quality (Pearson's, $p \geq 0.05$). Gray shaded areas indicate the 95% confidence interval.

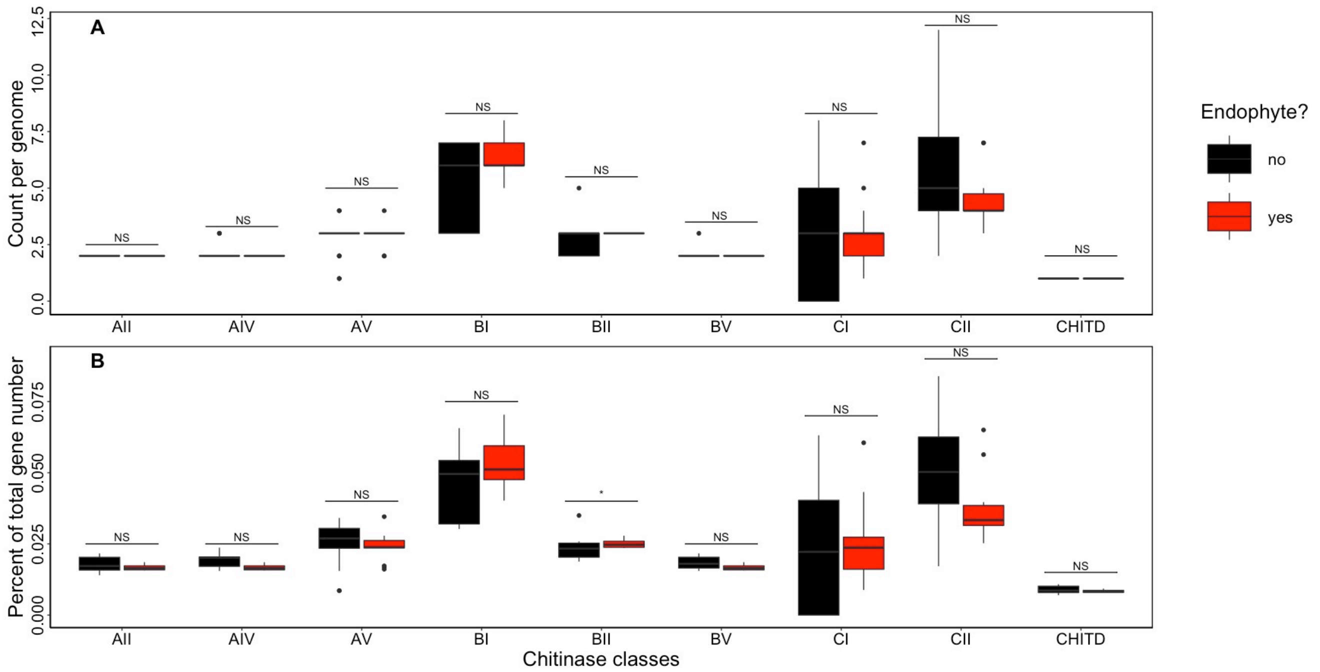


**Fig S22. Endophytes only differ in chitinase BII gene/total gene proportion compared to non-endophytes.**
(A) Endophyte genomes do not significantly differ in any chitinase class overall count when compared with non-endophyte genomes (Wilcoxon, $p > 0.05$) (B) (A) Endophyte genomes only have a significantly higher proportion of chitinase class BII genes to total genes per genome as compared to non-endophyte genomes (Wilcoxon, $p \leq 0.05$).

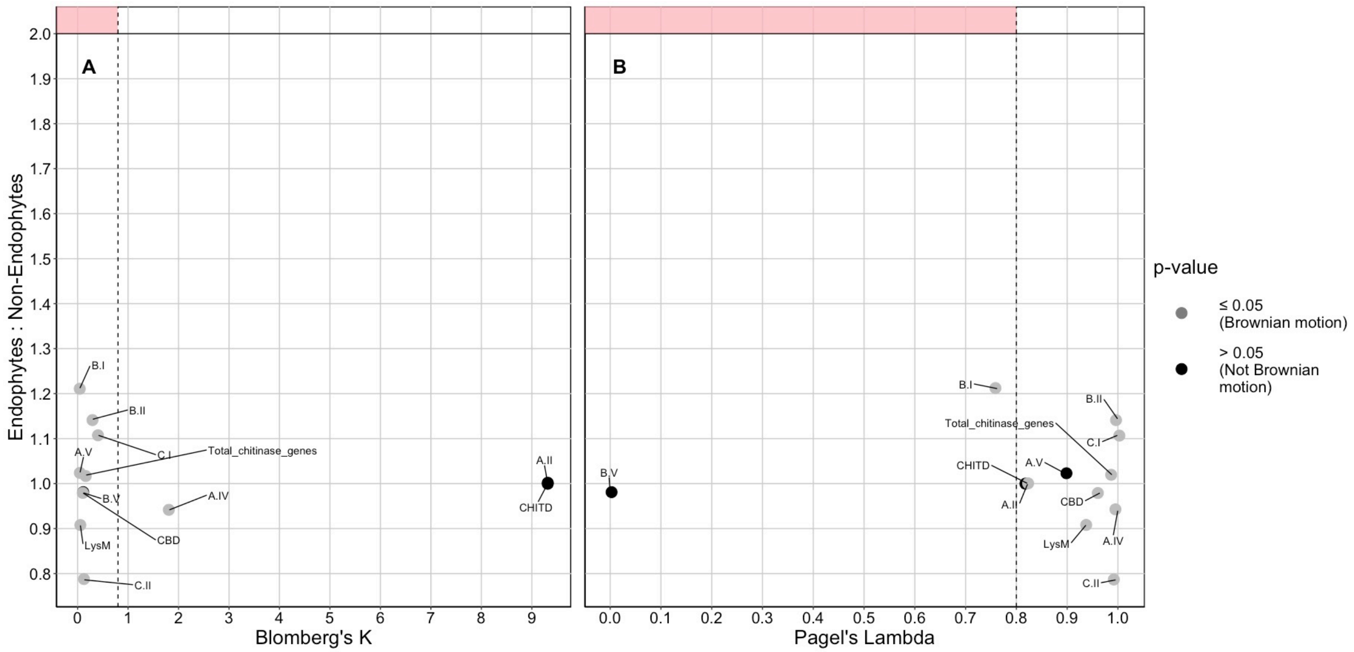


**Fig S23. No chitinase classes are overrepresented in endophytic *Trichoderma* genomes.**
Most evaluated chitinase class distributions were determined to be consistent with Brownian motion evolution as determined by their (A) Blomberg's K value and significance (p-value $\leq$ 0.05) and (B) Pagel's Lambda value and significance (p $\leq$ 0.05). Black points in the shaded area of the graph would indicate chitinase gene classes which exhibit overdispersal in the *Trichoderma* phylogeny as well as overrepresentation in endophytic genomes.

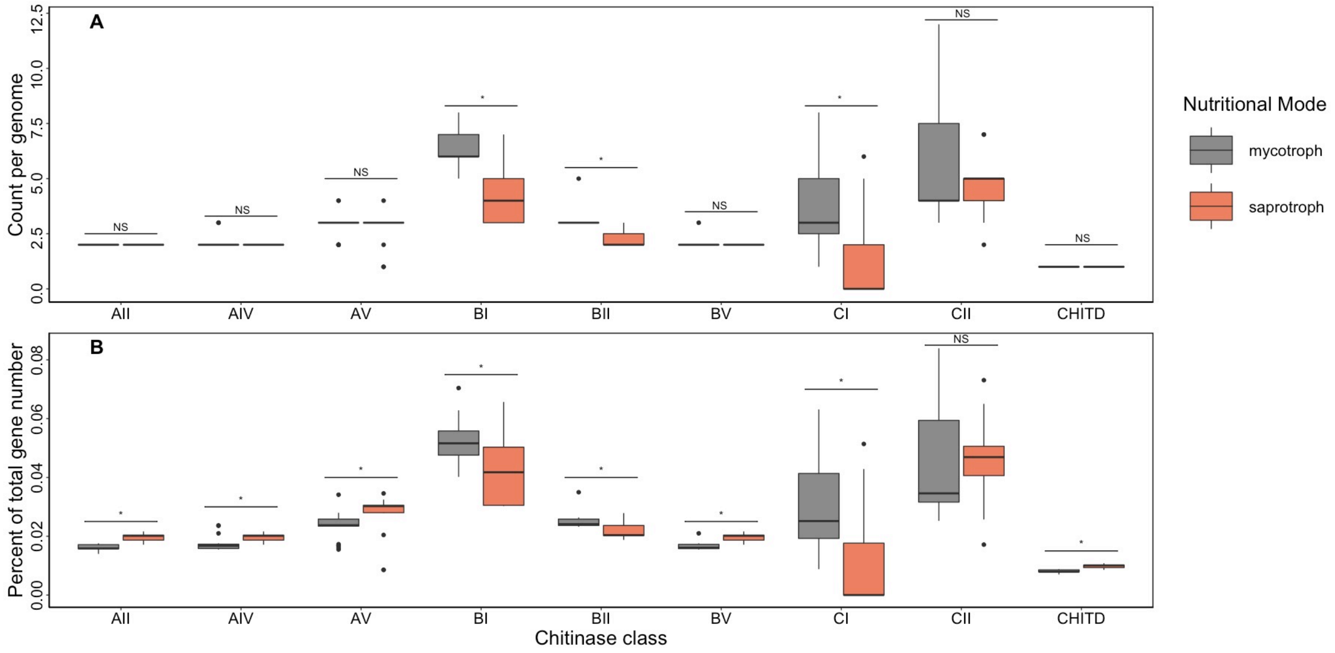


**Fig S24. Mycotrophs differ in both count and proportion of chitinase classes compared to saprotrophs.** (A) Mycotroph genomes have significantly greater numbers of BI, BII, and CI class chitinase genes compared to saprotroph genomes (2-sample t-test, $p \leq 0.05$). (B) Saprotoph genomes have significantly greater percentage of total gene number for AII, AIV, AV, BI, BII, BV, CI, and CHITD class chitinase genes compared to mycotroph genomes (2-sample t-test, $p \leq 0.05$).

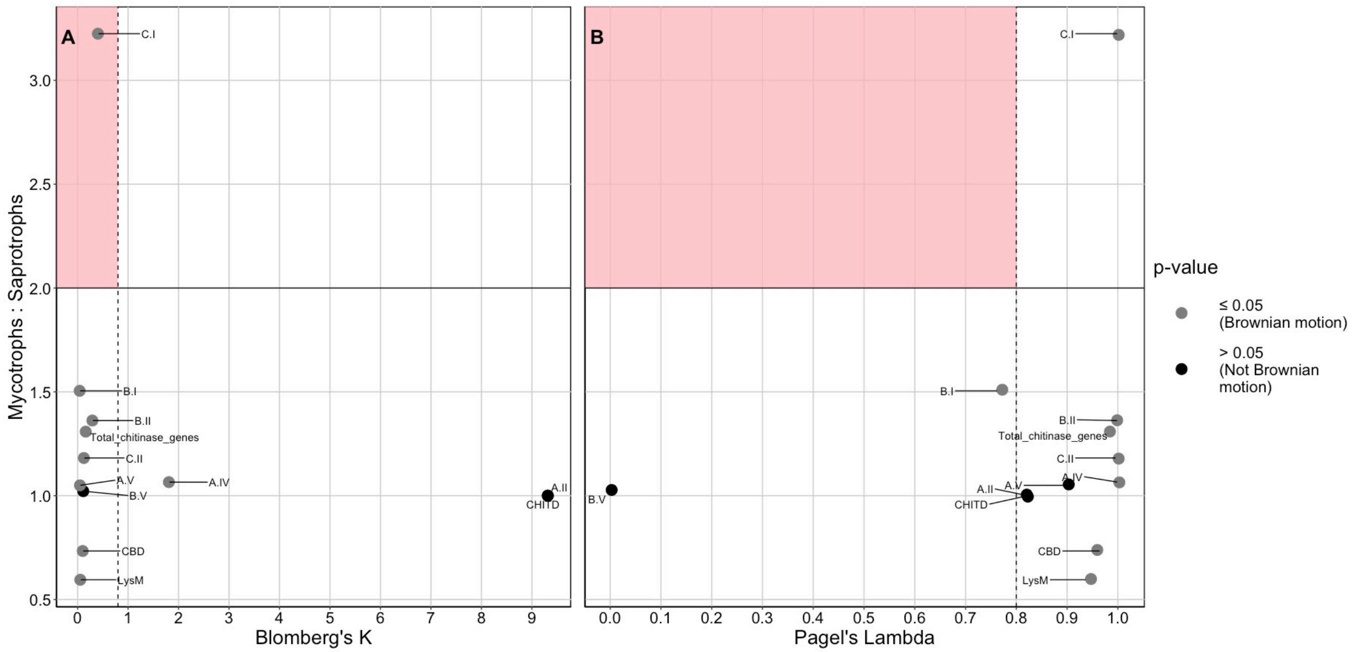


**Fig S25. Most chitinase class distributions were consistent with *Trichoderma* phylogeny.**
Most evaluated chitinase class distributions were determined to be consistent with Brownian motion evolution as determined by their (A) Blomberg's K value and significance (p-value $\leq$ 0.05) and (B) Pagel's Lambda value and significance (p $\leq$ 0.05). Black points in the shaded area of the graph would indicate chitinase gene classes which exhibit overdispersal in the *Trichoderma* phylogeny as well as overrepresentation in mycotroph genomes.

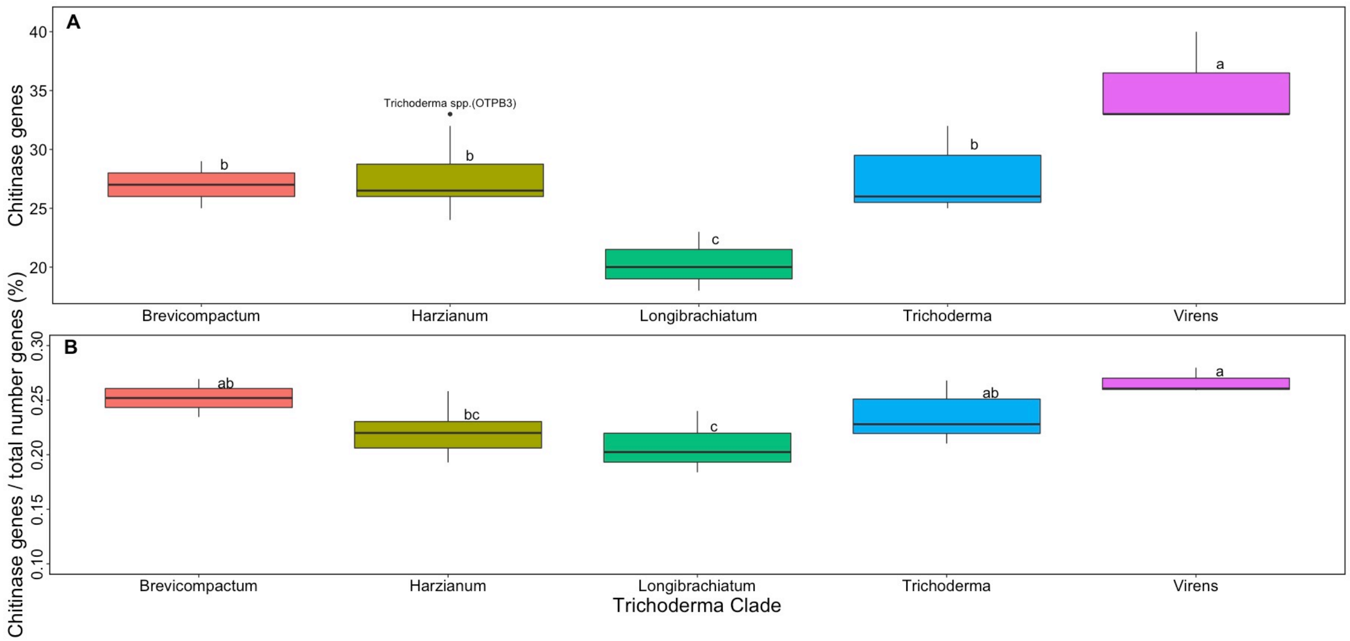


**Fig S26. Genomes in different *Trichoderma* clades differ in their chitinase gene content and proportion.** There are significant differences between *Trichoderma* clades in their (A) number of chitinase genes ($p \leq 0.05$) and (B) percent of chitinase genes ($p \leq 0.05$). Outlier genomes are labeled. Significant differences are indicated by different letters. Differences were calculated with ANOVA, post hoc tested with Tukey's HSD.

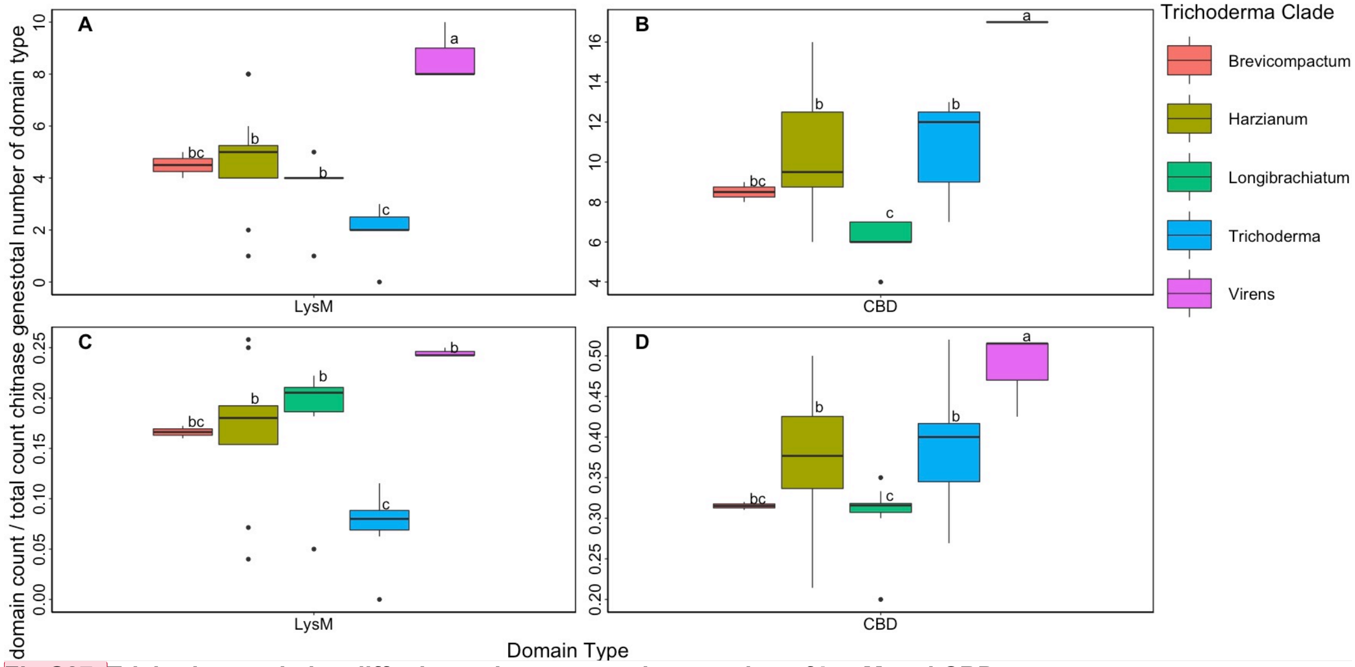


**Fig S27. Trichoderma clades differ in total amount and proportion of LysM and CBD content.** (A) Virens has highest number of LysM and section Trichoderma has the fewest (ANOVA, corrected with Holm-Bonferroni, $p \leq 0.05$). (B) Virens has highest number of CBD; Brevicompactum and section Trichoderma have the fewest (ANOVA, corrected with Holm-Bonferroni, $p \leq 0.05$). (C) Section Trichoderma has the lowest percentage of LysM to total gene count (ANOVA, corrected with Holm-Bonferroni, $p \leq 0.05$). (D) Section Trichoderma has the lowest percentage of CBD to total gene count (ANOVA, corrected with Holm-Bonferroni, $p \leq 0.05$). Significant differences are indicated by different letters, as determined by Tukey's HSD.

**Commented [MOU7]:** fix y axis (too close)

**Commented [MOU8R7]:** specify that C and D are percentages

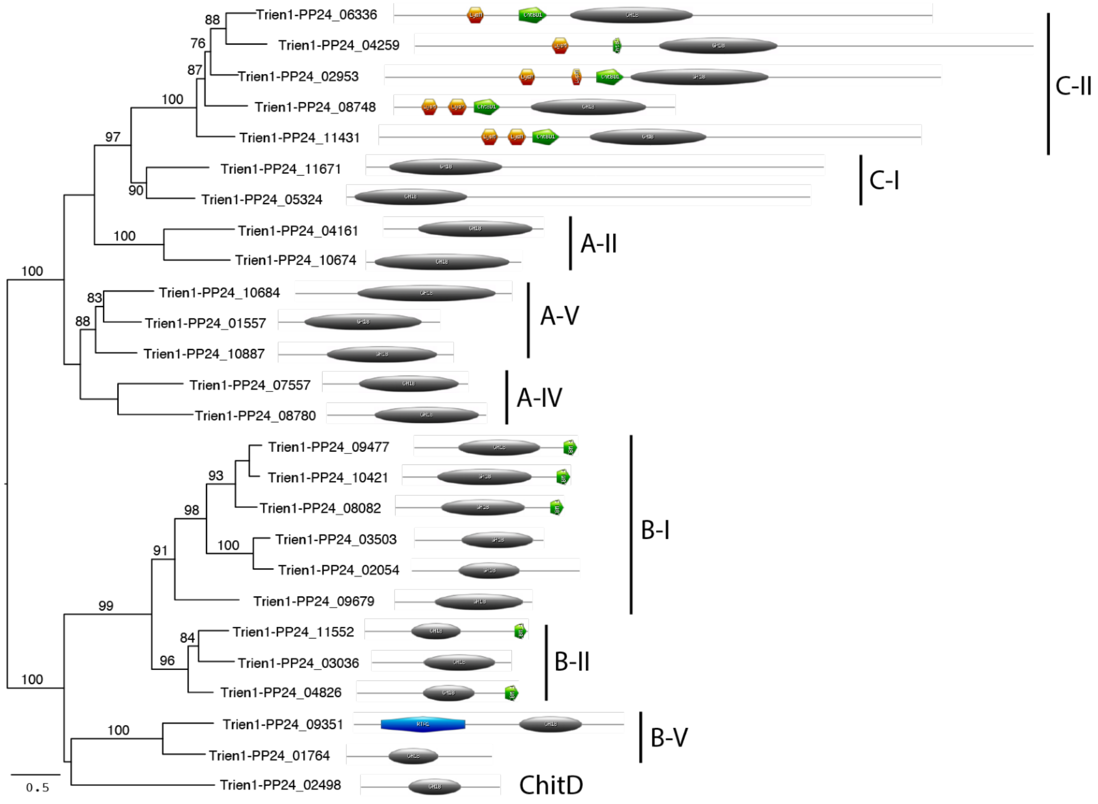


**Fig S28. Domain architecture of the 26 chitinases in *T. endophyticum* strain PP24.**
Chitinase classes are indicated according to (Goughenour 2020). GH-18 domains are represented by black ovals, chitin-binding domains (CBD) are represented as green pentagons, Lysin domains (LysM) are represented as orange hexagons, and an RTA1 Superfamily domain is represented by a blue cylinder.

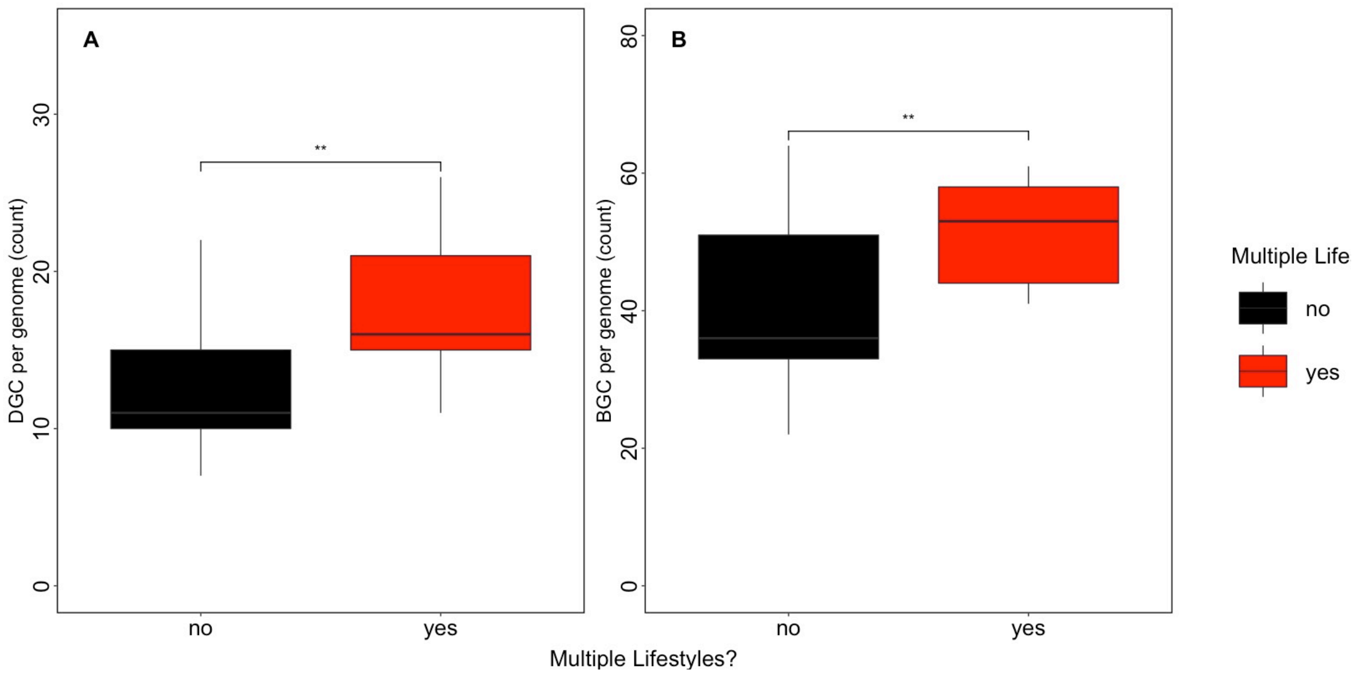


**Fig S29. Trichoderma genomes with multiple potential lifestyles have more metabolic gene clusters.**
Genomes of *Trichoderma* species recorded as having multiple lifestyles have (A) a significantly greater diversity of DGC (two-sample t-test, $p \leq 0.05$) and (B) a significantly greater number of BGC (two-sample t-test, $p \leq 0.05$) as compared to genomes of *Trichoderma* species recorded as only having a single potential lifestyle.

**Table S1. Species name, strain identifier, source of genomic reads, lifestyle information, nutritional mode, and geographic origin of each *Trichoderma* isolate used in this study.**

| Verified species name | Given species name | Assembly source (SRA accession) | Genome assembly accession | Strain | Nutritional mode | Lifestyle | Country of Origin | Lifestyle (Endophyte or non-endophyte) |
|---|---|---|---|---|---|---|---|---|
| *T. afroharzianum** | *T. harzianum* | NCBI | GCA_000988865.1 | T6776 | mycotroph | fungi, soil | Italy | Endophyte |
| *T. afroharzianum** | *T. viride* | NCBI (SRX751293) | This study | LTR-2 | mycotroph | fungi, soil | China | Endophyte |
| *T. arundinaceum* | *T. arundinaceum* | NCBI | GCA_003012105.1 | IBT 40837 | saprotroph | soil | Iran | Non-endophyte |
| *T. asperellum* | *T. asperellum* | NCBI | GCA_000733085.2 | B05 | saprotroph | soil | France | Non-endophyte |
| *T. asperellum* | *T. asperellum* | NCBI | GCA_003025105.1 | CBS 433.97 | saprotroph | soil | United States | Non-endophyte |
| *T. atroviride* | *T. atroviride* | NCBI | GCA_001599035.1 | JCM 9410 | mycotroph | soil, wood/litter | Japan | Non-endophyte |
| *T. atroviride* | *T. atroviride* | NCBI | GCA_002916895.1 | LY357 | mycotroph | soil, wood/litter | China | Non-endophyte |
| *T. atroviride* | *T. atroviride* | NCBI | GCA_000963795.1 | XS2015 | mycotroph | soil, wood/litter | The Netherlands | Non-endophyte |
| *T. atroviride* | *T. atroviride* | NCBI | GCA_000171015.2 | IMI 206040 | mycotroph | soil, wood/litter | Sweden | Non-endophyte |

| Verified species name | Given species name | Assembly source (SRA accession) | Genome assembly accession | Strain | Nutritional mode | Lifestyle | Country of Origin | Lifestyle (Endophyte or non-endophyte) |
|---|---|---|---|---|---|---|---|---|
| *T. bissettii** | *T. koningii* | NCBI | GCA_001950475.1 | JCM 1883 | saprotroph | only known from a human sinus cavity | Wales | Non-endophyte |
| *T. brevicompactum* | *T. brevicompactum* | NCBI | GCA_003012085.1 | IBT 40841 | saprotroph | endophyte, soil | Iran | Endophyte |
| *T. cf. atroviride* | *T. cf. atroviride* | NCBI (SRX1601956) | This study | LU132 | mycotroph | soil, wood/litter | New Zealand | Non-endophyte |
| *T. cf. atroviride* | *T. cf. atroviride* | NCBI (SRX1605616) | This study | LU140 | mycotroph | soil, wood/litter | New Zealand | Non-endophyte |
| *T. citrinoviride* | *T. citrinoviride* | NCBI | GCA_003025115.1 | TUCIM 6016 | saprotroph | wood/litter | Unknown | Non-endophyte |
| *T. endophyticum* | *T. endophyticum* | **This study** | This study | LA10 | mycotroph | endophyte, soil | Peru | Endophyte |
| *T. endophyticum* | *T. endophyticum* | **This study** | This study | LA29 | mycotroph | endophyte, soil | Peru | Endophyte |
| *T. endophyticum* | *T. endophyticum* | **This study** | This study | PP24 | mycotroph | endophyte, soil | Peru | Endophyte |

| Verified species name | Given species name | Assembly source (SRA accession) | Genome assembly accession | Strain | Nutritional mode | Lifestyle | Country of Origin | Lifestyle (Endophyte or non-endophyte) |
|---|---|---|---|---|---|---|---|---|
| *T. endophyticum* | *T. endophyticum* | **This study** | This study | PP89 | mycotroph | endophyte, soil | Peru | Endophyte |
| *T. gamsii* | *T. gamsii* | NCBI | GCA_002894205.1 | A5MH | mycotroph | endophyte, soil | Australia | Endophyte |
| *T. gamsii* | *T. gamsii* | NCBI | GCA_001481775.2 | T6085 | mycotroph | endophyte, soil | Ukraine | Endophyte |
| *T. guizhouense* | *T. guizhouense* | NCBI | GCA_002022785.1 | NJAU 4742 | mycotroph | fungi, wood/litter | China | Endophyte |
| *T. hamatum* | *T. hamatum* | NCBI | GCA_000331835.2 | GD12 | saprotroph | soil | England | Endophyte |
| *T. harzianum* | *T. harzianum* | NCBI | GCA_001990665.1 | B97 | mycotroph | soil | France | Non-endophyte |
| *T. harzianum* | *T. harzianum* | NCBI | GCA_003025095.1 | CBS 226.95 | mycotroph | soil | England | Non-endophyte |
| *T. koningiopsis* | *T. koningiopsis* | NCBI | GCA_002246955.1 | POS7 | mycotroph | endophyte, soil | Argentina | Endophyte |
| *T. longibrachiatum* | *T. longibrachiatum* | NCBI | GCA_003025155.1 | ATCC 18648 | saprotroph | soil | United States | Non-endophyte |
| *T. longibrachiatum* | *T. longibrachiatum* | NCBI | GCA_000332775.1 | SMF2 | saprotroph | soil | China | Non-endophyte |

| Verified species name | Given species name | Assembly source (SRA accession) | Genome assembly accession | Strain | Nutritional mode | Lifestyle | Country of Origin | Lifestyle (Endophyte or non-endophyte) |
|---|---|---|---|---|---|---|---|---|
| *T. parareesei* | *T. parareesei* | NCBI | GCA_001050175.1 | CBS 125925 | saprotroph | wood/litter | Argentina | Non-endophyte |
| *T. pleuroticola** | *T. harzianum* | NCBI | GCA_002894145.1 | Tr1 | mycotroph | fungi | China | Non-endophyte |
| *T. reesei* | *T. reesei* | NCBI | GCA_001999515.1 | CBS 999.97 | saprotroph | wood/litter | French Guiana | Non-endophyte |
| *T. reesei* | *T. reesei* | NCBI | GCA_002006585.1 | QM6a | saprotroph | wood/litter | Solomon Islands | Non-endophyte |
| *T. reesei* | *T. reesei* | NCBI (SRX059777) | This study | QM 9136 | saprotroph | wood/litter | Mutant derived from QM6a | Non-endophyte |
| *T. reesei* | *T. reesei* | NCBI (SRX060131) | This study | QM 9978 | saprotroph | wood/litter | Mutant derived from QM6a | Non-endophyte |
| *T. reesei* | *T. reesei* | NCBI | GCA_000513815.1 | RUT C30 | saprotroph | wood/litter | United States | Non-endophyte |
| *T. simmonsii** | *T. virens* | NCBI | GCA_001931985.1 | IMV 00454 | mycotroph | fungi, wood/litter | Ukraine | Endophyte |
| *T. virens* | *T. virens* | NCBI | GCA_000800515.1 | FT 333 | mycotroph | fungi, soil, wood/litter | Taiwan | Non-endophyte |
| *T. virens* | *T. virens* | NCBI | GCA_001835465.1 | IMI 304061 | mycotroph | fungi, soil, wood/litter | India | Non-endophyte |

| Verified species name | Given species name | Assembly source (SRA accession) | Genome assembly accession | Strain | Nutritional mode | Lifestyle | Country of Origin | Lifestyle (Endophyte or non-endophyte) |
|---|---|---|---|---|---|---|---|---|
| *T. virens* | *T. virens* | NCBI | GCA_000170995.2 | Gv29.8 | mycotroph | fungi, soil, wood/litter | Unknown | Non-endophyte |
| *Trichoderma sp.** | *T. harzianum* | NCBI (SRX1433332 ) | This study | OTPB3 | mycotroph | unknown | India | Non-endophyte |

*Some isolates were previously identified as different species but were re-identified for this study.

**Table S2. Genome statistics of the 39 Trichoderma isolates.**

| Verified species name | Strain | Complete and single-copy BUSCOs (%) | Number genes (post-Orthofiller) | N50 | Genome length (Mb) |
|---|---|---|---|---|---|
| *T. arundinaceum* | IBT 40837 | 96.7 | 10,658 | 61,935 | 36.9 |
| *T. asperellum* | B05 | 96.4 | 11,676 | 99,299 | 37.7 |
| *T. asperellum* | CBS 433.97 | 98.8 | 11,666 | 2,128,412 | 37.5 |
| *T. atroviride* | JCM 9410 | 98.9 | 11,600 | 5,619,901 | 37.3 |
| *T. cf. atroviride* | LU132 | 72.7 | 12,669 | 9,580 | 35.4 |
| *T. cf. atroviride* | LU140 | 61.2 | 8,243 | 5,723 | 33.7 |

| Verified species name | Strain | Complete and single-copy BUSCOs (%) | Number genes (post-Orthofiller) | N50 | Genome length (Mb) |
|---|---|---|---|---|---|
| *T. atroviride* | LY357 | 98.5 | 11,565 | 100,133 | 35.9 |
| *T. atroviride* | XS2015 | 98.9 | 11,722 | 2,105,403 | 36.4 |
| *T. atroviride* | IMI 206040 | 98.7 | 11,624 | 2,007,903 | 36.1 |
| *T. brevicompactum* | IBT 40841 | 97.4 | 10,764 | 58,500 | 37.0 |
| *T. citrinoviride* | TUCIM 6016 | 95.5 | 9,918 | 1,846,965 | 33.2 |
| *T. endophyticum* | LA10 | 99.0 | 12,619 | 1,095,838 | 39.2 |
| *T. endophyticum* | LA29 | 99.0 | 12,635 | 1,266,890 | 39.2 |
| *T. endophyticum* | PP24 | 99.0 | 12,608 | 713,317 | 38.9 |
| *T. endophyticum* | PP89 | 99.1 | 12,606 | 1,414,947 | 38.9 |
| *T. gamsii* | A5MH | 98.7 | 11,567 | 594,470 | 38.5 |
| *T. gamsii* | T6085 | 98.8 | 11,893 | 697,391 | 37.9 |

| Verified species name | Strain | Complete and single-copy BUSCOs (%) | Number genes (post-Orthofiller) | N50 | Genome length (Mb) |
|---|---|---|---|---|---|
| *T. guizhouense* | NJAU 4742 | 98.1 | 12,438 | 2,414,983 | 38.3 |
| *T. harzianum* | B97 | 98.3 | 12,806 | 113,316 | 40.7 |
| *T. harzianum* | CBS 226.95 | 98.8 | 12,920 | 2,414,909 | 41.0 |
| *Trichoderma sp.* | OTPB3 | 98.1 | 12,777 | 142,666 | 42.0 |
| *T. afroharzianum* | T6776 | 98.0 | 12,418 | 68,846 | 39.7 |
| *T. pleuroticola* | Tr1 | 98.1 | 11,921 | 626,242 | 38.8 |
| *T. hamatum* | GD12 | 97.9 | 11,572 | 179,274 | 38.4 |
| *T. bissettii* | JCM 1883 | 98.5 | 9,242 | 4,316,785 | 32.3 |
| *T. koningiopsis* | POS7 | 97.4 | 11,405 | 1,749,579 | 36.6 |
| *T. longibrachiatum* | ATCC 18648 | 87.5 | 9,789 | 1,606,350 | 32.2 |
| *T. longibrachiatum* | SMF2 | 98.4 | 9,854 | 863,071 | 31.7 |

| Verified species name | Strain | Complete and single-copy BUSCOs (%) | Number genes (post-Orthofiller) | N50 | Genome length (Mb) |
|---|---|---|---|---|---|
| *T. parareesei* | CBS 125925 | 97.1 | 9,578 | 68,608 | 32.1 |
| *T. reesei* | CBS 999.97 | 98.6 | 9,844 | 1,217,941 | 32.5 |
| *T. reesei* | QM6a | 99.0 | 9,839 | 5,311,445 | 34.9 |
| *T. reesei* | QM 9136 | 97.8 | 9,963 | 60,397 | 32.6 |
| *T. reesei* | QM 9978 | 97.9 | 9,918 | 61,591 | 33.0 |
| *T. reesei* | RUT C30 | 98.8 | 9,815 | 1,023,047 | 32.7 |
| *T. virens* | FT 333 | 98.1 | 12,734 | 173,918 | 38.6 |
| *T. virens* | IMI 304061 | 98.8 | 14,297 | 722,602 | 45.8 |
| *T. afroharzianum* | LTR-2 | 98.8 | 11,362 | 92,065 | 38.2 |
| *T. simmonsii* | IMV 00454 | 98.5 | 12,733 | 1,319,489 | 42.0 |
| *T. virens* | Gv29.8 | 99.0 | 12,664 | 1,836,662 | 39.0 |

**Table S3. Mycoparasitism gene orthogroups of interest found in the highest endophyte:non-endophyte ratio.**

| Mycoparasitism Gene orthogroup | Pagel's Lambda | Pagel's Lambda p-value | Blomberg's K | Blomberg's K p-value | M:S* | E:NE** | Eggnog Annotation |
|---|---|---|---|---|---|---|---|
| OG0010098 | 0.999 | 8.71E-12 | 0.325 | 0.001 | 9.130 | 4.714 | Unknown Function |
| OG0000661 | 0.917 | 1.11E-08 | 0.334 | 0.001 | 4.099 | 3.143 | Function unknown (NAD(P)H-binding) |
| OG0009104 | 0.921 | 5.62E-08 | **0.0140** | **0.082** | 4.783 | 3.048 | Unknown Function |
| OG0010147 | 0.999 | 2.38E-17 | 0.471 | 0.001 | 4.565 | 2.857 | Function unknown (Zinc Finger) |
| OG0009580 | 0.999 | 2.50E-30 | 2.99 | 0.001 | 13.043 | 2.786 | Post-translational modification, protein turnover, and chaperones (Eukaryotic aspartyl protease) |
| OG0009038 | 0.999 | 2.16E-10 | 0.230 | 0.002 | 3.587 | 2.743 | Unknown Function |
| OG0009716 | 0.999 | 7.65E-20 | 0.817 | 0.001 | 3.261 | 2.694 | Unknown Function |
| OG0008134 | 0.961 | 1.59E-07 | **0.004** | **0.284** | 6.522 | 2.637 | Unknown Function (X-Pro dipeptidyl-peptidase C-terminal non-catalytic domain) |
| OG0010671 | 0.780 | 0.00094367 | 0.0324 | 0.02 | 1.304 | 2.400 | Unknown Function |
| OG0009627 | **0.161** | **0.211** | 0.039 | 0.009 | 2.446 | 2.357 | Unknown Function |

*Ratio of gene count in mycotroph to saprotroph *Trichoderma* genomes
**Ratio of gene count in endophytic to non-endophytic *Trichoderma* genomes

**Table S4. Mycoparasitism gene orthogroups of interest found in the highest mycotroph:saprotroph ratios as well as orthogroups only found in mycotrophs.**

| Mycoparasitism Gene orthogroup | Pagel's Lambda | Pagel's Lambda p-value | Blomberg's K | Blomberg's K p-value | M:S* | E:NE** | Eggnog Annotation |
|---|---|---|---|---|---|---|---|
| OG0008901 | 0.99 | 7.14E-24 | 1.029 | 0.001 | --*** | 1.978 | Amino acid transport and metabolism (Amino acid transport and metabolism) |
| OG0009237 | 0.99 | 1.23E-21 | 0.830 | 0.001 | -- | 1.714 | Unknown Function (LamB/YcsF family) |
| OG0009490 | 0.999 | 4.45E-28 | 1.997 | 0.001 | -- | 1.558 | Unknown Function (spectrin binding) |
| OG0010126 | 0.999 | 1.31E-23 | 0.93 | 0.001 | -- | 1.714 | Unknown Function |
| OG0010265 | 0.999 | 4.53E-41 | 8.061 | 0.001 | -- | 1.959 | Carbohydrate transport and metabolism (glycerone kinase activity) |
| OG0010647 | 0.999 | 3.14E-14 | 0.304 | 0.001 | -- | 2.000 | Unknown Function (Domain of unknown function DUF3328) |
| OG0011909 | 0.999 | 1.12E-24 | 1.617 | 0.001 | -- | 1.286 | No orthologs found |
| OG0011914 | 0.817 | 0.00596218 | 0.050 | 0.031 | -- | 0.686 | Unknown Function |
| OG0012676 | 0.999 | 5.02E-23 | 1.307 | 0.001 | -- | 0.429 | Unknown Function |
| OG0012734 | 0.999 | 5.02E-23 | 1.307 | 0.001 | -- | 0.429 | Nucleotide transport and metabolism, Translation, ribosomal structure and biogenesis (phospholipid biosynthetic process) |
| OG0013241 | 0.999 | 8.90E-15 | 0.410 | 0.001 | -- | 0.000 | Unknown Function (zinc finger) |
| OG0014884 | 0.999 | 2.87E-09 | 0.252 | 0.016 | -- | 0.000 | Unknown Function (Heterokaryon incompatibility protein (HET) |
| OG0016824 | **0.096** | **0.35427001** | **0.020** | **0.383** | -- | 0.000 | Post-translational modification, protein turnover, and chaperones (Belongs to the peptidase C1 family) |
| OG0009580 | 0.999 | 2.50E-30 | 2.99 | 0.001 | 13.043 | 2.786 | Post-translational modification, protein turnover, and chaperones (Eukaryotic aspartyl protease) |
| OG0009990 | 0.999 | 5.95E-22 | 0.848 | 0.001 | 10.434 | 1.929 | Unknown Function |

| | | | | | | | |
|---|---|---|---|---|---|---|---|
| OG0010098 | 0.999 | 8.71E-12 | 0.325 | 0.001 | 9.130 | 4.714 | Unknown Function |
| OG0002822 | 0.999 | 1.34E-19 | 0.68 | 0.001 | 7.826 | 2.000 | Unknown Function (Serine hydrolase) (FSH1) |
| OG0009146 | 0.999 | 7.54E-11 | 0.224 | 0.001 | 7.500 | 1.857 | Translation, ribosomal structure and biogenesis (amidase C869.01) |
| OG0008134 | 0.961 | 1.59E-07 | **0.004** | **0.284** | 6.521 | 2.63 | Unknown Function (X-Pro dipeptidyl-peptidase C-terminal non-catalytic domain) |
| OG0009419 | 0.999 | 9.95E-23 | 1.014 | 0.001 | 6.522 | 1.714 | Carbohydrate transport and metabolism, Cell wall/membrane/envelope biogenesis (NmrA-like family) |
| OG0008747 | 0.564 | 0.00033682 | 0.267 | 0.001 | 5.652 | 2.110 | Function unknown (TAP-like protein) |
| OG0008746 | 0.99 | 1.70E-14 | 0.307 | 0.001 | 5.435 | 1.486 | Unknown Function |
| OG0011291 | 0.747 | 2.51E-06 | 0.01 | 0.167 | 5.217 | 1.371 | Unknown Function |
| OG0011298 | 0.999 | 3.12E-05 | 0.12 | 0.002 | 5.217 | 2.143 | Unknown Function |

*Ratio of gene count in mycotrophic to saprotrophic *Trichoderma* genomes
**Ratio of gene count in endophytic to non-endophytic *Trichoderma* genomes
***Gene orthogroups that were not detected in saprotrophic genomes are designated with a "--" in place of a mycotroph:saprotroph ratio.

**Table S5. Placement of Brevicompactum clade ergot alkaloid BGC gene phylogenies**

| Gene | In clade of predominantly Hypocreales spp. | In clade of predominantly Xylariales spp. | Node support | Noteworthy results |
|---|---|---|---|---|
| Tribre1_123771 elymoclavine monooxygenase (*cloA*) | X | - | 100 | Brevicompactum clade is sister to other Hypocreales |
| Tribre1_123772 Chanoclavine synthase catalase (*easC*) | - | - | | Other *Trichoderma* in a clade with a homolog of tribre1 query gene |

| | | | | |
|---|---|---|---|---|
| Tribre1_123773 Lysergyl peptide synthetase subunit 2 (*lps2*) | - | - | | |
| Tribre1_123774 chanoclavine-I dehydrogenase (*easD)* | - | X | 96 | aspcor2, psevol1 within clade |
| Tribre1_123775 Chanoclavine-I aldehyde oxidoreductase (*easA*) | - | X | 100 | aspcor2, psevol1 within clade, Pleosporales in outgroup |
| Tribre1_123776 argoclavine dehydrogenase (*easG)* | - | X | 95 | aspcor2 sister to clade |
| Tribre1_123777 chanoclavine-I synthase oxidoreductase (*easE*) | - | X | 100 | aspcor2, psevol1 within clade |
| Dimethylallyltryptophan N-methyltransferase (*easF*) | - | X | 100 | aspcor2, psevol1 within clade |
| Tribre1_123778 Dimethylallyltryptophan synthase (*dmaW*) | - | X | 100 | aspcor2 within clade; bipsor1.1, cocsat1 (Pleosporales) in clade. Pleosporales have 100 support with Xylariales sp. (mictri1) |

| | | | | |
|---|---|---|---|---|
| Tribre1_123779 hypothetical protein | - | X | 100 | Absent from Clavicipitaceae, largely recovered in Xylariales, other Trichoderma in the sameclade, 1 copy/genome |
| Tribre1_123780 oxygenase (*easH*) | - | X | 100 | aspcor2, parnod1.1, epifes1.1 in clade |
| Tribre1_123781 Lysergyl peptide synthetase subunit 1 (*lps1*) | - | X | 52 | aspcor2, epifes1.1 in clade |

**Table S6**. **Functional annotations of the top 21 mycoparasitism genes contributing to the global PC1 and PC2 in the pPCA analysis.**

| Orthogroup | Functional Category | Description |
|---|---|---|
| OG0000020 | Secondary metabolites biosynthesis, transport, and catabolism | Enoyl-(Acyl carrier protein) reductase |
| OG0009047 | Function unknown | -- |
| OG0008954 | Function unknown | oligopeptide transmembrane transporter activity |
| OG0009301 | Function unknown | -- |
| OG0008688 | Translation, ribosomal structure and biogenesis | maturation of LSU-rRNA |
| OG0000599 | Function unknown | -- |
| OG0008901 | Amino acid transport and metabolism | glutamine biosynthetic process |
| OG0000008 | Secondary metabolites biosynthesis, transport, and catabolism | phosphopantetheine binding |
| OG0000642 | Post-translational modification, protein turnover, and chaperones | Belongs to the peptidase S1 family |
| OG0010265 | Carbohydrate transport and metabolism | glycerone kinase activity |
| OG0011331 | Defense mechanisms | ATPase activity |

| | | |
|---|---|---|
| OG0011314 | Function unknown | -- |
| OG0011345 | Function unknown | -- |
| OG0008929 | Function unknown | -- |
| OG0008903 | Inorganic ion transport and metabolism | alkaline phosphatase activity |
| OG0008978 | Function unknown | trichodiene synthase activity |
| OG0001105 | Inorganic ion transport and metabolism | O-methyltransferase activity |
| OG0001172 | Energy production and conversion | alpha-methylacyl-CoA racemase activity |
| OG0000193 | Defense mechanisms | ATPase activity |
| OG0000045 | Function unknown | -- |
| OG0000267 | Function unknown | -- |
